# Directed Chemical Evolution of Self-Assembling Artificial Proteins Utilizing a Supramolecular System

**DOI:** 10.64898/2025.12.10.693500

**Authors:** Ashwini R. Khyade, Britto S. Sandanaraj

## Abstract

Nature utilizes non-directed evolution to generate a vast array of biological macromolecules with remarkable diversity and functionality, though at a relatively slow pace. With advances in biotechnology, both directed and non-directed evolution approaches have been employed to design a wide range of biomacromolecules, including self-assembling artificial proteins (SAPs). However, these biological methods are limited to the natural set of twenty amino acids, resulting in SAPs that often lack functional diversity. Chemical approaches, such as micelle-assisted protein labeling technology (MAPLabTech), have enabled the creation of functional SAPs. Despite its potential, MAPLabTech remains a complex, low-throughput, and time-consuming methodology. To overcome these challenges, herein, we disclose a new method, termed **S**upramolecule-**A**ssisted **P**rotein **Lab**eling **Tech**nology **(SAPLabTech)**. In this method, γ-cyclodextrin acts as a host to solubilize hydrophobic chemical probes, forming a cyclodextrin–probe supramolecular complex that enables site-specific bioconjugation reaction and the synthesis of well-defined monodisperse SAPs in quantitative yield. The versatility of this technology is demonstrated by employing three distinct classes of chemical probes with varying warhead functionalities, linker lengths, and tail hydrophobicity to construct diverse SAP libraries with rich structural features.

## Introduction

Self-assembling artificial proteins (SAPs) have emerged as powerful scaffolds for advanced applications.^1–8^ Computational protein design, pioneered by Baker and others, has proven to be a highly effective method for constructing SAPs from first principles.^3,8–10^ Although these computationally designed SAPs differ substantially from natural proteins in both structure and function. They are typically synthesized in a bacterial system employing standard recombinant DNA technology and thus composed exclusively of the twenty naturally occurring amino acids.^9,11,12^ In contrast, proteins in higher organisms undergo extensive post-translational modifications - hundreds of distinct chemical alterations that greatly expand the functional diversity of the proteome.^13–16^ These modifications endow natural proteins with a vast range of chemical functionalities essential for complex biological processes.^17–19^

Unlike genetic methods, chemical approaches enable the creation of SAPs from an unrestricted palette of chemical building blocks, offering far greater structural and functional diversity. However, before 2018, most studies were limited to a few proteins, and, most importantly, SAPs designed through chemical methods lacked rigorous analytical and biophysical characterization. ^20–24^ To address these challenges, our group developed micelle-assisted protein labeling technology (MAPLabTech) - a versatile chemical strategy for designing and synthesizing well-defined, monodisperse SAPs.^25^Over the past several years, we have applied MAPLabTech to generate hundreds of distinct SAPs and established scalable purification methods for their synthesis. In addition, we have carried out comprehensive analytical and biophysical studies of SAPs.^25–32^ These studies have provided deep mechanistic insight into the self-assembly behavior of SAPs, enabling precise control over their assembly^26–33^ and disassembly dynamics.^25^ We have demonstrated that MAPLabTech has ability to fine-tune critical parameters of protein nanoassemblies such as hydrodynamic size, molecular weight, oligomeric state, and surface charge with an extraordinary precision.^26^ In addition, this technology was employed for the design of stimuli-responsive protein nanomaterials ^25,27, 28^ and artificial protein nanoreactors.^31^

Despite its maturity, reliability, and applicability, MAPLabTech has certain limitations, including (i) slow reaction kinetics, (ii) incomplete conversion, and (iii) labor-intensive purification.^25–32^ Therefore, the development of new chemical technologies with improved attributes is imperative. Towards that goal, herein, we disclose an alternative chemical strategy - **supramolecule-assisted protein labeling technology (SAPLabTech)**. This method enables the rapid synthesis of libraries of well-defined, monodisperse SAPs with enhanced structural diversity (Fig. 1).

**Figure 1:**
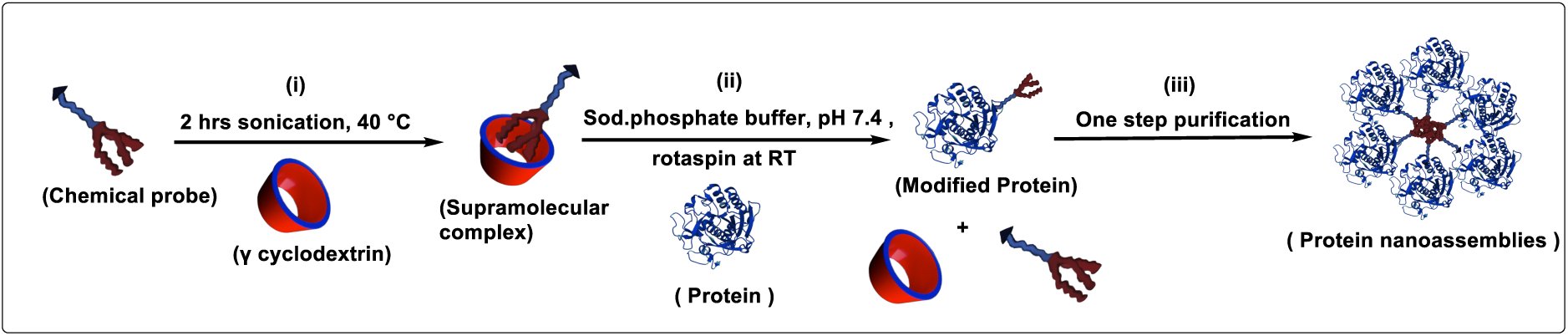
Cartoon representation of Supramolecule-assisted Protein Labeling Technology (SAPLabTech). **(i)** A hydrophobic probe is solubilized in aqueous solution via supramolecular complex formation using sonication and heating at 40 °C for 2 h. **(ii)** The complex is incubated with the protein at room temperature under to achieve modification. **(iii)** protein nano-assemblies is purified through a single step.

## Results

### Encapsulation of Hydrophobic Chemical Probe

Synthesis of SAPs involves site-specific labeling of a soluble globular hydrophilic protein by a hydrophobic chemical probe to generate facially amphiphilic proteins. However, labeling of proteins in a 100% aqueous medium is not possible due to the extreme hydrophobicity of chemical probes. To circumvent this issue, we hypothesized that well-defined monomeric water-soluble monodisperse supramolecular scaffolds (host molecules) with a larger hydrophobic interior could be used for the solubilization of hydrophobic chemical probes. The ideal host for the protein-bioconjugation reaction should have the following attributes: (i) high aqueous solubility in water, (ii) ability to solubilize a wide range of chemical probes, (iii) minimal non-specific protein binding, and (iv) most importantly, the warhead functionality of the probe should be exposed to bulk solvent. Considering these stringent requirements, cyclodextrins (CDs) were chosen as the host molecules because they naturally fulfill all the criteria, offering biocompatibility, renewable origin, and well-established regulatory approval.

Structurally, CDs are cyclic oligosaccharides composed of α-1,4-linked glucopyranose units that form a hydrophobic internal cavity capable of encapsulating small organic molecules under mild aqueous conditions.^33–35^ Their cavity dimensions (4.7 Å - α-CD, 6.0 Å - β-CD, and 8.0 Å - γ-CD) allow selective accommodation of diverse guest molecules. In addition to their high solubility, biodegradability, and GRAS (Generally Recognized as Safe) status, CDs can be chemically modified to improve further solubility and complex stability.^36–40^ These tunable properties often surpass those of other macrocyclic hosts, which generally suffer from lower solubility.^41–45^ Initially, we tested the ability of all the cyclodextrins (α, β, and γ) to solubilize the hydrophobic chemical probe. The results from absoption studies shown that both α- and γ-cyclodextrins are capable of solubilizing the probe (MI-TEG-C12-3T, Fig 2a). In parallel, we have carried NMR studies, the presence of aromatic protons indicates formation of supramolecular complex (Fig. 2b). We also found that γ-cyclodextrin has exhibited better solubilizing ability than α-cyclodextrin and hence was used for further studies

**Figure 2.**
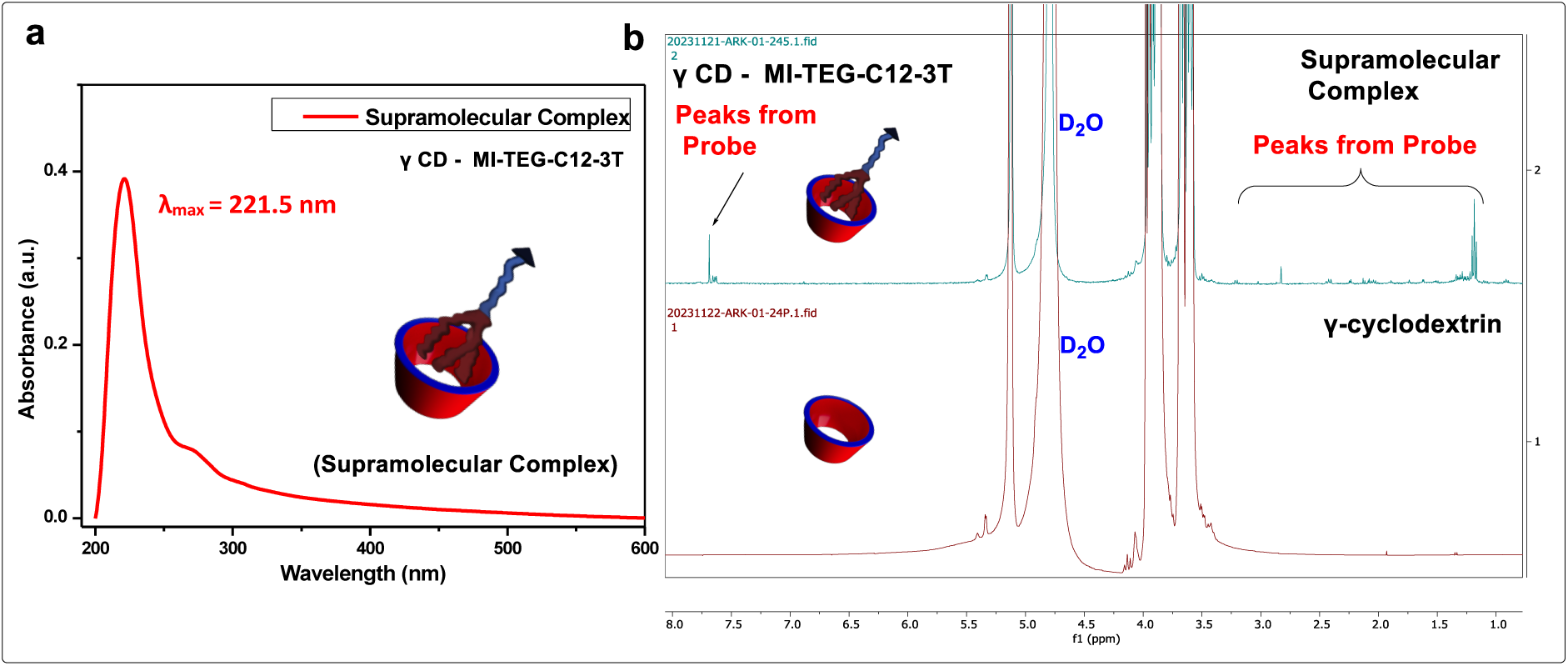
Evidence for the Formation of the Supramolecular Complex. **a.** UV–visible spectroscopy study of γ-cyclodextrin:MI-TEG-C12-3T supramolecular complex. **b.** ^1^H NMR study of γ-cyclodextrin:MI-TEG-C12-3T supramolecular complex.

### Molecular Design of Chemical Probes

Taking inspiration from our previous study,^25–32^ we designed a small library of hydrophobic chemical probes composed of three key structural elements: A bioconjugation tag for site-selective protein modification, a linear spacer for conformational flexibility, and a hydrophobic tail that promotes self-assembly of the resulting facially amphiphilic proteins. Each library is designed uniquely to target a set of proteins, taking advantage of their unique chemical reactivity. Using this design framework, we successfully designed and synthesized three sets of probe families containing maleimide (MI), fluorophosphonate (FP), and N-pyridine carboxaldehyde (NPC) as reactive functional groups (Fig.3b,4b,5b).

The MI probes consist of a maleimide moiety (for cysteine targeting), an oligoethylene glycol linker, and a hydrophobic alkyl chain. These probes enable the selective modification of cysteine residues, while leaving other nucleophilic sites, such as lysine amines or serine hydroxyl functionalities, unaffected.^32^This chemo-selectivity makes the maleimide approach especially valuable for preparing structurally defined bioconjugates.^32^The FP probes are designed for active-site-directed labeling of serine proteases. This class of probes reacts rapidly and site-specifically with the catalytic serine residue, enabling covalent attachment.^25–30^ The NPC probes were specifically engineered for site-selective bioconjugation of the *N*-terminal residue of globular proteins.^31,46^ The probe libraries’ diversity is expanded by changing linker length (TEG *vs* OEG), hydrophobic tags (C12 *vs* C18), and varying architectures (1T *vs* 3T). All probes were synthesized through a multi-step organic synthesis, followed by purification and rigorous characterization using standard analytical techniques. The detailed synthetic procedure is provided in the supporting information.

### Chemical Evolution of SAPs from Cysteine-Containing Proteins

Initially, we wanted to test our methodology on a simple system and therefore chose bovine serum albumin (BSA) as our target protein, as this protein is widely used in a variety of bioconjugation chemistries.^47–49^ BSA exhibits excellent solubility, a well-defined globular structure, and a single surface-exposed cysteine residue.^50,51^ Initially, we prepared a supramolecular complex of γ-CD with MI probe **1**, as described in the experimental method section, followed by a bioconjugation reaction with BSA at 25 °C for overnight (Fig. 3a). MALDI-TOF was used to monitor the bioconjugation reaction. To our delight, we observed a new peak at 67,567 Da corresponding to bioconjugate BSA-MI-TEG-C12-3T, in addition to the native BSA peak at 66,616 Da. To test whether this methodology applies to other globular proteins containing a single cysteine residue, we selected human serum albumin (HSA) as a test case. As expected, the reaction yielded a bioconjugate (HSA-MI-TEG-C12-3T) with a molecular weight of 67,644 Da, demonstrating that the method applies to proteins containing a single surface-exposed cysteine residue.

**Figure 3.**
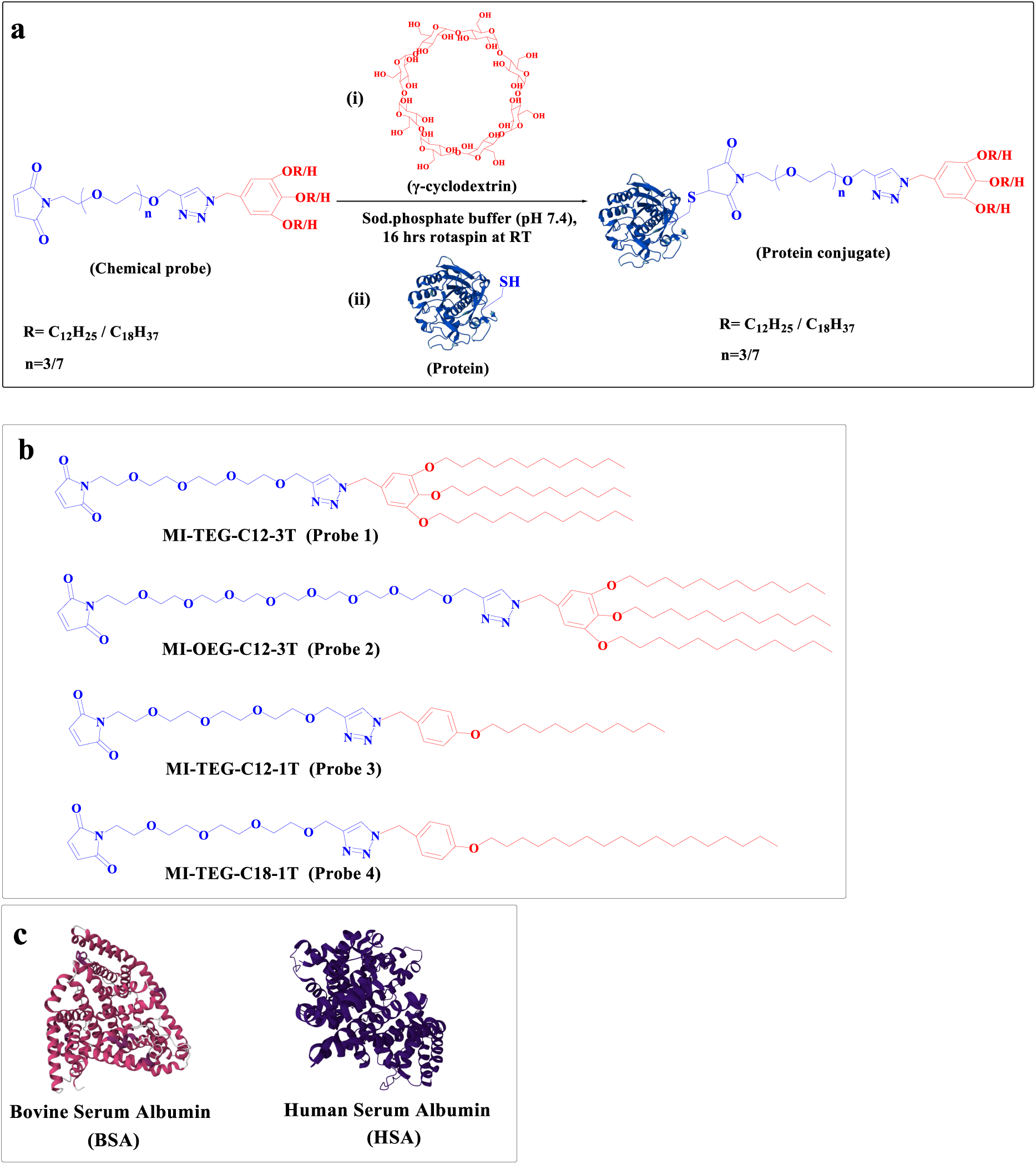
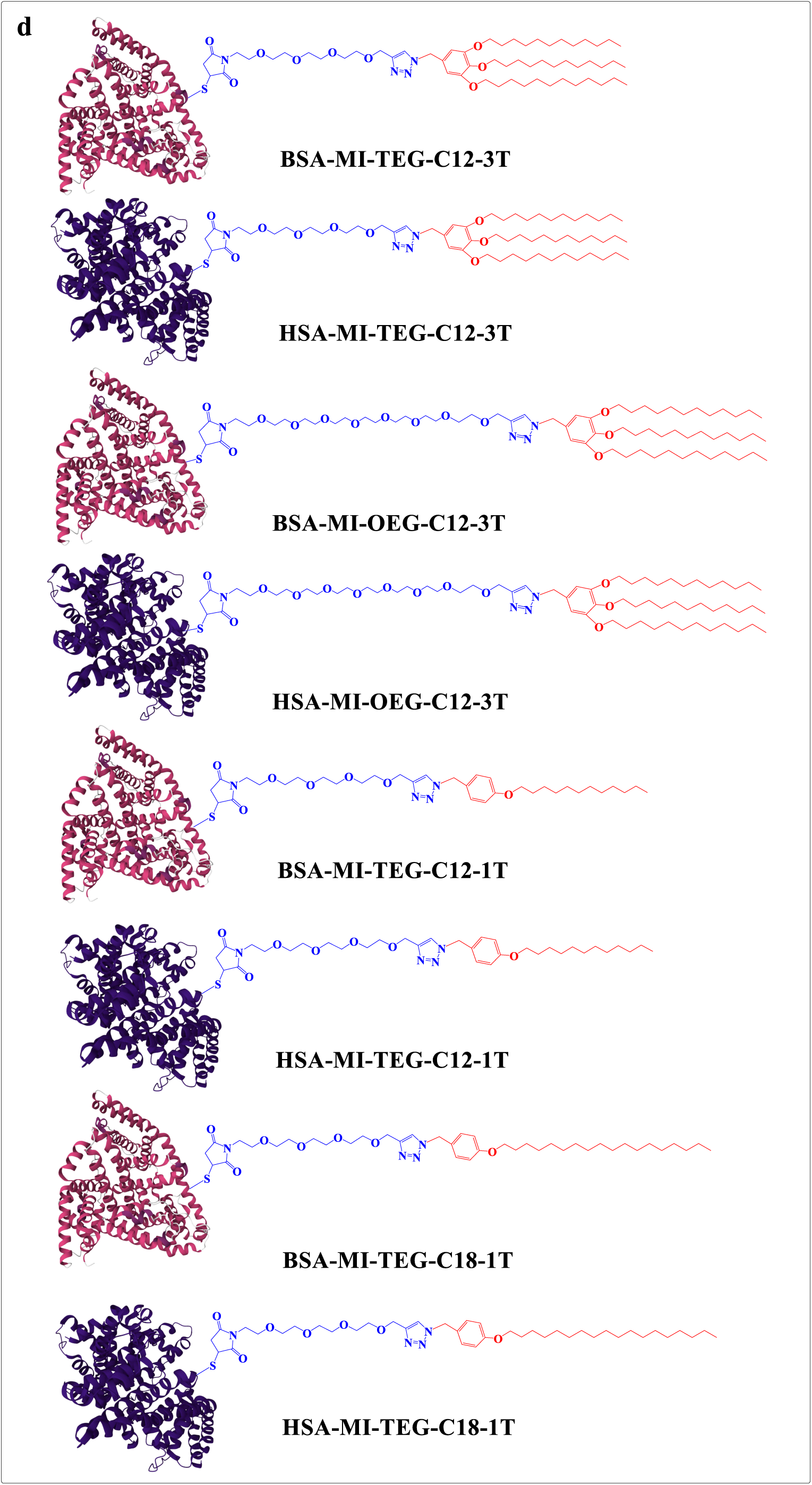
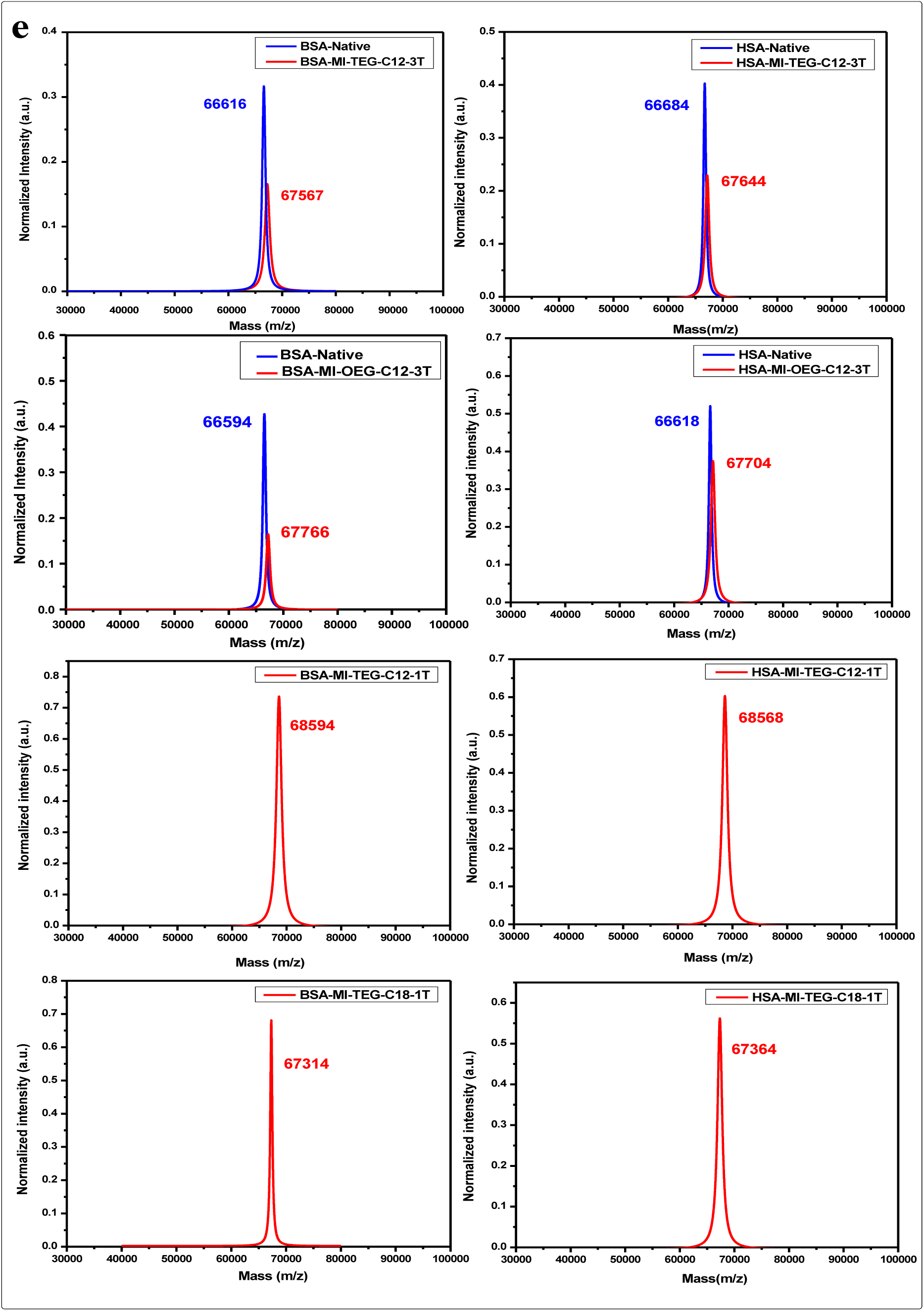
**a.** Schematic representation of the general bioconjugation strategy for maleimide (MI) probes using Supramolecule-Assisted Protein Labeling Technology **(SAPLabTech). b.** Chemical structure of the hydrophobic maleimide probe. **C.** Representative crystal structures of target proteins used for bioconjugation. **d.** A library of protein conjugates **e.** MALDI-TOF-MS results of the protein conjugate library

To assess the role of linker length and increase probe library diversity, we synthesized a new probe **2** (MI-OEG-C12-3T), having an octaethylene glycol (OEG) spacer. As expected, this probe also reacted with both BSA and HSA proteins, as evident from MALDI-TOF results (Fig.3e). In addition, we also synthesized MI probes **3** and **4** having different hydrophobic tail lengths (C12-1T vs C18-1T). We were surprised to observe that the most hydrophobic probe **4**, could be solubilized using γ-cyclodextrin, and both supramolecular complexes were able to react with BSA and HSA proteins (Fig. 3e). To our delight, we observed quantitative conversion for both the probes. Although useful, MI-based probes can only be used for a very limited set of globular proteins containing a single surface-exposed cysteine residue; hence, the impact of this method is very limited. Therefore, we set out to test whether this technology could be used for a chemical probe with broad coverage of protein reactivity.

### Chemical evolution of SAPs from Serine Proteases

FP probes have been previously used to target the entire class of serine hydrolases, which comprises approximately 200 proteins in humans alone, and thus prove to be a broad chemical probe for targeting a wide range of proteins.^25,52,53^ Motivated by these facts, we employed SAPLabTech for the labeling of serine proteases. Initially, we synthesized the fluorophosphonate probe FP-TEG-C12-3T, which is then solubilized using γ-cyclodextrin to form a supramolecular complex (Fig. 4a). As a test case, we first attempted bioconjugation of proteinase K (serine protease) utilizing SAPLabTech, and the reaction progress was monitored using MALDI-TOF. To our delight, we observed a single peak at 29,901 Da for the expected product, with no peak for the native protein, indicating a quantitative conversion. Since we observed quantitative conversion within 18 hours, we were curious to understand the kinetics of this bioconjugation reaction. Therefore, we carried out reactions at different time points and monitored reaction progress using MALDI-TOF. To our delight, we observed a 60% conversion within one hour and quantitative conversion in 2 hours, indicating heightened reactivity of the FP probes (Fig. 4e).

**Figure 4.**
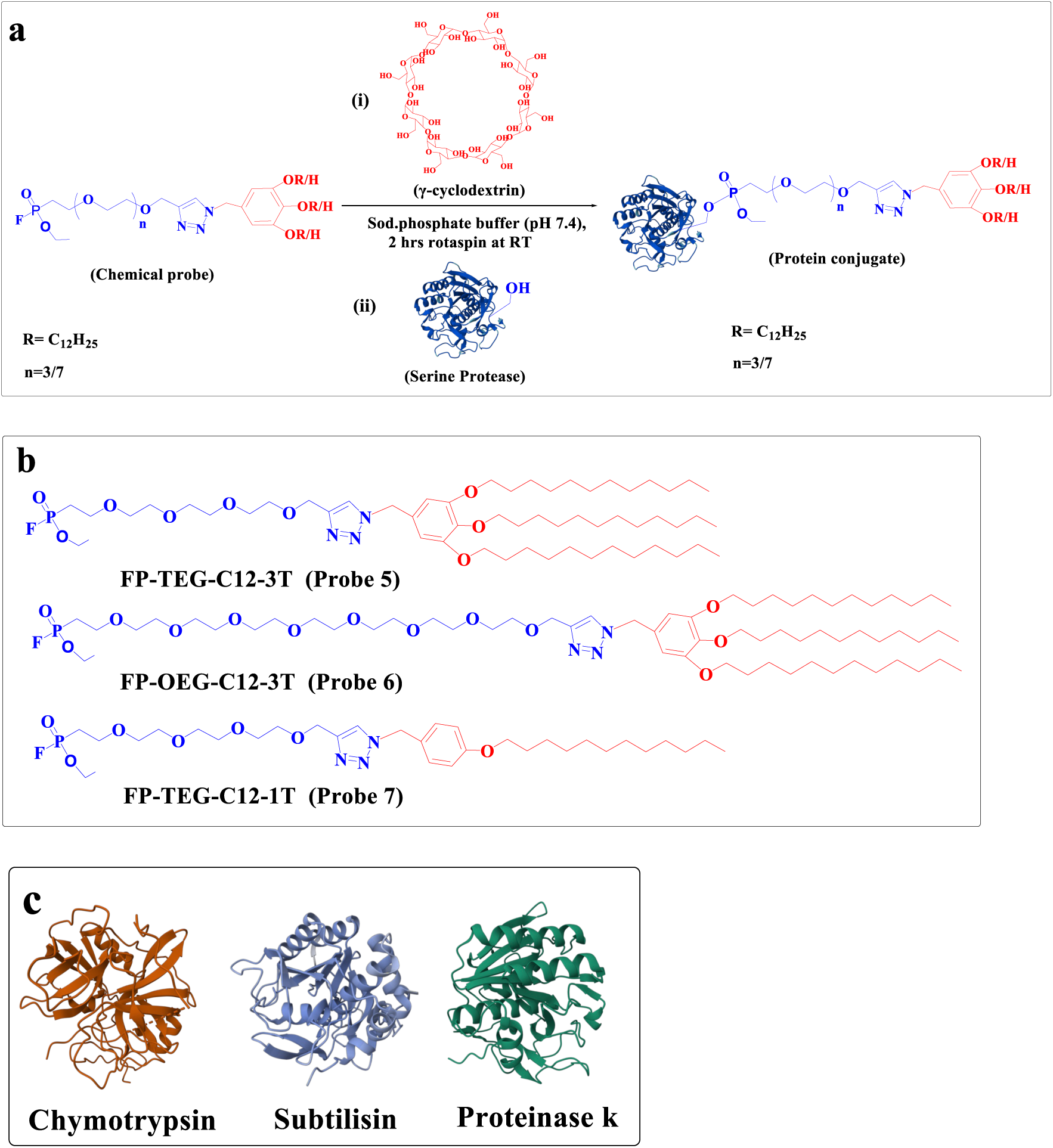
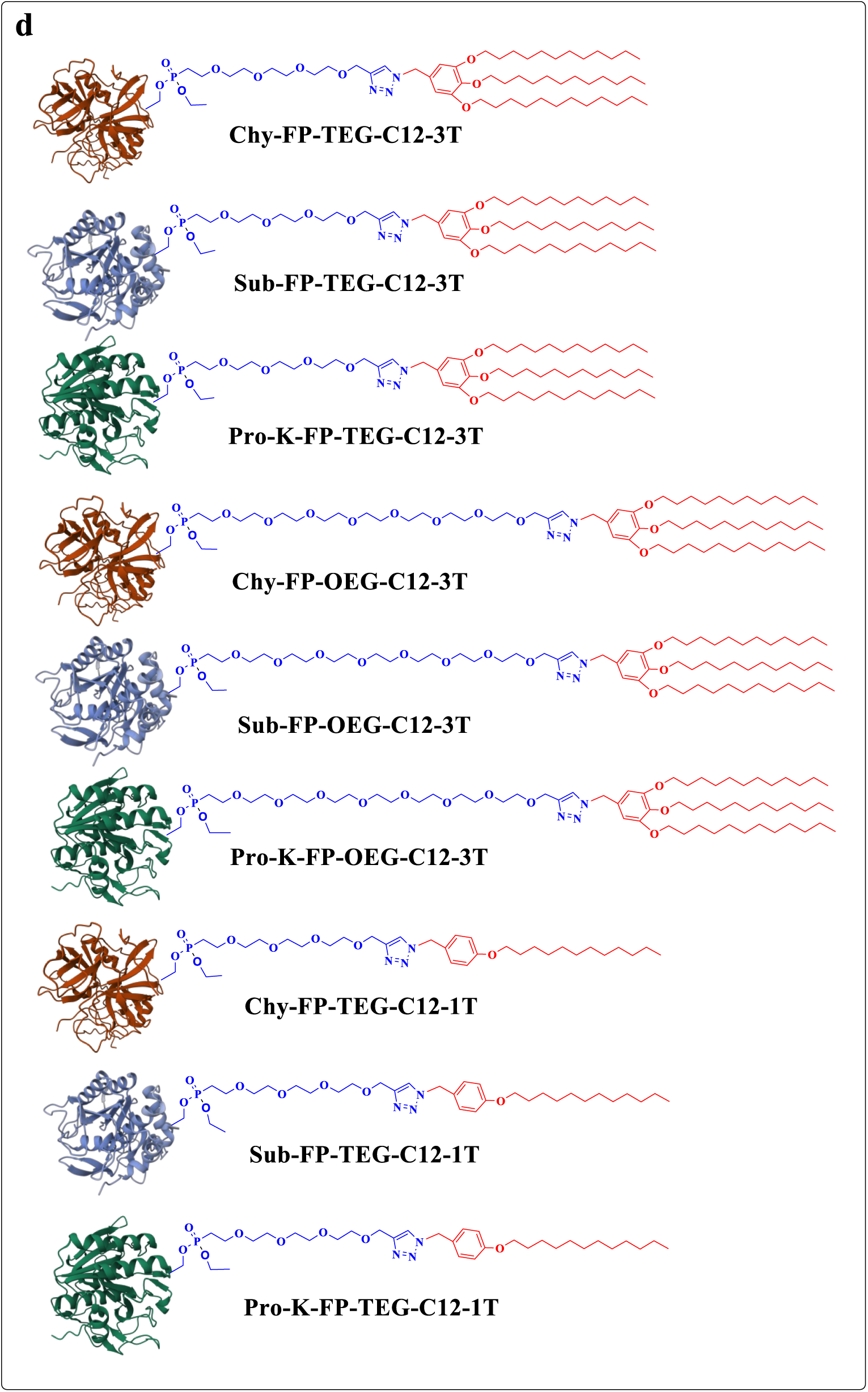
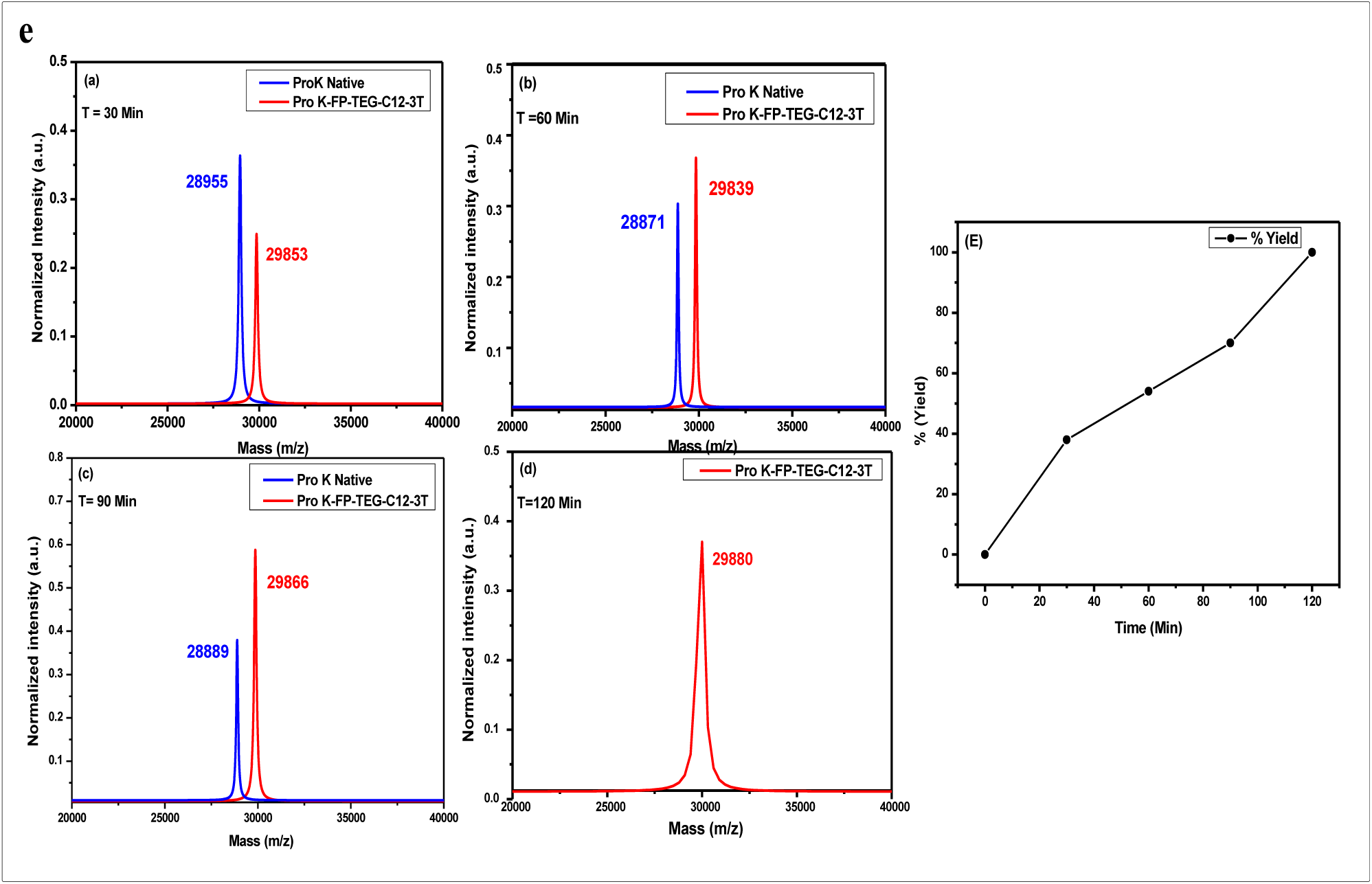
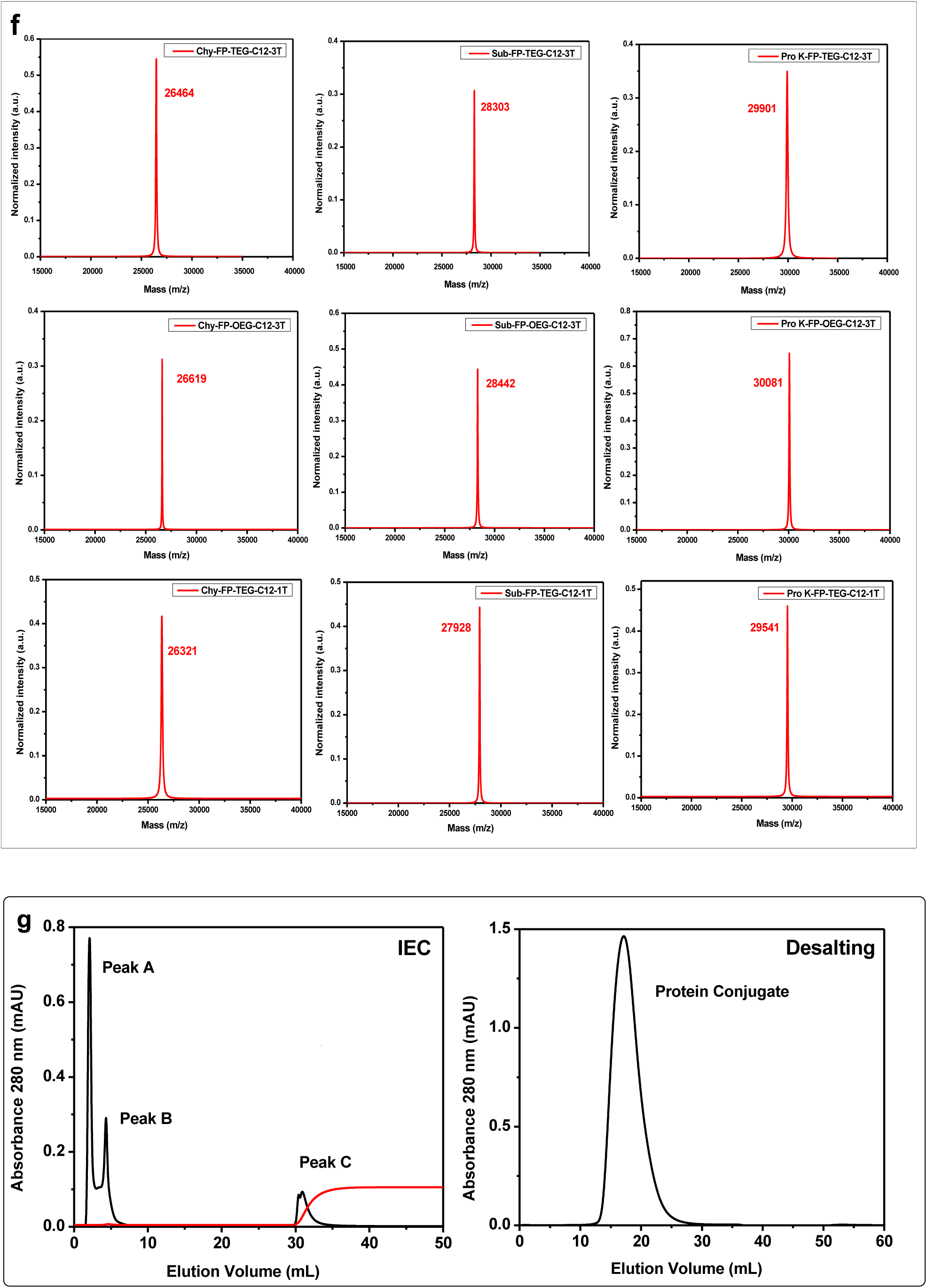
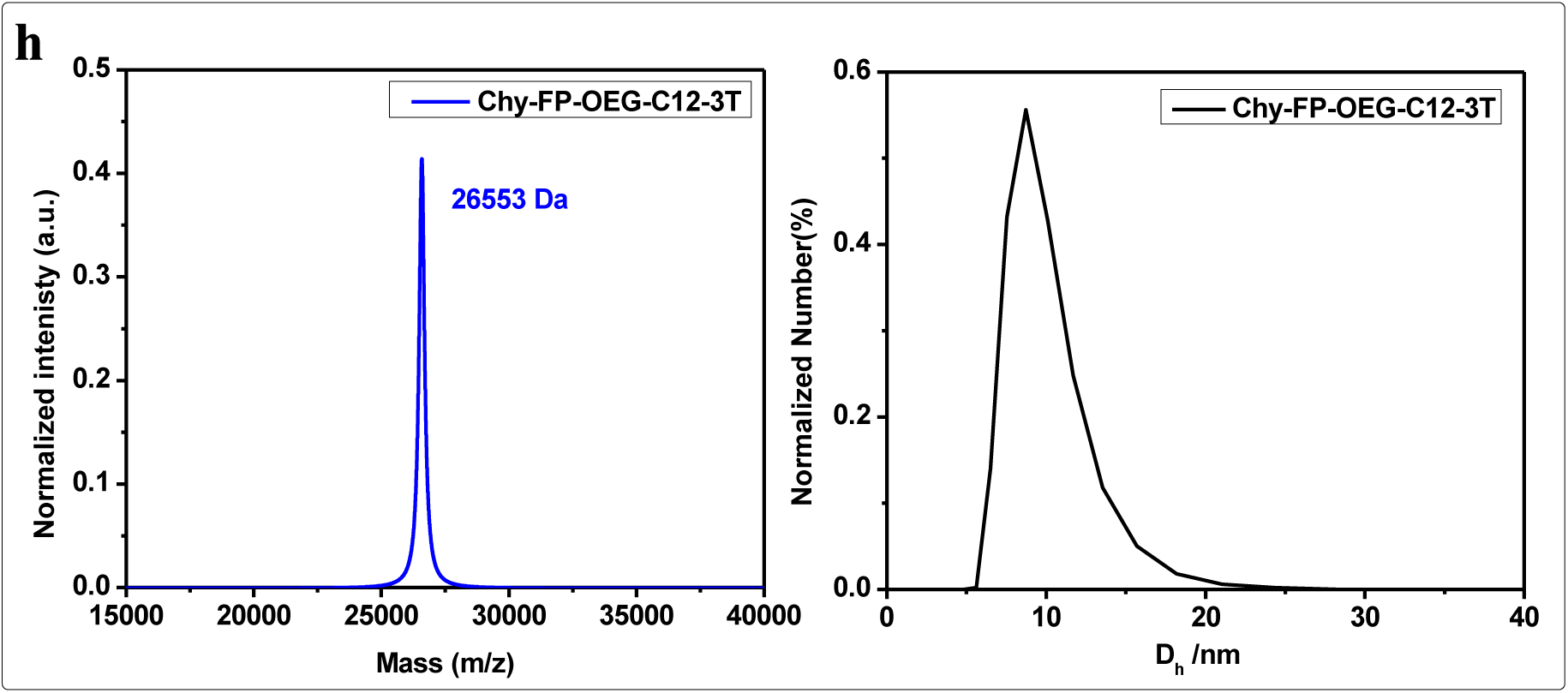
**a.** Schematic illustration of the general bioconjugation strategy for fluorophosphonate (FP) probes using SAPLabTech**. b.** Chemical structure of the hydrophobic fluorophosphonate probes. **C.** Representative crystal structures of target proteins used for bioconjugation. **d.** A library of protein conjugates **e.** Time-dependent bioconjugation reaction. **f.** MALDI-TOF results of a library of protein conjugates **g.** Purification of Chy-FP-OEG-C12-3T using IEC, followed by desalting. **h.** The MALDI-TOF and DLS results of the Chy-FP-OEG-C12-3T conjugate.

To evaluate the generality of the method, the same protocol was applied to other serine proteases, including chymotrypsin and subtilisin. Complete conversion was achieved within 2 hours, with new peaks observed at 26,464 Da and 28,303 Da, corresponding to Chy–FP-TEG-C12-3T and Sub-FP-TEG-C12-3T, respectively (Fig. 4f). These results demonstrate that the SAPLabTech is broadly applicable across different serine proteases, irrespective of their size or charge. The method was further validated using the FP-OEG-C12-3T probe, which features an longer and more flexible OEG spacer. Quantitative bioconjugation was achieved within 2 hours for proteinase K, chymotrypsin, and subtilisin, yielding single MALDI-TOF peaks at 30,081 Da, 26,619 Da, and 28,442 Da, respectively. These results confirm that variations in spacer length do not affect the reaction efficiency. Subsequent bioconjugation experiments with chemical probes with different hydrophobic tags (FP-TEG-C12-1T and FP-TEG-C18-1T probes), revealed distinct trends in reactivity. The FP-TEG-C12-1T probe yielded complete conversion for all three proteases, as evident from the MALDI-ToF results (Figures a, b, and c).

To evaluate the scalability of SAPLabTech, we next scaled up the synthesis of Chy–FP–OEG–C12–3T bioconjugate (50 mg scale). Purification of the crude reaction mixture was achieved through an ion-exchange chromatography (Fig.4g), as detailed in the experimental section. The overall isolated yield of the purified bioconjugate was 50%, representing a substantial improvement over the MAPLabTech approach,^25^ which yielded only about 20% with an additional purification step.^26–33^ Dynamic light scattering (DLS) studies further demonstrated that the conjugate self-assembled in an aqueous solution, forming monodisperse protein nanoparticles with a diameter of 10 nm (Fig. 4h). These results match with the DLS study of Chy–FP–OEG–C12–3T made through MAPLabTech.^25^ Although the utilization of FP probes advanced the applicability of SAPLabTech from a handful of proteins to hundreds of proteins, it is still limited considering the vast number of proteins that are present in the nature. Therefore, we set out to look for a chemical probe that can virtually react with any globular protein.

### Universal Chemical Method for Chemical Evolution of SAPs

NPC probes have been previously shown to react with a wide range of proteins, irrespective of their class, structure, and function.^54^ We have also previously employed MAPLabTech, along with the NPC chemical probe, to create functional SAPs.^31^ The supramolecular complex was formed as described in the previous section, followed by a bioconjugation reaction with BSA (Fig. 5a). MALDI-TOF results revealed a distinct new peak at 67,933 Da, corresponding to the BSA-NPC-TEG-C12-3T conjugate, confirming successful labelling of the *N*-terminal residue. Encouraged by these results, we extended the methodology to HSA under identical conditions. MALDI-TOF results revealed the appearance of a new conjugate peak at 67,747 Da, corresponding to HSA-NPC-TEG-C12-3T, alongside the native HSA peak (Fig. 5e). To further assess the generality of this methodology, chymotrypsin, a serine protease with a different size and surface charge distribution^31^, was also subjected to bioconjugation with the NPC-TEG-C12-3T probe. As expected, we observed a new peak at 26,616 Da, corresponding to Chy-NPC-TEG-C12-3T, confirming that the SAPLabTech facilitates bioconjugation irrespective of protein class, charge, or size. Subsequent experiments were performed using NPC-TEG-C12-1T and NPC-TEG-C18-1T probes, which feature distinct hydrophobic tails, to evaluate the influence of tail hydrophobicity on conjugation efficiency. The NPC-TEG-C12-1T probe yielded quantitative conversion for both BSA and HSA, as evidenced by the presence of sharp and well-defined peaks at 67,424 Da (BSA–NPC-TEG-C12-1T) and 67,434 Da (HSA–NPC-TEG-C12-1T), respectively.

**Figure 5.**
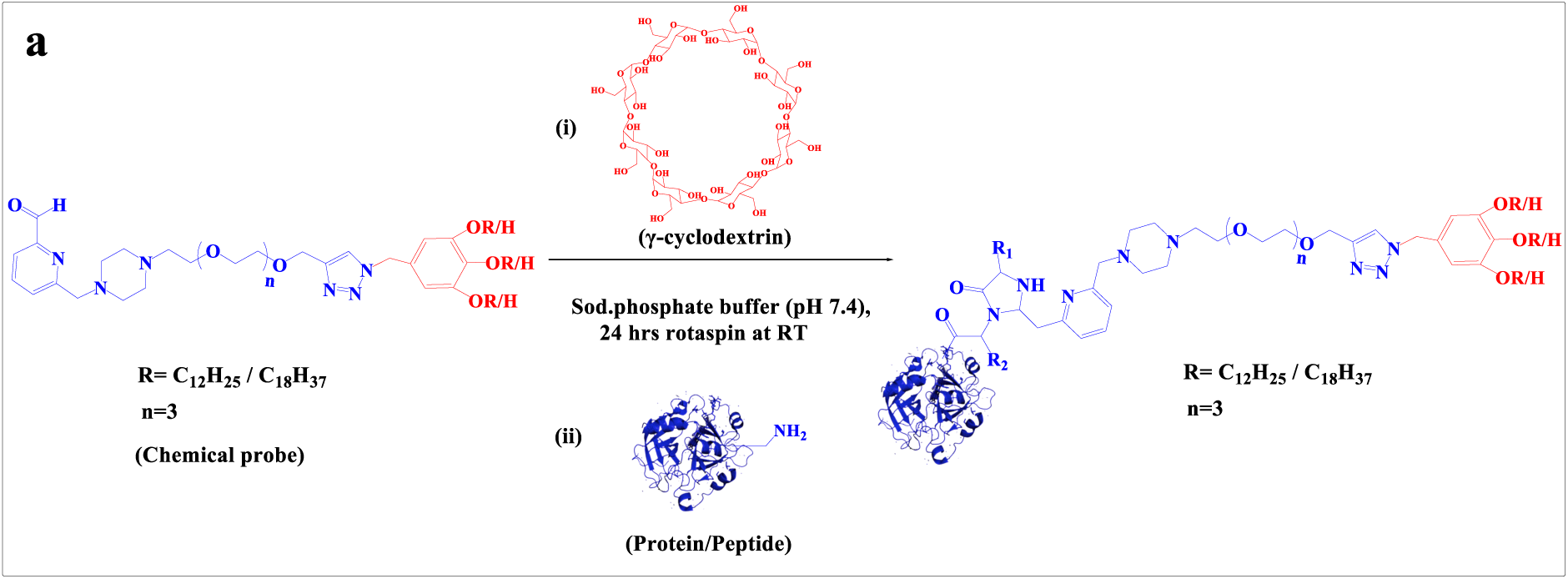
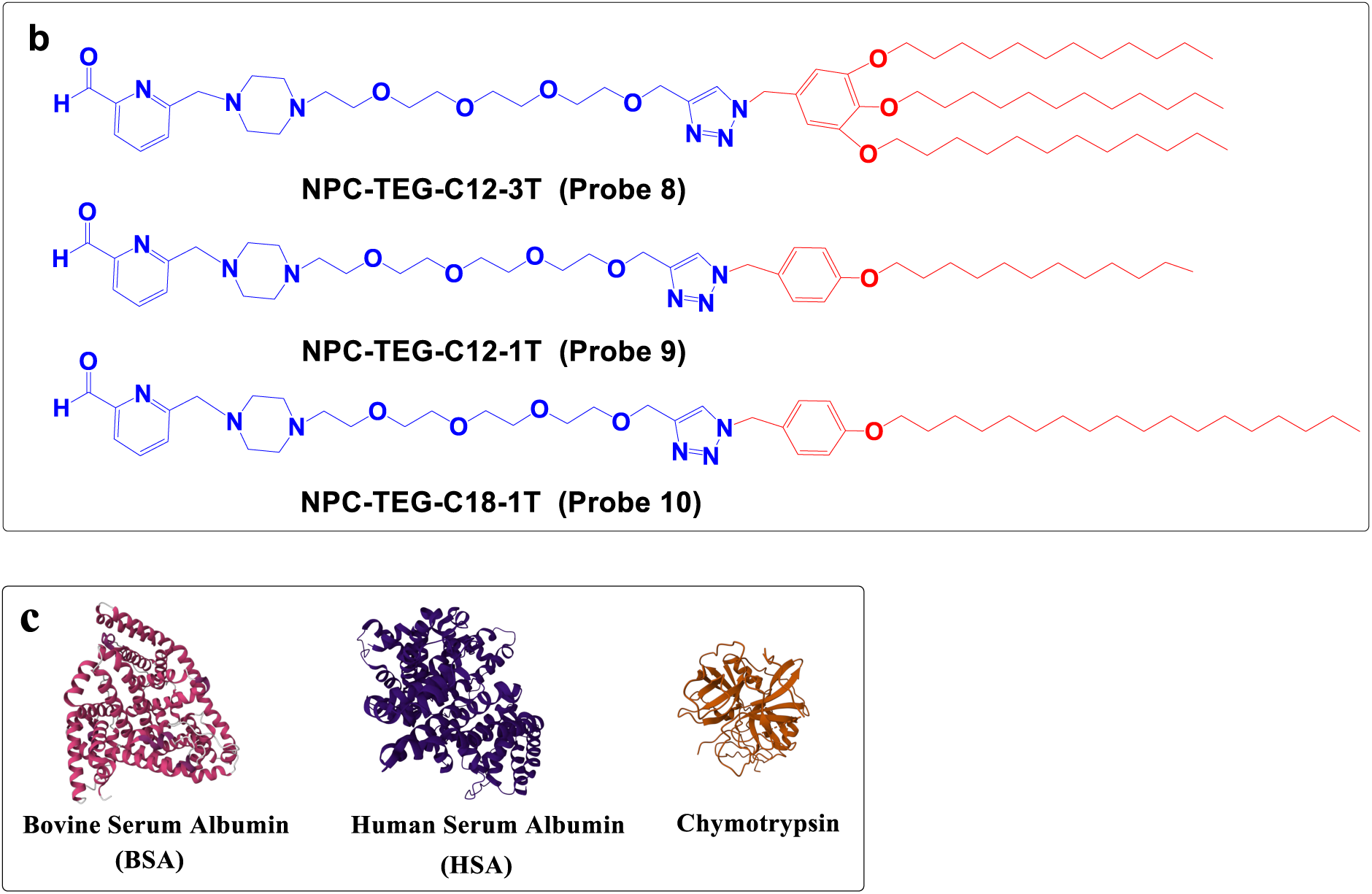
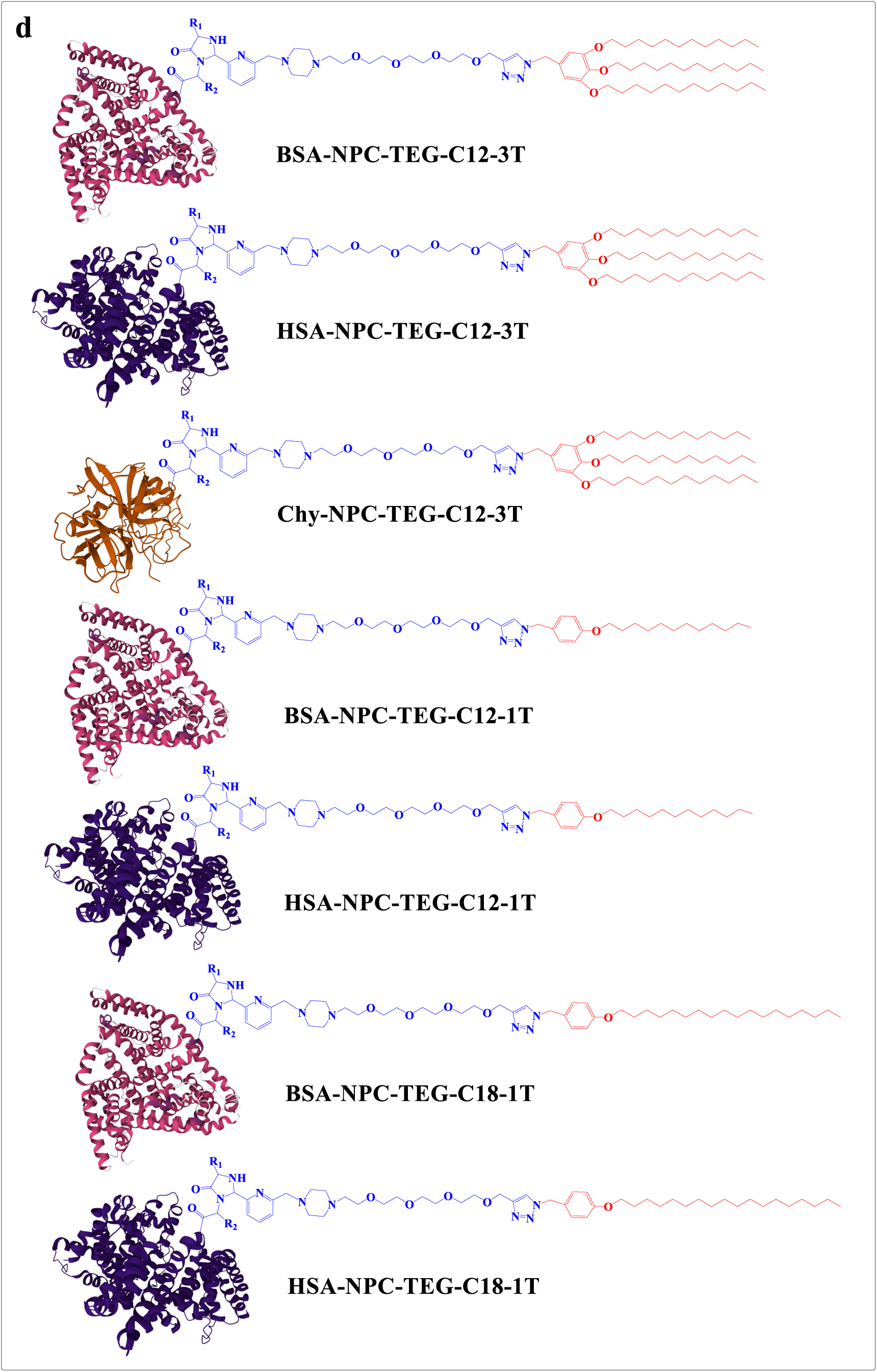
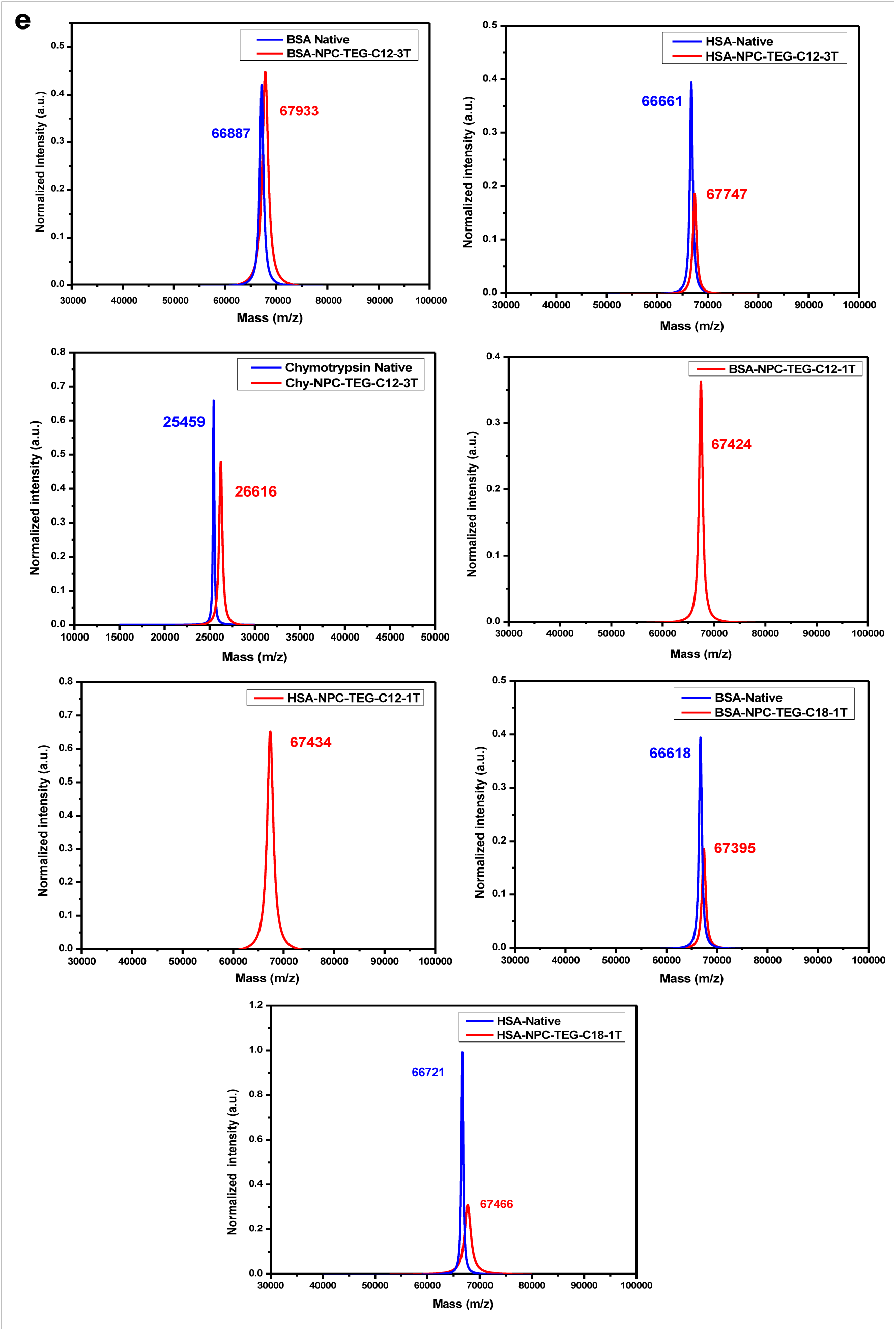
**a.** Schematic representation of the general bioconjugation strategy for N-pyridine carboxaldehyde (NPC) probes using **(SAPLabTech). b.** Chemical structure of the NPC probe. **C.** Representative crystal structures of target proteins used for bioconjugation. **d.** A library of protein conjugates **e.** MALDI-TOF results of the protein conjugate library.

Similarly, successful conjugation was achieved with the NPC-TEG-C18-1T probe, with new peaks observed at 67,395 Da for BSA-NPC-TEG-C18-1 T and 67,466 Da for HSA-NPC-TEG-C18-1T, clearly distinct from the native protein peaks (Fig. 5e). The above result is significant because the yield of the bioconjugation reaction with water-soluble, hydrophilic NPC probes is typically around 60-70%.^54^ Collectively, these findings validate the broad applicability and robustness of SAPLabTech across diverse protein classes and reactive functionalities.

## Discussion

The remarkable diversity of naturally occurring biomacromolecules (nucleic acids, proteins, and carbohydrates) is a result of natural evolution. Directed evolution, based on genetic methods, has emerged as a robust technology for designing biomacromolecules with specialized functions with an extraordinary precision.^55–57^ Although extremely powerful, genetic methods operate on limited building blocks (4 in the case of nucleic acid and 20 in the case of proteins), and their potential is somewhat limited.^9,11,12^ Previously, SAPs have been synthesized using directed genetic evolution; however, most of them lack function.^20–24^

Our group invented a chemical method called MAPLabTech, which provides opportunities to chemically evolve SAPs containing building blocks other than the standard 20 amino acids. This method provides an opportunity to incorporate stimulus-sensitive functionalities, such as photo-sensitive,^25,28^ and redox-sensitive groups onto SAPs.^30^Although extremely powerful, this method has certain limitations, such as slow reaction rates, incomplete conversions, and difficult purification. The success of any evolution-based methods highly depends on the throughput and scalability, and MAPLabTech lacked both. Therefore, the present study is focused on inventing an alternative technology to direct the chemical evolution of SAPs. SAPLabTech marks a significant leap forward in the evolution of self-assembling artificial proteins (SAPs). By using γ-cyclodextrin to form supramolecular complexes with hydrophobic probes, SAPLabTech effectively enhances the solubility of hydrophobic chemical probes in 100% aqueous conditions, creating a simple, fast, and reliable platform for the expedited synthesis of SAP with excellent diversity and functionality.

Our results demonstrate that γ-cyclodextrin (γ-CD), through its hydrophobic cavity and hydrophilic exterior, serves as an efficient supramolecular host that dramatically enhances the solubility of hydrophobic chemical probes of different classes. The formation of γ-CD–probe supramolecular complexes provides homogeneous reaction conditions, thereby eliminating the semi-heterogeneous conditions posed by MAPLabTech.^25–32^

Using MI probes, we first demonstrated the site-selective modification of cysteine-containing proteins such as BSA and HSA. Moreover, the compatibility of this method with probes of varied spacer lengths (TEG vs. OEG) and hydrophobic tail architectures (C12 vs C18, 1T vs 3T) highlights its adaptability for diverse probe designs. Encouraged by these results, we extended SAPLabTech to the synthesis of SAPs from serine proteases using FP probes. Traditionally, MAPLabTech-mediated bioconjugations required up to 18 hours to achieve a conversion of ∼60–70% due to semi-heterogeneous reaction conditions and slower diffusion of the micellar system.^25^ In stark contrast, SAPLabTech achieved near-quantitative conversions within just two hours. Time-dependent MALDI-TOF MS analyses confirmed that over 60% of bioconjugation occurred within one hour - underscoring the remarkable rate enhancement. Furthermore, the consistency of results across proteins of varying functions (enzymes, fatty acid carriers, etc.), molecular weights, and isoelectric points establishes the generality of this platform.

To further expand the scope and demonstrate the universality of SAPLabTech, we employed NPC probes to achieve N-terminal bioconjugation. The NPC-TEG-C12-3T probe reacted efficiently with BSA, HSA, and chymotrypsin, yielding SAPs within 24 hours. A similar efficiency was shown when varying the linker length or hydrophobic tail (C12 *vs.* C18, 1T *vs*. 3T). The reproducible high-yield labeling across structurally and functionally diverse proteins clearly demonstrates that this method provides a truly universal chemical bioconjugation approach. Notably, SAPLabTech achieved an overall yield of approximately 50% using a simplified one-step purification process, whereas the MAPLabTech method yielded only about 20% after three-step purification. This improvement highlights the advantages of this method in minimizing purification losses and enhancing conjugation efficiency. Furthermore, DLS results reveal that the Chy–FP–OEG–C12–3T self-assembles into a monodisperse protein nanoparticle with a size of 10 nm.

## Conclusion

In this work, we introduce SAPLabTech as a powerful, versatile, and broadly applicable strategy for creating self-assembling artificial proteins (SAPs). By leveraging the solubilizing capability of γ-cyclodextrin, this approach enhances the aqueous dispersibility of hydrophobic probes, thereby enabling rapid, site-specific, and high-yield labeling of diverse protein classes, including those bearing reactive cysteine residues, active serine residues, and free N-terminal amino acid-containing proteins. SAPLabTech effectively overcomes the key limitations of previous methodologies, offering shorter reaction times, quantitative conversions, and streamlined single purification, all while preserving the protein’s inherent ability to self-assemble into uniform, well-defined nanostructures. Overall, SAPLabTech represents a simple, scalable, and efficient chemical platform for the design and synthesis of precisely engineered protein nanoassemblies. By bridging the principles of synthetic protein chemistry and supramolecular host–guest interactions, this approach establishes a robust foundation for the next generation of functional protein-based materials, with promising implications for targeted therapeutics, diagnostic systems, and the design of advanced biomaterials.

## Experimental Methods

### Materials

All reagents and chemicals were obtained from reputable commercial suppliers and used as received without further purification. γ-Cyclodextrin (γ-CD) was purchased commercially and employed directly in all experiments.

### Nuclear Magnetic Resonance (NMR) Spectroscopy

^1^H and ^13^C NMR spectra were recorded on a Bruker 400 MHz spectrometer. Proton chemical shifts (δ) are reported in ppm relative to residual solvent peaks in CDCl- (δ 7.26) and D-O (δ 4.79). Carbon chemical shifts were referenced to CDCl- (δ 77.16). High-purity solvents (CDCl- and D-O) were purchased from Sigma-Aldrich.

### Protein Selection

For thiol–maleimide chemistry, bovine serum albumin (BSA, 66.6 kDa) and human serum albumin (HSA, 66.6 kDa) were selected as model cysteine-containing proteins. For fluorophosphonate chemistry, serine proteases including chymotrypsin (25.4 kDa), subtilisin (27.3 kDa), and proteinase K (28.9 kDa) were chosen. N-terminal modification experiments were performed using BSA, HSA, and chymotrypsin. All proteins were sourced from Sigma-Aldrich, and their molecular weights were verified prior to modification.

### Matrix Preparation and Molecular Weight Determination

MALDI-TOF MS (Shimadzu MALDI-8020) was used for the precise determination of molecular weights of native and modified proteins. Proteins <40 kDa: 2′,6′-Dihydroxyacetophenone (DHAP, 3.5 mg) and diammonium hydrogen citrate (DAHC, 4.5 mg) were dissolved separately in 150 µL ethanol and 200 µL Milli-Q water, followed by 1 min sonication and vortexing. The DAHC solution (50 µL) was added to DHAP, mixed, to get final matrix and combined with protein samples in a 1:1:1 ratio with 2% trifluoroacetic acid (TFA). Samples were applied to MALDI plates and air-dried for 30–40 min prior to analysis (mass range 15,000–40,000 Da). Protein concentration was maintained at 50 µM.

Proteins >40 kDa: Sinapinic acid (15 mg) was dissolved in 1 mL of a 70:30 water: acetonitrile solution containing 0.1% TFA. Protein samples were mixed with matrix in a 1:5 ratio, transferred to MALDI plates (2 µL), and air-dried for 30 min. Mass scanning was performed over 40,000–80,000 Da with protein concentration at 50 µM.

### Protein Modification and Monitoring

Inclusion complexes were prepared by mixing 40–60 equivalents of the chemical probe with 20–40 equivalents of γ-CD in 500 µL of 50 mM sodium phosphate buffer (pH 7.4), followed by sonication at 40 °C for 2 h. A small amount of organic co-solvent (1% THF) was included to facilitate IC formation. The resulting host–guest complex was then combined with the protein solution (100 µM), adjusting the final reaction volume to 1 mL and the final protein concentration to 50 µM. The reactions were incubated at room temperature with gentle rotation (20 rpm) on a rotary mixer.

### Sample Preparation for MALDI-TOF MS

Proteins <40 kDa: 3 µL of reaction mixture was combined with 3 µL of 2% TFA and 3 µL of matrix, vortexed, and applied to the MALDI plate.

Proteins >40 kDa: 3 µL of reaction mixture was mixed with 15 µL of matrix, vortexed, and applied to the MALDI plate.

MALDI-TOF MS analysis allowed accurate monitoring of protein modification and confirmation of conjugate formation. Reaction times were adjusted according to protein type and probe chemistry.

### Purification of modified protein

The purification of modified proteins was performed using an AKTA Pure chromatography system (Cytiva, USA). The purification process involved ion-exchange chromatography (IEC) followed by desalting/buffer exchange. Cation-exchange chromatography was carried out using a HiScreen SP FF column (Cytiva). The column was pre-equilibrated with 50 mM sodium acetate buffer (pH 5.0) prior to sample loading. After sample injection, post-injection equilibration was continued until the complete removal of protein conjugate (Peak A), unreacted probe, and γ-cyclodextrin (Peak B). The impurity from native protein, along with protein conjugate (Peak C), was subsequently eluted using 50 mM sodium phosphate buffer (pH 7.4) containing 1 M NaCl as the elution buffer. A detailed analysis of each peak using MALDI-TOF mass spectrometry is provided in the supporting information.

Following ion-exchange purification, desalting/buffer exchange were performed using a HiPrep 26/10 Desalting column (Cytiva) to transfer the purified protein conjugate into Milli-Q water for further analysis and storage.

### MALDI-TOF MS for purified protein conjugate

The lyophilized protein conjugate was analysed using MALDI-TOF MS.

Protein solution A-A lyophilized protein conjugate weighing 1 mg was dissolved in sodium phosphate buffer at a pH of 7.4.

Triton-X 100 solution-2 µL of Triton X-100 in 1 mL of sodium phosphate buffer at pH 7.4.

Sample for MALDI-TOF MS – 4 µL of Triton X-100 solution was mixed with 96 µL of protein solution A. The analysis was performed using the same matrix mentioned above, depending on the protein molecular weight.

### Dynamic Light Scattering (DLS) studies

The hydrodynamic diameter of the protein complexes was determined using a Zetasizer Nano 2590 instrument (Malvern Instruments, UK). Protein samples were prepared at a concentration of 200µM in 50 mM sodium phosphate buffer (pH 7.4). The 1 mL filtered sample was transferred into disposable polystyrene cuvettes, and the mean particle size was measured at a scattering angle of 90° under standard conditions.

## ASSOCIATED CONTENT

### Supplementary Information

### Author Information

**Notes**: We have filed Indian and US patent for the technology disclosed in this paper.

## ACKNOWLEDGEMENTS

We would like to thank IISER-Pune and Molbio Diagnostic Pvt Ltd for funding.

## References

(1) Huang, P.-S.; Boyken, S. E.; Baker, D. The Coming of Age of de Novo Protein Design. Nature 2016, 537 (7620), 320–327. 10.1038/nature19946.

(2) Yeates, T. O. Geometric Principles for Designing Highly Symmetric Self-Assembling Protein Nanomaterials. Annu Rev Biophys 2017, 46 (1), 23–42. 10.1146/annurev-biophys-070816-033928.

(3) King, N. P.; Sheffler, W.; Sawaya, M. R.; Vollmar, B. S.; Sumida, J. P.; André, I.; Gonen, T.; Yeates, T. O.; Baker, D. Computational Design of Self-Assembling Protein Nanomaterials with Atomic Level Accuracy. Science (1979) 2012, 336 (6085), 1171–1174. 10.1126/science.1219364.

(4) Salgado, E. N.; Radford, R. J.; Tezcan, F. A. Metal-Directed Protein Self-Assembly. Acc Chem Res 2010, 43 (5), 661–672. 10.1021/ar900273t.

(5) Luo, Q.; Hou, C.; Bai, Y.; Wang, R.; Liu, J. Protein Assembly: Versatile Approaches to Construct Highly Ordered Nanostructures. Chem Rev 2016, 116 (22), 13571–13632. 10.1021/acs.chemrev.6b00228.

(6) Ma, Y.; Nolte, R. J. M.; Cornelissen, J. J. L. M. Virus-Based Nanocarriers for Drug Delivery. Adv Drug Deliv Rev 2012, 64 (9), 811–825. 10.1016/j.addr.2012.01.005.

(7) Ljubetič, A.; Lapenta, F.; Gradišar, H.; Drobnak, I.; Aupič, J.; Strmšek, Ž.; Lainšček, D.; Hafner-Bratkovič, I.; Majerle, A.; Krivec, N.; Benčina, M.; Pisanski, T.; Veličković, T. Ć.; Round, A.; Carazo, J. M.; Melero, R.; Jerala, R. Design of Coiled-Coil Protein-Origami Cages That Self-Assemble in Vitro and in Vivo. Nat Biotechnol 2017, 35 (11), 1094–1101. 10.1038/nbt.3994.

(8) Hsia, Y.; Bale, J. B.; Gonen, S.; Shi, D.; Sheffler, W.; Fong, K. K.; Nattermann, U.; Xu, C.; Huang, P.-S.; Ravichandran, R.; Yi, S.; Davis, T. N.; Gonen, T.; King, N. P.; Baker, D. Design of a Hyperstable 60-Subunit Protein Icosahedron. Nature 2016, 535 (7610), 136–139. 10.1038/nature18010.

(9) Dou, J.; Vorobieva, A. A.; Sheffler, W.; Doyle, L. A.; Park, H.; Bick, M. J.; Mao, B.; Foight, G. W.; Lee, M. Y.; Gagnon, L. A.; Carter, L.; Sankaran, B.; Ovchinnikov, S.; Marcos, E.; Huang, P.-S.; Vaughan, J. C.; Stoddard, B. L.; Baker, D. De Novo Design of a Fluorescence-Activating β-Barrel. Nature 2018, 561 (7724), 485–491. 10.1038/s41586-018-0509-0.

(10) Tinberg, C. E.; Khare, S. D.; Dou, J.; Doyle, L.; Nelson, J. W.; Schena, A.; Jankowski, W.; Kalodimos, C. G.; Johnsson, K.; Stoddard, B. L.; Baker, D. Computational Design of Ligand-Binding Proteins with High Affinity and Selectivity. Nature 2013, 501 (7466), 212–216. 10.1038/nature12443.

(11) Rocklin, G. J.; Chidyausiku, T. M.; Goreshnik, I.; Ford, A.; Houliston, S.; Lemak, A.; Carter, L.; Ravichandran, R.; Mulligan, V. K.; Chevalier, A.; Arrowsmith, C. H.; Baker, D. Global Analysis of Protein Folding Using Massively Parallel Design, Synthesis, and Testing. Science (1979) 2017, 357 (6347), 168–175. 10.1126/science.aan0693.

(12) Silva, D.-A.; Yu, S.; Ulge, U. Y.; Spangler, J. B.; Jude, K. M.; Labão-Almeida, C.; Ali, L. R.; Quijano-Rubio, A.; Ruterbusch, M.; Leung, I.; Biary, T.; Crowley, S. J.; Marcos, E.; Walkey, C. D.; Weitzner, B. D.; Pardo-Avila, F.; Castellanos, J.; Carter, L.; Stewart, L.; Riddell, S. R.; Pepper, M.; Bernardes, G. J. L.; Dougan, M.; Garcia, K. C.; Baker, D. De Novo Design of Potent and Selective Mimics of IL-2 and IL-15. Nature 2019, 565 (7738), 186–191. 10.1038/s41586-018-0830-7.

(13) Walsh, C. T.; Garneau-Tsodikova, S.; Gatto, G. J. Protein Posttranslational Modifications: The Chemistry of Proteome Diversifications. Angewandte Chemie International Edition 2005, 44 (45), 7342–7372. 10.1002/anie.200501023.

(14) Khoury, G. A.; Baliban, R. C.; Floudas, C. A. Proteome-Wide Post-Translational Modification Statistics: Frequency Analysis and Curation of the Swiss-Prot Database. Sci Rep 2011, 1 (1), 90. 10.1038/srep00090.

(15) Nørregaard Jensen, O. Modification-Specific Proteomics: Characterization of Post-Translational Modifications by Mass Spectrometry. Curr Opin Chem Biol 2004, 8 (1), 33–41. 10.1016/j.cbpa.2003.12.009.

(16) Seo, J.-W.; Lee, K.-J. Post-Translational Modifications and Their Biological Functions: Proteomic Analysis and Systematic Approaches. BMB Rep 2004, 37 (1), 35–44. 10.5483/BMBRep.2004.37.1.035.

(17) Mann, M.; Jensen, O. N. Proteomic Analysis of Post-Translational Modifications. Nat Biotechnol 2003, 21 (3), 255–261. 10.1038/nbt0303-255.

(18) Karve, T. M.; Cheema, A. K. Small Changes Huge Impact: The Role of Protein Posttranslational Modifications in Cellular Homeostasis and Disease. J Amino Acids 2011, 2011, 1–13. 10.4061/2011/207691.

(19) Deribe, Y. L.; Pawson, T.; Dikic, I. Post-Translational Modifications in Signal Integration. Nat Struct Mol Biol 2010, 17 (6), 666–672. 10.1038/nsmb.1842.

(20) Chin, J. W. Expanding and Reprogramming the Genetic Code of Cells and Animals. Annu Rev Biochem 2014, 83 (1), 379–408. 10.1146/annurev-biochem-060713-035737.

(21) Velonia, K.; Rowan, A. E.; Nolte, R. J. M. Lipase Polystyrene Giant Amphiphiles. J Am Chem Soc 2002, 124 (16), 4224–4225. 10.1021/ja017809b.

(22) Gauba, V.; Hartgerink, J. D. Self-Assembled Heterotrimeric Collagen Triple Helices Directed through Electrostatic Interactions. J Am Chem Soc 2007, 129 (9), 2683–2690. 10.1021/ja0683640.

(23) Wong, L. S.; Khan, F.; Micklefield, J. Selective Covalent Protein Immobilization: Strategies and Applications. Chem Rev 2009, 109 (9), 4025–4053. 10.1021/cr8004668.

(24) Chari, R. V. J.; Miller, M. L.; Widdison, W. C. Antibody–Drug Conjugates: An Emerging Concept in Cancer Therapy. Angewandte Chemie International Edition 2014, 53 (15), 3796–3827. 10.1002/anie.201307628.

(25) Sandanaraj, B. S.; Reddy, M. M.; Bhandari, P. J.; Kumar, S.; Aswal, V. K. Rational Design of Supramolecular Dynamic Protein Assemblies by Using a Micelle-Assisted Activity-Based Protein-Labeling Technology. Chemistry – A European Journal 2018, 24 (60), 16085–16096. 10.1002/chem.201802824.

(26) Sandanaraj, B. S.; Reddy, M. M.; Rao, K. J.; Bhandari, P. J. Rational Design of Semi-Synthetic Protein Complexes with the Defined Oligomeric State. ChemistrySelect 2019, 4 (20), 6397–6402. 10.1002/slct.201901317.

(27) Sandanaraj, B. S.; Bhandari, P. J.; Reddy, M. M.; Lohote, A. B.; Sahoo, B. Design, Synthesis, and Self-Assembly Studies of a Suite of Monodisperse, Facially Amphiphilic, Protein–Dendron Conjugates. ChemBioChem 2020, 21 (3), 408–416. 10.1002/cbic.201900341.

(28) Bhandari, P. J.; Sandanaraj, B. S. Programmed and Sequential Disassembly of Multi-responsive Supramolecular Protein Nanoassemblies: A Detailed Mechanistic Investigation. ChemBioChem 2021, 22 (5), 876–887. 10.1002/cbic.202000581.

(29) Bhandari, P. J.; Reddy, M. M.; Rao, K. J.; Sandanaraj, B. S. Rapid Chemical Synthesis of Self-Assembling Semi-Synthetic Proteins. J Org Chem 2021, 86 (13), 8576–8589. 10.1021/acs.joc.1c00195.

(30) Bhandari, P. J.; Sandanaraj, B. S. Rational Design of Programmable Monodisperse Semi-Synthetic Protein Nanomaterials Containing Engineered Disulfide Functionality**. ChemBioChem 2021, 22 (20), 2966–2972. 10.1002/cbic.202100288.

(31) Reddy, M. M.; Bathla, P.; Sandanaraj, B. S. A Universal Chemical Method for Rational Design of Protein-Based Nanoreactors**. ChemBioChem 2021, 22 (21), 3042–3048. 10.1002/cbic.202100315.

(32) Reddy, M. M.; Bhandari, P.; Hati, K. C.; Sandanaraj, B. S. Rational Design of Self-Assembling Artificial Proteins Utilizing a Micelle-Assisted Protein Labeling Technology (MAPLabTech): Testing the Scope**. ChemBioChem 2022, 23 (7). 10.1002/cbic.202100607.

(33) Kurkov, S. V.; Loftsson, T. Cyclodextrins. Int J Pharm 2013, 453 (1), 167–180. 10.1016/j.ijpharm.2012.06.055.

(34) Szejtli, J. Introduction and General Overview of Cyclodextrin Chemistry. Chem Rev 1998, 98 (5), 1743–1754. 10.1021/cr970022c.

(35) Rekharsky, M. V.; Inoue, Y. Complexation Thermodynamics of Cyclodextrins. Chem Rev 1998, 98 (5), 1875–1918. 10.1021/cr970015o.

(36) Uekama, K.; Hirayama, F.; Irie, T. Cyclodextrin Drug Carrier Systems. Chem Rev 1998, 98, (5) 2045–2076. 10.1021/cr970025p.

(37) Saenger, W. Cyclodextrin Inclusion Compounds in Research and Industry. Angewandte Chemie International Edition in English 1980, 19 (5), 344–362. 10.1002/anie.198003441.

(38) Loftsson, T.; Duchene, D. Cyclodextrins and Their Pharmaceutical Applications. Int J Pharm 2007, 329 (1–2), 1–11. 10.1016/j.ijpharm.2006.10.044.

(39) Jambhekar, S. S.; Breen, P. Cyclodextrins in Pharmaceutical Formulations I: Structure and Physicochemical Properties, Formation of Complexes, and Types of Complex. Drug Discov Today 2016, 21 (2), 356–362. 10.1016/j.drudis.2015.11.017.

(40) Hedges, A. R. Industrial Applications of Cyclodextrins. Chem Rev 1998, 98 (5), 2035–2044. 10.1021/cr970014w.

(41) Geng, W.-C.; Jiang, Z.-T.; Chen, S.-L.; Guo, D.-S. Supramolecular Interaction in the Action of Drug Delivery Systems. Chem Sci 2024, 15 (21), 7811–7823. 10.1039/D3SC04585D.

(42) Torchilin, V. P. Structure and Design of Polymeric Surfactant-Based Drug Delivery Systems. Journal of Controlled Release 2001, 73 (2–3), 137–172. 10.1016/S0168-3659(01)00299-1.

(43) Pouton, C. W. Formulation of Poorly Water-Soluble Drugs for Oral Administration: Physicochemical and Physiological Issues and the Lipid Formulation Classification System. European Journal of Pharmaceutical Sciences 2006, 29 (3–4), 278–287. 10.1016/j.ejps.2006.04.016.

(44) Loftsson, T.; Brewster, M. E. Pharmaceutical Applications of Cyclodextrins: Basic Science and Product Development. Journal of Pharmacy and Pharmacology 2010, 62 (11), 1607–1621. 10.1111/j.2042-7158.2010.01030.x.

(45) Savjani, K. T.; Gajjar, A. K.; Savjani, J. K. Drug Solubility: Importance and Enhancement Techniques. ISRN Pharm 2012, 2012, 1–10. 10.5402/2012/195727.

(46) Gilmore, J. M.; Scheck, R. A.; Esse-Kahn, A. P.; Joshi, N. S.; Francis, M. B. N-Terminal Protein Modification through a Biomimetic Transamination Reaction. Angewandte Chemie International Edition 2006, 45 (32), 5307–5311. 10.1002/anie.200600368.

(47) Narla, S. N.; Pinnamaneni, P.; Nie, H.; Li, Y.; Sun, X.-L. BSA–Boronic Acid Conjugate as Lectin Mimetics. Biochem Biophys Res Commun 2014, 443 (2), 562–567. 10.1016/j.bbrc.2013.12.006.

(48) Rohiwal, S. S.; Ellederova, Z.; Tiwari, A. P.; Alqarni, M.; Elazab, S. T.; El-Saber Batiha, G.; Pawar, S. H.; Thorat, N. D. Self-Assembly of Bovine Serum Albumin (BSA)–Dextran Bio-Nanoconjugate: Structural, Antioxidant and *in Vitro* Wound Healing Studies. RSC Adv 2021, 11 (8), 4308–4317. 10.1039/D0RA09301G.

(49) Li, X.; Li, T.; Guo, J.; Wei, Y.; Jing, X.; Chen, X.; Huang, Y. PEGylation of Bovine Serum Albumin Using Click Chemistry for the Application as Drug Carriers. Biotechnol Prog 2012, 28 (3), 856–861. 10.1002/btpr.1526.

(50) Mehtala, J. G.; Kulczar, C.; Lavan, M.; Knipp, G.; Wei, A. Cys34-PEGylated Human Serum Albumin for Drug Binding and Delivery. Bioconjug Chem 2015, 26 (5), 941–949. 10.1021/acs.bioconjchem.5b00143.

(51) Koniev, O.; Wagner, A. Developments and Recent Advancements in the Field of Endogenous Amino Acid Selective Bond Forming Reactions for Bioconjugation. Chem. Soc. Rev. 2015, 44 (15), 5495–5551. 10.1039/C5CS00048C.

(52) Wang, C.; Abegg, D.; Dwyer, B. G.; Adibekian, A. Discovery and Evaluation of New Activity-Based Probes for Serine Hydrolases. ChemBioChem 2019, 20 (17), 2212–2216. 10.1002/cbic.201900126.

(53) Abdel-Daim, A.; Ohura, K.; Imai, T. A Novel Quantification Method for Serine Hydrolases in Cellular Expression System Using Fluorophosphonate-Biotin Probe. European Journal of Pharmaceutical Sciences 2018, 114, 267–274. 10.1016/j.ejps.2017.12.016.

(54) MacDonald, J. I.; Munch, H. K.; Moore, T.; Francis, M. B. One-Step Site-Specific Modification of Native Proteins with 2-Pyridinecarboxyaldehydes. Nat Chem Biol 2015, 11 (5), 326–331. 10.1038/nchembio.1792.

(55) Wang, Y.; Xue, P.; Cao, M.; Yu, T.; Lane, S. T.; Zhao, H. Directed Evolution: Methodologies and Applications. Chem Rev 2021, 121 (20), 12384–12444. 10.1021/acs.chemrev.1c00260.

(56) Lane, M. D.; Seelig, B. Advances in the Directed Evolution of Proteins. Curr Opin Chem Biol 2014, 22, 129–136. 10.1016/j.cbpa.2014.09.013.

(57) Packer, M. S.; Liu, D. R. Methods for the Directed Evolution of Proteins. Nat Rev Genet 2015, 16 (7), 379–394. 10.1038/nrg3927.

