## Supplementary material for "Directed Chemical Evolution of Self-Assembling Artificial Proteins Utilizing a Supramolecular System": Supp Info

**Table of Contents:**


**1. Synthesis of protein conjugates**

**1.1. Summary of MALDI-TOF MS Data for protein conjugates.**

**1.2. MALDI-TOF MS Data of IEC peaks.**

**2. Synthesis of chemical probes and their intermediates**

**2.1. The general procedure for the synthesis of the hydrophilic spacer and its intermediates**

2.1.1. Procedure for synthesis of tetraethylene and octaethylene glycol spacer(1b/1d)

2.1.2. Procedure for synthesis of phosphonate hydrophilic spacer (1g/1h)

2.1.3. Procedure for synthesis of hydrophilic spacer(1i)

2.1.4. Synthesis of pyridinium intermediate

2.1.5. Synthesis of the hydrophilic spacer and its intermediates

**2.2. Procedure for synthesis of the hydrophobic tail**

2.2.1. Procedure for synthesis of 3-tail(3T) hydrophobic azide and its intermediates

2.2.2. Procedure for synthesis of 1-tail(1T) hydrophobic azide and its intermediates

**2.3. The general procedure of synthesis of the chemical probes.**

2.3.1. The general procedure of synthesis of the Thiol Maleimide probe

2.3.2. The general procedure of synthesis of the fluorophosphonate probe

2.3.3. The general procedure of synthesis of the N-terminal probe

3. **References**

#### 1. Synthesis of Protein Conjugates

##### 1.1. Summary of MALDI-TOF MS Data for Protein Conjugates.

| MW of Probe (Da) | Protein MW (Da) | Protein conjugate | MW of conjugate Cal. (Da) | MW of Conjugate Obt. (Da) |
|---|---|---|---|---|
| MI-TEG-C12-3T (997) | BSA (66627) | BSA-MI-TEG-C12-3T | 67624 | 67567 |
| | HSA (66674) | HSA-MI-TEG-C12-3T | 67671 | 67644 |
| MI-OEG-C12-3T (1173) | BSA (66627) | BSA-MI-OEG-C12-3T | 67800 | 67766 |
| | HSA (66674) | HSA-MI-OEG-C12-3T | 67874 | 67704 |
| MI-TEG-C12-1T (628) | BSA (66627) | BSA-MI-TEG-C12-1T | 67255 | 68594 |
| | HSA (66674) | HSA-MI-TEG-C12-1T | 67302 | 68568 |
| MI-TEG-C18-1T (712) | BSA (66627) | BSA-MI-TEG-C12-1T | 67339 | 67314 |
| | HSA (66674) | HSA-MI-TEG-C18-1T | 67386 | 67364 |
| FP-TEG-C12-3T (1012) | Chy (25486) | Chy-FP-TEG-C12-3T | 26498 | 26464 |
| | Sub (27285) | Sub-FP-TEG-C12-3T | 28297 | 28303 |
| | Pro K (28955) | Pro K-FP-TEG-C12-3T | 29967 | 29901 |
| FP-OEG-C12-3T (1188) | Chy (25486) | Chy-FP-OEG-C12-3T | 26674 | 26619 |
| | Sub (27285) | Sub-FP-OEG-C12-3T | 28473 | 28442 |
| | Pro K (28955) | Pro K-FP-OEG-C12-3T | 30143 | 30081 |
| FP-TEG-C12-1T (643) | Chy (25486) | Chy-FP-TEG-C12-1T | 26129 | 26321 |
| | Sub (27285) | Sub-FP-TEG-C12-1T | 27928 | 27928 |
| | Pro K (28955) | Pro K-FP-TEG-C12-1T | 29598 | 29541 |
| NPC-TEG-C12-3T (1105) | BSA (66627) | BSA-NPC-TEG-C12-3T | 67732 | 67933 |
| | HSA (66674) | HSA-NPC-TEG-C12-3T | 67779 | 67747 |
| | Chy (25486) | Chy-NPC-TEG-C12-3T | 26591 | 26616 |
| NPC-TEG-C12-1T (717) | BSA (66627) | BSA-NPC-TEG-C12-1T | 67344 | 67424 |
| | HSA (66674) | HSA-NPC-TEG-C12-1T | 67391 | 67434 |
| NPC-TEG-C18-1T (821) | BSA (66627) | BSA-NPC-TEG-C18-1T | 67448 | 67395 |
| | HSA (66674) | HSA-NPC-TEG-C18-1T | 67495 | 67466 |

**Supplementary Table 1:** Summary of MALDI-TOF MS Data for Protein Conjugates.

#### 1.2. MALDI-TOF MS Data of IEC fractions.

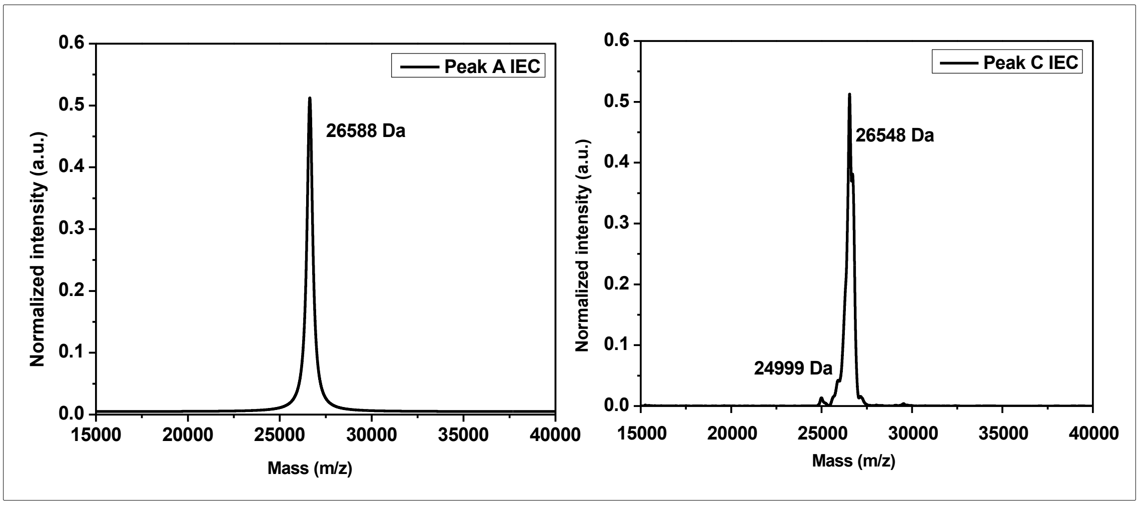


**Supplementary Fig. 1:** MALDI-TOF-MS Data of IEC peaks of Chy-FP-OEG-C12-3T protein conjugate.

The MALDI-TOF-MS data of IEC peaks for the Chy-FP-OEG-C12-3T protein conjugate indicates that peak A represents a pure protein conjugate, while peak C contains impurities along with the protein conjugate.

#### 2. Synthesis of chemical probes and their intermediates

All compounds were characterized using Nuclear Magnetic Resonance Spectroscopy (NMR), specifically $^{1}$H, $^{13}$C, and $^{1□}$F for fluorinated compounds. Additionally, they were analyzed using either Matrix-Assisted Laser Desorption/Ionization-Time of Flight Mass Spectrometry (MALDI-TOF MS) or High-Resolution Mass Spectrometry (HRMS). The NMR spectra were recorded on a Bruker 400 MHz, with tetramethylsilane (TMS) serving as an internal standard (chemical shifts in ppm). Unless otherwise noted, all $^{1}$H values are expressed in parts per million (ppm) and are referenced to the residual proton resonances of chloroform ($CHCl_3$) at 7.26 ppm in its respective deuterated solvent. All $^{13}$C and $^{19}$F NMR spectra were measured decoupled from the $^{1}$H nuclei and are also reported in parts per million (ppm).

All reactions were monitored using thin-layer chromatography on precoated silica gel plates, with detection through an ultraviolet (UV) lamp, phosphomolybdic acid (PMA), or ninhydrin staining. Chromatographic separations were carried out using column chromatography with normal-phase silica gel of 100-200 mesh size. The room temperature (RT) ranged from 21 °C to 35 °C. To describe the multiplicities of the signals, the following abbreviations were used:

s = singlet, d = doublet, dd = doublet of doublet, t = triplet, q = quartet, m = multiplet, and quint = quintet.

#### 2.1. The general procedure for the synthesis of the hydrophilic spacer and its intermediates

##### 2.1.1. Procedure for synthesis of tetraethylene and octaethylene glycol spacer(1b/1d)

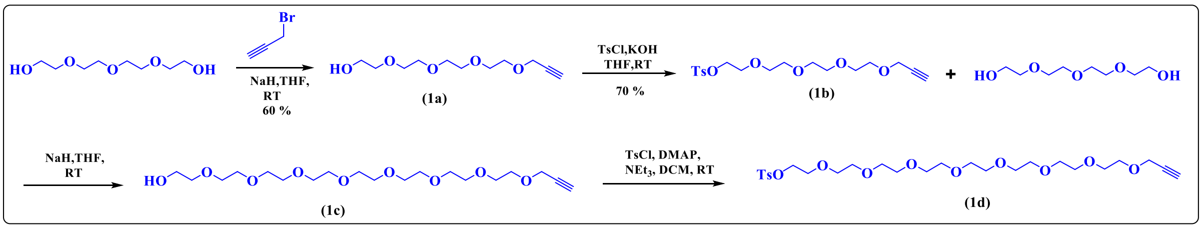


**Supplementary scheme 1:** Synthetic scheme for synthesizing tetraethylene and octaethylene glycol spacer.

The tetraethylene and octaethylene glycol spacer and their intermediates are synthesized using a reported procedure[1].

##### 2.1.2. Procedure for synthesis of hydrophilic spacer equipped with fluorophosphonate functionality (1g/1h)

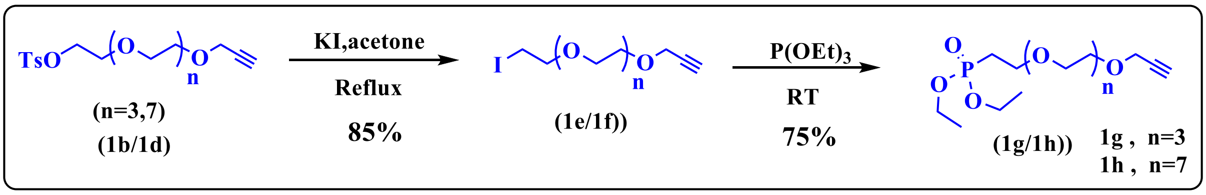


**Supplementary Scheme 2:** General scheme for the synthesis of tetraethylene and octaethylene glycol spacer.

The tetraethylene and octaethylene glycol spacer and their intermediates are synthesized using a reported procedure[1].

##### 2.1.3. Procedure for synthesis of hydrophilic spacer(1i)

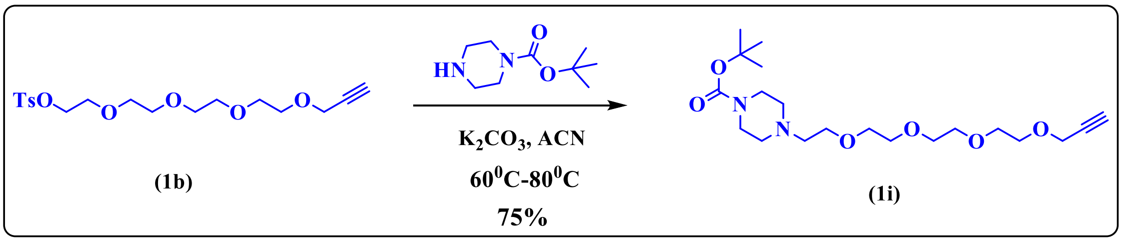


**Supplementary Scheme 3:** General scheme for the synthesis of hydrophilic spacer

The hydrophilic spacer (1i) is synthesized by using tosylated tetraethylene glycol(1b).

The detailed procedure is mentioned below:

###### 2.1.3.1. Synthesis of Compound (1i) – procedure A

Tosylate (1b) s(1 eq) and BOC-piperazine (4 eq), $K_2CO_3$ (2 eq) were dissolved in THF in an RBF. The mixture was then refluxed at 65°C for 16 hours. After the reaction was completed, water was added, and the mixture was extracted with DCM, concentrated, and purified by NPC using a MeOH/DCM solvent system **(Scheme 3)**.

##### 2.1.4. Synthesis of pyridinium intermediate

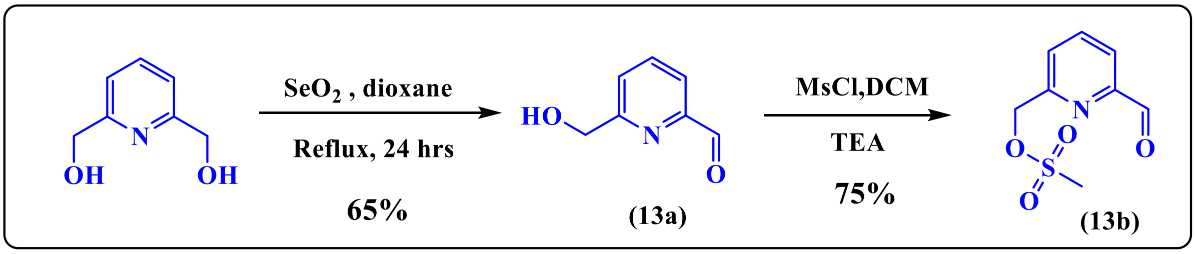


**Supplementary Scheme 4:** General scheme for the synthesis of pyridinium intermediate

The pyridinium intermediate is synthesized using 2,6-pyridinedimethanol.

The detailed procedure is mentioned below:

###### 2.1.4.1. Synthesis of Compound (13a)-procedure B

To a solution of 2,6-pyridinedimethanol (1 eq) in 1,4-dioxane, selenium dioxide ($SeO_2$) (0.5 eq) was added. The resulting mixture was sonicated for 5 minutes and then stirred at 80°C for 24 hours. The reaction was then cooled to room temperature and diluted with DCM. The mixture was filtered through Celite, and the filtrate was concentrated under reduced pressure. The resulting crude material was purified by NPC using EtOAc/Hexane to obtain a liquid that subsequently solidified to form an off-white solid (13a).

###### 2.1.4.2. Synthesis of Compound (13b)-procedure C

MsCl (2 eq) was added to the solution of alcohol (13a) (1eq) in DCM at 0 □C, $Et_3N$ was added to the above mixture, maintaining the temperature at 0 □C, the reaction mixture was stirred for 16 hrs at RT. Upon completion, the reaction mixture was extracted with DCM, and the combined organic layer was dried over $Na_2SO_4$ and concentrated under reduced pressure to obtain the crude product, which was then purified by NPC using EtOAc/Hexane to yield the mesylate product (13b).

##### 2.1.5. Synthesis of the hydrophilic spacer and its intermediates

###### 2.1.5.1. Compound (1i)

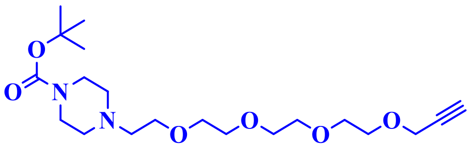

**(1i)**

Mol. formula: $C_{20}H_{36}N2O_6$

Mol. Weight: 400.52 g/mol

Physical appearance: brownish liquid

Yield: 60 %

The compound **(1i)** is synthesized using procedure A, using Tosylate **(1b)** (5.26 g, 13.61 mmol), BOC-piperazine (3.8 g,20.41 mmol), and $K_2CO_3$ (3.7 g, 27.22 mmol). The crude product was purified to obtain the pure product (4.90 g, 12.24 mmol, 60%) using NPC and a mixture of MeOH/DCM as the eluent. $R_f$ Value= 0.45 in 5% MeOH/DCM **$^1$H NMR (400 MHz, $CDCl_3$) δ:** 4.14 (d, *J* = 2.4 Hz, 2H), 3.67 – 3.52 (m, 15H), 3.42 – 3.33 (m, 4H), 2.61 (s, 1H), 2.53 (d, *J* = 11.4 Hz, 2H), 2.39 (dt, *J* = 6.2, 3.4 Hz, 5H), 1.39 (s, 9H). **$^{13}$C NMR (100 MHz, $CDCl_3$) δ:** 79.63, 79.56, 74.62, 70.57, 70.54, 70.38, 70.32, 69.25, 69.07, 68.65, 58.37, 57.83, 53.32, 28.41. **MALDI-TOF MS (M+Na):** 423.56 g/mol.

###### 2.1.5.2. Compound (13a)

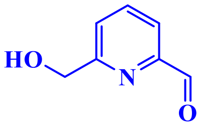

**(13a)**

Mol. formula: $C_7H_7NO_2$

Mol. Weight: 137.14 g/mol

Physical appearance: white solid

Yield: 60 %

The compound **(13a)** is synthesized using procedure B, using 2,6-pyridinedimethanol (7 g, 50.3 mmol) and $SeO_2$ (2.7 g, 25.15 mmol). The pure product is a liquid that solidified to an off-white solid later (2.06 g, 15.09 mmol, 60%), with an $R_f$ Value of 0.45 in 5% MeOH/DCM. **$^{1}$H NMR (400 MHz, $CDCl_3$) δ:** 10.05 (d, $J$ = 0.6 Hz, 1H), 7.92 – 7.79 (m, 2H), 7.58 – 7.46 (m, 1H), 4.86 (s, 2H), 3.67 (s, 1H). **$^{13}$C NMR (100 MHz, $CDCl_3$) δ:** 192.82, 160.09, 160.06, 151.37, 137.84, 137.56, 124.68, 120.79, 120.35, 69.23, 66.82, 64.35, 63.91, 53.23.

**2.1.5.3. Compound (13b)**

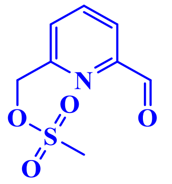

**(13b)**

Mol. formula: $C_8H_9NO_4S$

Mol. Weight: 137.13 g/mol

Physical appearance: white solid

Yield: 60 %

The compound **(13b)** is synthesized using procedure C, using methane sulfonyl chloride (2.94 g, 20.4 mmol), $Et_3N$ (4.41 g, 43.7 mmol), and compound **(13a).** After purification, a white solid was obtained as the pure product (1.19 g, 8.7 mmol, 60%), with an $R_f$ Value of 0.45 in 50% EtOAc/Hexane. **$^{1}$H NMR (400 MHz, $CDCl_3$) δ:** 9.99 (d, $J$ = 0.8 Hz, 1H), 7.95 – 7.83 (m, 2H), 7.68 (dd, $J$ = 7.2, 1.6 Hz, 1H), 4.71 (s, 2H). **$^{13}$C NMR (100 MHz, $CDCl_3$) δ:** 192.98, 192.39, 157.45, 152.29, 138.27, 137.99, 127.14, 46.07.

**2.2. Procedure for synthesis of the hydrophobic tail**

**2.2.1. Procedure for synthesis of 3-tail(3T) hydrophobic azide and its intermediates**

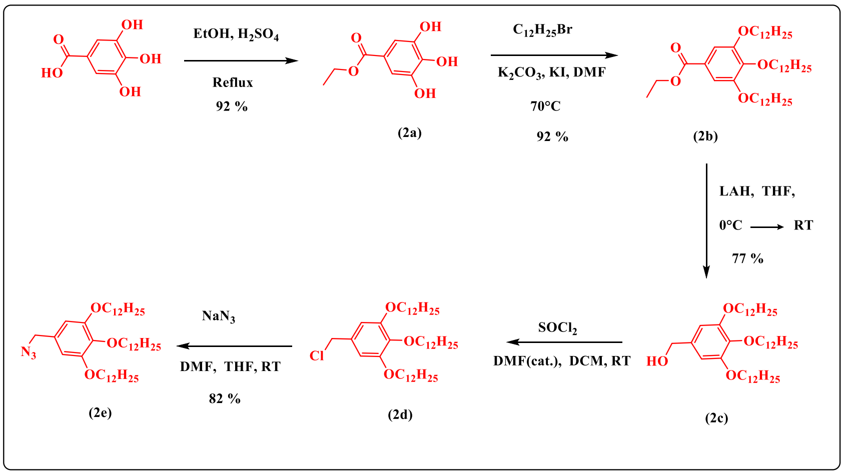


**Supplementary Scheme 5:** General scheme for the synthesis of 3-tail(3T) hydrophobic azide and its intermediates

The 3-tail(3T) hydrophobic azide and its intermediates are synthesized by using a reported procedure[1].

**2.2.2. Procedure for synthesis of 1-tail(1T) hydrophobic azide and their intermediates**

1-tail(1T) hydrophobic azide and its intermediates are synthesized using a reported procedure[1].

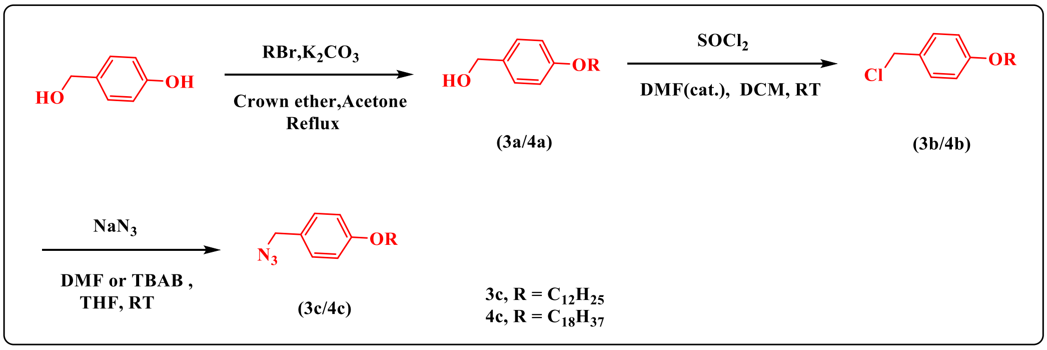


**Supplementary Scheme 6:** General scheme for the synthesis of 1-tail(1T) hydrophobic azide and its intermediates

**2.3. The general procedure of synthesis of the chemical probes.**

**2.3.1. The general procedure of synthesis of the Thiol Maleimide probe**

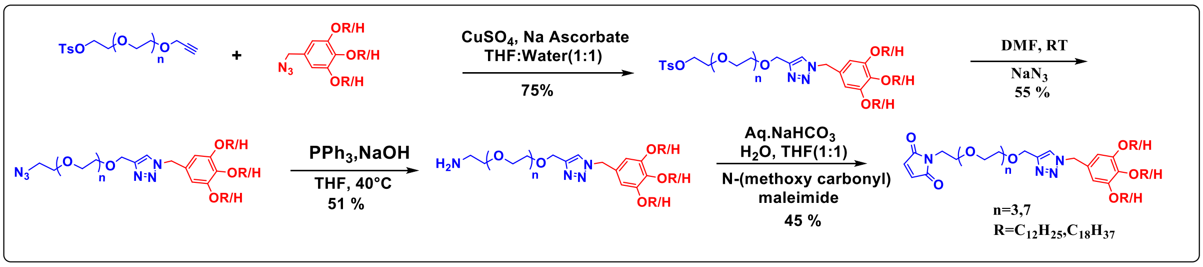


**Supplementary Scheme 7:** General scheme for the synthesis of the Thiol Maleimide probe.

All the chemical probes were synthesized using a [2 + 3] dipolar cycloaddition (click reaction) involving hydrophilic alkynes and hydrophobic azides. This reaction was performed in the presence of sodium ascorbate and copper (II) sulphate ($CuSO_4$) in a 50% THF/$H_2O$ solution. The azidation of tosylated click products was done using $NaN_3$ in DMF. The azide was then reduced to an amine using triphenylphosphine ($PPh_3$). The final step involves converting the amine into a maleimide using N-methoxycarbonyl maleimide. The detailed procedure is mentioned below:

##### 2.3.1.1. Synthesis of Click Product-Procedure D

Hydrophilic alkyne (1 eq) and hydrophobic azide (4 eq) were dissolved in degassed THF and stirred until a clear solution was obtained. Then, degassed $H_2O$ was added and stirred vigorously for an additional 10 minutes. Freshly prepared 1M Na ascorbate (0.05 eq) and 1M $CuSO_4$ (0.1 eq) were added to the reaction mixture thrice in intervals of 45 mins and allowed to react for 16 hrs at RT. Upon completion, the reaction mixture was extracted with DCM, and the combined organic layer was dried over $Na_2SO_4$ and concentrated under reduced pressure to obtain the crude product. The crude product was then purified by NPC using a MeOH/DCM solvent system to yield the clicked product.

##### 2.3.1.2. Synthesis of Azide Product-Procedure E

The amphiphilic tosylate was dissolved in DMF at RT, then $NaN_3$(2 eq) was added to the reaction mixture and allowed to react at RT for 12 hrs. After this, the reaction mixture was concentrated and purified by NPC without any workup.

##### 2.3.1.3. Synthesis of Amine Product-Procedure F

Above, the azide was dissolved in THF, and then a $PPh_3$ (4 eq) solution in THF was added slowly and stirred at 40°C. After 10 minutes, 1 M NaOH was added while maintaining the pH of the solution at 11. The reaction mixture was stirred for 16 hours at 40°C. After the reaction was completed, the RM was neutralized with acidic water, followed by DCM extraction, and then purified using NPC.

**2.3.1.4. Synthesis of Maleimide Product-Procedure G**

Amine was dissolved in a (1:1) THF: sat. Aq. solution of $NaHCO_3$ and cooled on an ice bath. Then N- (methoxycarbonyl) maleimide (1.2 eq) was added in portions under stirring. The mixture was stirred for 2 hours at 0°C, followed by 6 hours at 30°C. After extraction with DCM, the organic phase was dried over anhydrous $Na_2SO_4$, concentrated, and purified by NPC using a (MeOH/DCM) mixture, yielding the products (5d/6d).

**2.3.1.5. Synthesis of Maleimide probe and its intermediates**

**2.3.1.5.1. Compound (5a)**

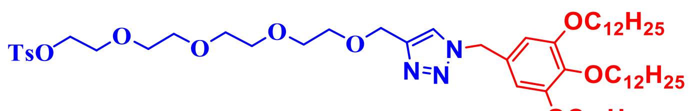


**(5a)**

Mol. formula: $C_{61}H_{105}N_3O_{10}S$

Mol. weight: 1071.75 g/mol.

Physical appearance: yellow waxy solid

Yield: 85%

The compound **(5a)** is synthesized using general procedure D, using Alkyne **(1b)** (0.250 g, 0.646 mmol), azide **(2e)** (3.5 g, 2.58 mmol), $CuSO_4$ (16 mg, 0.06 mmol), and Na ascorbate (6 mg, 0.032 mmol). The resulting crude material was purified by NPC using MeOH/DCM (0.600 g, 0.55 mmol, 85 %), $R_f$ value = 0.33 in 10 % MeOH/DCM.**$^1$H NMR (400 MHz, $CDCl_3$) δ:**7.81 – 7.75 (m, 2H), 7.54 – 7.47 (m, 1H), 7.33 (d, *J* = 8.0 Hz, 2H), 6.45 (s, 2H), 5.37 (s, 2H), 4.65 (s, 2H), 4.22 – 4.11 (m, 2H), 3.91 (q, *J* = 6.3 Hz, 6H), 3.69 – 3.55 (m, 14H), 2.43 (s, 3H), 1.78 – 1.71 (m, 7H), 1.48 – 1.40 (m, 6H), 1.25 (s, 48H), 0.87 (t, *J* = 6.7 Hz, 9H).**$^{13}$C NMR (100 MHz, $CDCl_3$) δ:**153.69, 144.93, 138.58, 133.11, 129.96, 129.52, 128.10, 106.89, 73.59, 70.85, 70.71, 70.66, 70.62, 69.89, 69.37, 68.80, 64.84, 54.66, 32.06, 30.45, 29.88, 29.86, 29.83, 29.79, 29.77, 29.73, 29.55, 29.52, 29.50, 26.24, 26.21, 22.82, 21.76, 14.24. **MALDI-TOF MS (M+Na):**1095.22 g/mol.

##### 2.3.1.5.2. Compound (5b)

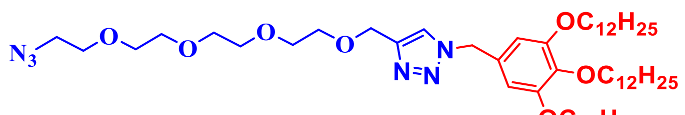

**(5b)**

Mol. formula: $C_{54}H_{98}N_6O_7$

Mol. weight: 943.41 g/mol.

Physical appearance: yellow waxy solid

Yield: 76 %

The compound **(5b)** is synthesized using general procedure E, using, compound (**5a)** (0.600 g, 0.559 mmol) and $NaN_3$ (0.072 g, 1.11 mmol). Reaction mixture was purified by NPC without any workup (0.420 g, 76 %), $R_f$ value = 0.40 in 10 % MeOH/DCM.**$^1$H NMR (400 MHz, $CDCl_3$) δ:**7.48 (s, 1H), 6.45 (s, 2H), 5.37 (s, 2H), 4.66 (s, 2H), 3.91 (q, $J$ = 6.3 Hz, 6H), 3.72 – 3.58 (m, 14H), 3.37 (t, $J$ = 5.1 Hz, 2H), 1.80 – 1.69 (m, 6H), 1.45 (dq, $J$ = 8.4, 4.8 Hz, 6H), 1.33 – 1.25 (m, 49H), 0.89 – 0.85 (m, 9H).**$^{13}$C NMR (100 MHz, $CDCl_3$) δ:**153.71, 145.60, 138.61, 129.48, 122.55, 106.90, 77.36, 73.60, 70.80, 70.77, 70.74, 70.68, 70.62, 70.15, 69.90, 69.38, 64.85, 54.61, 50.80, 32.07, 32.05, 30.45, 29.88, 29.86, 29.83, 29.82, 29.79, 29.77, 29.73, 29.55, 29.52, 29.50, 26.24, 26.21, 22.82, 14.24.**MALDI-TOF MS (M+Na):** 966.04 g/mol.

##### 2.3.1.5.3. Compound (5c)

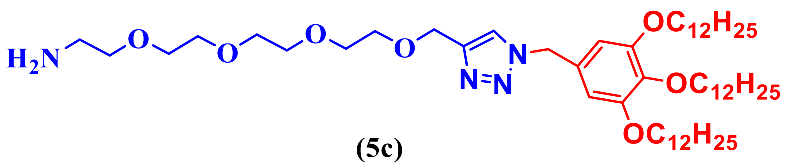

**(5c)**

Mol. formula: $C_{54}H_{100}N_4O_7$

Mol. weight: 916.74 g/mol.

Physical appearance: yellow waxy solid

Yield: 55 %

The compound **(5c)** is synthesized using general procedure F, using, compound **(5b)** (0.110 gm, 0.116 mmol) and $PPh_3$ (0.112 gm, 0.466 mmol). The reaction mixture was extracted with DCM and purified using NPC (0.058 gm,0.063 mmol, 55%), $R_f$ value = 0.40 in 10 % MeOH/DCM.**$^{1}$H NMR (400 MHz, $CDCl_3$) δ:** 7.56 (s, 1H), 6.48 (s, 2H), 5.39 (s, 2H), 4.64 (s, 2H), 3.91 (t, $J$ = 6.5 Hz, 6H), 3.69 (d, $J$ = 1.3 Hz, 2H), 3.65 – 3.57 (m, 12H), 2.84 (t, $J$ = 5.0 Hz, 2H), 1.79 – 1.69 (m, 6H), 1.47 – 1.40 (m, 6H), 1.25 (s, 48H), 0.87 (t, $J$ = 6.8 Hz, 9H).**$^{13}$C NMR (100 MHz, $CDCl_3$) δ:** 153.68, 144.95, 138.55, 129.45, 122.79, 106.95, 73.58, 71.00, 70.48, 70.39, 70.37, 70.20, 70.02, 69.36, 64.67, 54.66, 41.06, 32.06, 32.05, 30.44, 29.87, 29.85, 29.83, 29.82, 29.78, 29.76, 29.73, 29.63, 29.55, 29.51, 29.49, 26.23, 26.22, 22.81, 14.24.**MALDI-TOF MS (M+Na):** 940.03 g/mol.

**2.3.1.5.4. Compound (5d)**

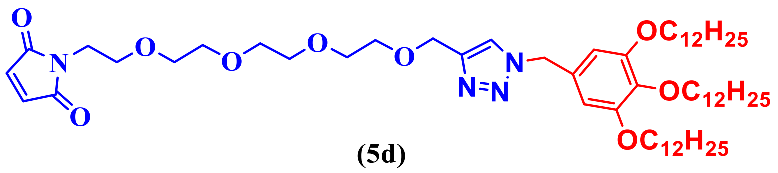


**(5d)**

Mol. formula: $C_{58}H_{100}N_4O_9$

Mol. weight: 997.46 g/mol.

Physical appearance: brownish oil

Yield: 45 %

The compound **(5d)** is synthesized using general procedure G, using, compound **(5c)** (2.1 gm, 2.29 mmol) and N-(methoxycarbonyl) maleimide (0.424 gm, 2.74 mmol). Purification by NPC using MeOH/DCM yielded the product **(5d)** as a brownish oil. (1.02 gm, 1.03 mmol 45 %), $R_f$ value = 0.35 in 5 % MeOH/DCM.**$^{1}$H NMR (400 MHz, $CDCl_3$) δ:** 7.49 (s, 1H), 6.68 (s, 2H), 6.46 (s, 2H), 5.38 (s, 2H), 4.66 (s, 2H), 3.94 – 3.88 (m, 6H), 3.71 – 3.56 (m, 16H), 1.81 – 1.76 (m, 4H), 1.71 (d, $J$ = 7.9 Hz, 2H), 1.47 – 1.41 (m, 6H), 1.25 (d, $J$ = 2.4 Hz, 48H), 0.87 (d, $J$ = 7.0 Hz, 9H).**$^{13}$C NMR (400 MHz, $CDCl_3$ ) δ:** 170.79, 153.72, 145.61, 138.63, 134.28, 129.46, 122.63, 106.93, 77.37, 73.61, 70.72, 70.66, 70.61, 70.17, 69.92, 69.39, 67.96, 64.81, 54.66, 37.26, 32.07, 30.46, 29.89, 29.88, 29.84, 29.80, 29.78, 29.75, 29.65, 29.57, 29.53, 29.51, 26.25, 26.23, 22.83, 14.26.**MALDI-TOF MS (M+Na):** 1019.46 g/mol.

#### 2.3.1.5.5. Compound (6a)

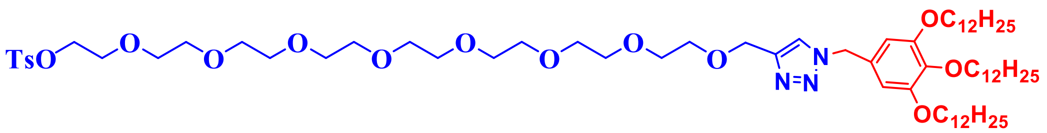


**(6a)**

Mol. formula: $C_{69}H_{12}1N_3O_{14}S$

Mol. weight: 1248.79 g/mol.

Physical appearance: yellow waxy solid

Yield: 85 %

The compound **(6a)** is synthesized using general procedure D using Alkyne **(1d)** (1.63 g, 2.91 mmol), azide **(2e)** (7.98 g, 11.6 mmol), $CuSO_4$ (72 mg, 0.291 mmol), and Na ascorbate (28 mg, 0.145 mmol). The resulting crude material was purified by NPC using MeOH/DCM (3.08 g, 2.47 mmol, 85 %), $R_f$ value = 0.33 in 10 % MeOH/DCM. **$^1$H NMR (400 MHz, $CDCl_3$) δ:** 7.81 – 7.77 (m, 2H), 7.54 (s, 1H), 7.34 (d, *J* = 8.1 Hz, 2H), 6.45 (s, 2H), 5.38 (s, 2H), 4.65 (s, 2H), 4.17 – 4.13 (m, 2H), 3.91 (h, *J* = 5.0 Hz, 6H), 3.69 – 3.57 (m, 30H), 2.44 (s, 3H), 1.80 – 1.75 (m, 4H), 1.69 (d, *J* = 6.7 Hz, 2H), 1.43 (q, *J* = 6.7 Hz, 6H), 1.27 (d, *J* = 7.7 Hz, 48H), 0.87 (t, *J* = 6.7 Hz, 9H).**$^{13}$C NMR (100 MHz, $CDCl_3$) δ:** 153.71, 144.93, 138.64, 133.15, 129.96, 129.41, 128.12, 106.93, 77.36, 73.61, 70.87, 70.73, 70.67, 70.64, 69.93, 69.39, 68.82, 32.06, 30.46, 29.89, 29.87, 29.84, 29.80, 29.78, 29.74, 29.56, 29.53, 29.51, 26.25, 26.22, 22.83, 21.78, 14.25.**MALDI-TOF MS (M+K):**1287.06 g/mol.

#### 2.3.1.5.6. Compound (6b)

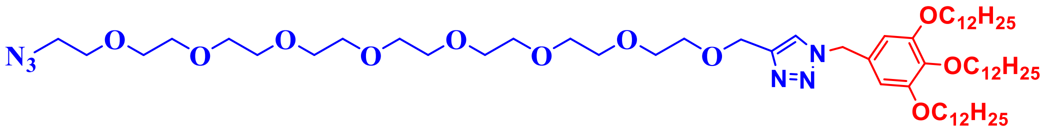


**(6b)**

Mol. formula: $C_{62}H_{114}N_6O_{11}$

Mol. weight: 1119.63 g/mol.

Physical appearance: yellow waxy solid

Yield: 45%

The compound **(6b)** is synthesized using general procedure E, using, compound **(6a)** (6 g, 4.80 mmol) and $NaN_3$ (0.62 g, 9.6 mmol). The reaction mixture was purified by NPC (2.4 g, 2.16 mmol, 45 %), $R_f$ value = 0.42 in 10 % MeOH/DCM. **$^1$H NMR (400 MHz, $CDCl_3$) δ:** 7.48 (s, 1H), 6.44 (s, 2H), 5.37 (s, 2H), 4.65 (s, 2H), 3.93 – 3.85 (m, 6H), 3.68 – 3.60 (m, 30H), 3.37 (t, $J$ = 5.1 Hz, 2H), 1.80 – 1.68 (m, 6H), 1.43 (tt, $J$ = 12.1, 4.6 Hz, 6H), 1.26 (d, $J$ = 16.1 Hz, 49H), 0.86 (t, $J$ = 6.8 Hz, 9H).**$^{13}$C NMR (400 MHz, $CDCl_3$) δ:** 153.66, 145.55, 138.56, 129.48, 122.58, 106.85, 73.55, 70.78, 70.75, 70.71, 70.64, 70.62, 70.57, 70.12, 69.85, 69.33, 64.80, 54.57, 50.77, 32.03, 32.02, 30.41, 29.85, 29.83, 29.80, 29.79, 29.76, 29.73, 29.70, 29.51, 29.48, 29.46, 26.25, 26.21, 26.18, 22.78, 14.21.**MALDI-TOF MS (M+Na):**1142.14 g/mol.

##### 2.3.1.5.7. Compound (6c)

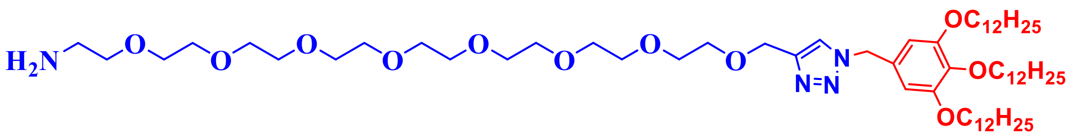


**(6c)**

Mol. formula: $C_{62}H_{116}N_4O_{11}$

Mol. weight: 1093.63 g/mol.

Physical appearance: yellow waxy solid

Yield: 51 %

The compound **(6c)** is synthesized using general procedure F, using, compound **(6b)** (2.41 g, 2.14 mmol) and $PPh_3$ (2.24 g, 8.56 mmol) dissolved in THF (5 ml). The reaction mixture was extracted with DCM and purified using NPC (1.34 g,1.22 mmol, 51 %), $R_f$ value = 0.29 in 10% MeOH/DCM. **$^1$H NMR (400 MHz, $CDCl_3$) δ:** 7.49 (s, 1H), 6.44 (s, 2H), 5.36 (s, 2H), 4.64 (s, 2H), 3.92 – 3.87 (m, 6H), 3.67 – 3.59 (m, 29H), 3.50 (t, $J$ = 5.2 Hz, 2H), 2.86 (q, $J$ = 4.2 Hz, 1H), 1.79 – 1.67 (m, 6H), 1.42 (dq, $J$ = 11.2, 7.0 Hz, 6H), 1.25 (d, $J$ = 6.6 Hz, 49H),

0.86 (t, $J$ = 6.7 Hz, 9H).$^{13}$**C NMR (400 MHz, $CDCl_3$) δ:** 153.65, 145.51, 138.54, 129.51, 122.61, 106.84, 73.55, 73.05, 70.63, 70.61, 70.55, 70.33, 69.83, 69.33, 64.79, 54.55, 41.75, 32.02, 30.41, 29.84, 29.82, 29.79, 29.75, 29.73, 29.69, 29.51, 29.48, 29.46, 26.20, 26.18, 22.78, 14.21.**MALDI-TOF MS (M+Na):** 1116.07 g/mol.

##### 2.3.1.5.8. Compound (6d)

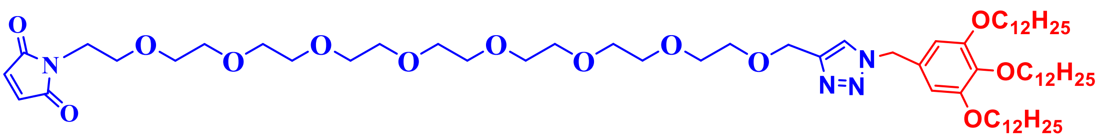


**(6d)**

Mol. formula: $C_{66}H_{116}N_4O_{13}$

Mol. weight: 1173.67 g/mol.

Physical appearance: yellow oil

Yield: 45 %

The compound **(6d)** is synthesized using general procedure G, using, compound **(6d)** (2.34 g, 2.14 mmol) and N-(methoxycarbonyl) maleimide (0.398 g, 2.56 mmol). Purification by NPC using MeOH/DCM yielded the product as a yellow oil. (1.12 g, 0.96 mmol 45 %), $R_f$ value = 0.33 in 10 % MeOH/DCM.$^{1}$**H NMR (400 MHz, $CDCl_3$) δ:** 7.49 (s, 1H), 6.68 (s, 2H), 6.44 (s, 2H), 5.36 (s, 2H), 4.64 (s, 2H), 3.89 (h, $J$ = 5.3 Hz, 6H), 3.73 – 3.53 (m, 32H), 1.73 (dq, $J$ = 18.8, 7.1 Hz, 6H), 1.42 (dq, $J$ = 11.4, 6.7 Hz, 6H), 1.24 (s, 48H), 0.85 (t, $J$ = 6.6 Hz, 9H).$^{13}$**C NMR (100 MHz, $CDCl_3$) δ:** 170.72, 153.63, 145.53, 138.53, 134.23, 129.40, 122.60, 106.84, 73.51, 70.66, 70.60, 70.58, 70.52, 70.10, 69.82, 69.30, 67.87, 64.70, 54.57, 37.18, 31.99, 30.38, 29.81, 29.79, 29.76, 29.72, 29.70, 29.66, 29.52, 29.48, 29.45, 29.43, 26.22, 26.18, 26.15, 22.75, 14.18**MALDI-TOF MS (M+K):**1212.09 g/mol.

##### 2.3.1.5.9. Compound (7a)

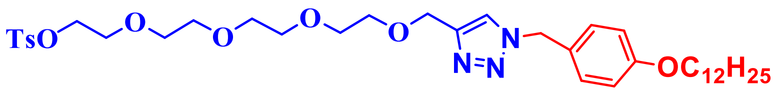

**(7a)**

Mol. formula: $C_{37}H_{57}N_3O_8S$

Mol. weight: 703.94 g/mol.

Physical appearance: Yellowish waxy solid

Yield:  75%

The compound **(7a)** is synthesized using general procedure E using Alkyne **(1d)** (2.31 g, 5.98 mmol), azide **(3c)** (7.8 g, 23.6 mmol), $CuSO_4$ (15 mg, 0.59 mmol), and Na ascorbate (59 mg, 0.299 mmol). The resulting crude material was purified by NPC using MeOH/DCM (3.1 g, 4.48 mmol, 75 %), with an $R_f$ value of 0.35 in 10% MeOH/DCM. **$^1$H NMR (400 MHz, $CDCl_3$) δ:** 7.80 – 7.69 (m, 2H), 7.55 – 7.46 (m, 1H), 7.29 (d, $J$ = 8.1 Hz, 2H), 7.17 (d, $J$ = 8.3 Hz, 2H), 6.90 – 6.73 (m, 2H), 5.39 (s, 2H), 4.60 (s, 2H), 4.20 – 4.03 (m, 2H), 3.89 (t, $J$ = 6.6 Hz, 2H), 3.71 – 3.48 (m, 14H), 2.40 (s, 3H), 1.80 – 1.65 (m, 2H), 1.41 (dd, $J$ = 10.1, 5.4 Hz, 2H), 1.33 – 1.21 (m, 17H), 0.84 (t, $J$ = 6.8 Hz, 3H). **$^{13}$C NMR (100 MHz, $CDCl_3$) δ:** 159.46, 144.80, 132.95, 129.82, 129.67, 129.39, 127.93, 126.33, 114.94, 114.27, 70.68, 70.54, 70.48, 70.45, 69.68, 69.27, 68.62, 68.08, 64.68, 53.75, 31.88, 31.48, 30.14, 29.66, 29.62, 29.60, 29.56, 29.54, 29.35, 29.31, 29.17, 25.99, 22.66, 22.30, 21.61, 14.11. **MALDI-TOF MS (M+K):**742.53 g/mol.

##### 2.3.1.5.10. Compound (7b)

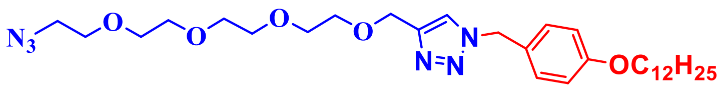

**(7b)**

Mol. formula: $C_{30}H_{50}N_6O_5$

Mol. weight: 574.38 g/mol.

Physical appearance: yellow waxy solid

Yield: 70%

The compound **(7b)** is synthesized using general procedure F, using, compound **(7a)** (3.1 g, 4.48 mmol) and $NaN_3$ (0.582 g, 8.96 mmol). The reaction mixture was purified by NPC (1.8 g, 3.13 mmol, 70 %), with an $R_f$ value of 0.42 in 10% MeOH/DCM. **$^1$H NMR (400 MHz, $CDCl_3$) δ:** 7.43 (s, 1H), 7.23 – 7.13 (m, 2H), 6.89 – 6.82 (m, 2H), 5.41 (s, 2H), 4.62 (s, 2H), 3.92 (t, *J* = 6.6 Hz, 2H), 3.69 – 3.58 (m, 14H), 3.35 (t, *J* = 5.0 Hz, 2H), 1.82 – 1.68 (m, 2H), 1.47 – 1.19 (m, 22H), 0.86 (t, *J* = 6.7 Hz, 4H). **$^{13}$C NMR (100 MHz, $CDCl_3$) δ:** 159.57, 145.51, 130.05, 129.88, 129.75, 128.02, 126.35, 122.31, 115.32, 115.05, 114.37, 70.73, 70.70, 70.67, 70.61, 70.55, 70.07, 69.78, 69.32, 68.72, 68.17, 64.76, 53.77, 50.73, 31.96, 31.56, 31.48, 30.20, 29.75, 29.71, 29.68, 29.64, 29.62, 29.43, 29.40, 29.25, 26.07, 22.74, 14.18. **MALDI-TOF MS (M+K):**613.4 g/mol.

**2.3.1.5.11. Compound (7c)**

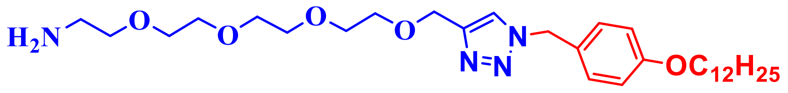


**(7c)**

Mol. formula: $C_{30}H_{52}N_4O_5$

Mol. weight: 548.77 g/mol.

Physical appearance: yellow waxy solid

Yield: 55 %

The compound **(7c)** is synthesized using general procedure G, using, compound **(7b)** (1.8 g, 3.13 mmol) and $PPh_3$ (3.2 g, 12.52 mmol). The reaction mixture was extracted with DCM and purified using NPC (0.943 g, 1.72 mmol, 55%), yielding a product with an $R_f$ value of 0.25 in 10% MeOH/DCM. **$^1$H NMR (400 MHz, $CDCl_3$) δ:** 7.58 (s, 1H), 7.25 – 7.21 (m, 2H), 6.88 – 6.83 (m, 2H), 5.44 (s, 2H), 4.63 (s, 2H), 3.92 (t, *J* = 6.6 Hz, 2H), 3.75 – 3.57 (m, 14H), 2.89 – 2.78 (m, 3H), 1.79 – 1.70 (m, 2H), 1.42 (td, *J* = 8.7, 4.4, 2.3 Hz, 3H), 1.37 – 1.25 (m, 17H), 0.87 (s, 3H). **$^{13}$C NMR (100 MHz, $CDCl_3$) δ:** 159.64, 144.77, 129.94, 126.38,

124.48, 123.58, 122.69, 119.14, 118.96, 115.10, 70.40, 70.31, 70.28, 70.13, 70.00, 68.25, 64.57, 53.91, 46.03, 40.76, 35.09, 34.55, 32.03, 31.62, 31.55, 30.30, 30.25, 29.81, 29.77, 29.74, 29.71, 29.68, 29.50, 29.46, 29.31, 26.14, 22.80, 14.23, 10.04, 8.04.**MALDI-TOF MS (M+Na):** 571.5 g/mol.

**2.3.1.5.12. Compound (7d)**

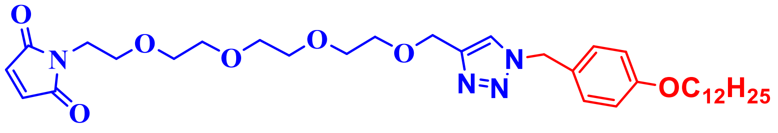


**(7d)**

Mol. formula: $C_{34}H_{52}N_4O_7$

Mol. weight: 628.81 g/mol.

Physical appearance: yellow oil

Yield: 45 %

The compound **(7d)** is synthesized using general procedure G, using, compound **(7c)** (0.943 g, 1.72 mmol) and N-(methoxycarbonyl) maleimide (1.28 g, 8.2 mmol). Purification by NPC using MeOH/DCM yielded the product as a yellow oil (0.486 g, 0.774 mmol, 45%), with an $R_f$ value of 0.36 in 10% MeOH/DCM. **$^1$H NMR (400 MHz, $CDCl_3$) δ:** 7.46 (s, 1H), 7.19 (d, $J$ = 8.4 Hz, 2H), 6.85 (d, $J$ = 8.4 Hz, 2H), 6.66 (s, 2H), 5.41 (s, 2H), 4.62 (s, 2H), 3.91 (t, $J$ = 6.5 Hz, 2H), 3.68 (q, $J$ = 7.1, 6.5 Hz, 2H), 3.64 – 3.55 (m, 12H), 1.79 – 1.70 (m, 2H), 1.44 – 1.38 (m, 3H), 1.36 – 1.24 (m, 16H), 0.85 (t, $J$ = 6.6 Hz, 4H). **$^{13}$C NMR (100 MHz, $CDCl_3$) δ:** 170.74, 159.57, 145.46, 139.32, 134.21, 129.76, 126.37, 122.44, 115.94, 115.05, 114.14, 70.63, 70.56, 70.51, 70.09, 69.77, 68.18, 67.88, 64.71, 53.79, 53.52, 52.31, 37.19, 34.84, 33.88, 31.99, 31.98, 31.72, 31.57, 30.36, 30.20, 29.76, 29.72, 29.69, 29.66, 29.63, 29.57, 29.55, 29.45, 29.42, 29.41, 29.26, 29.22, 29.01, 26.08, 25.67, 24.95, 22.75, 14.19.**MALDI-TOF MS (M+Na):** 651.51 g/mol.

**2.3.1.5.13. Compound (8a)**

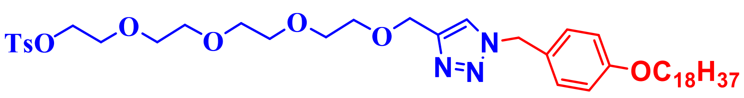

**(8a)**

Mol. formula: $C_{43}H_{69}N_3O_8S$

Mol. weight: 788.10 g/mol.

Physical appearance: Waxy white solid

Yield: 85%

The compound **(8a)** is synthesized using general procedure D using Alkyne **(1b)** (1.89 g, 4.89 mmol), azide **(4c)** (7.8 g, 19.56 mmol), $CuSO_4$ (123 mg, 0.48 mmol), and Na ascorbate (48 mg, 0.244 mmol). The resulting crude material was purified by NPC using MeOH/DCM (3.27 g, 4.15 mmol, 85 %), with an $R_f$ value of 0.34 in 10% MeOH/DCM. **$^1$H NMR (400 MHz, $CDCl_3$) δ:** 7.76 – 7.68 (m, 2H), 7.44 (s, 1H), 7.29 (d, *J* = 8.1 Hz, 2H), 7.21 – 7.07 (m, 2H), 6.88 – 6.78 (m, 2H), 5.39 (s, 2H), 4.60 (s, 2H), 4.16 – 4.06 (m, 2H), 3.89 (t, *J* = 6.6 Hz, 2H), 3.67 – 3.47 (m, 14H), 2.40 (s, 3H), 1.77 – 1.68 (m, 2H), 1.44 – 1.36 (m, 2H), 1.28 (s, 29H), 0.84 (t, *J* = 6.7 Hz, 3H). **$^{13}$C NMR (100 MHz, $CDCl_3$) δ:** 166.06, 151.99, 151.39, 139.58, 137.51, 136.43, 136.27, 136.00, 135.41, 134.54, 132.95, 128.97, 121.80, 121.54, 84.08, 83.76, 83.44, 77.29, 77.16, 77.10, 77.07, 77.06, 76.29, 75.88, 75.23, 74.68, 72.12, 71.49, 71.25, 60.26, 38.51, 38.09, 38.02, 37.15, 36.91, 36.28, 29.27, 28.21, 20.72, 20.32. **MALDI-TOF MS (M+Na):**811.6 g/mol.

##### 2.3.1.5.14. Compound (8b)

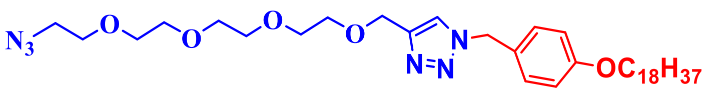

**(8b)**

Mol. formula:$C_{36}H_{62}N_6O_5$

Mol. weight: 658.93 g/mol.

Physical appearance: Waxy white solid

Yield: 55%

The compound **(8b)** is synthesized using general procedure E, using, compound **(8a)** (3.27 g, 4.15 mmol) and $NaN_3$ (0.539 g, 8.3 mmol). The reaction mixture was purified by NPC (1.50 g, 2.28 mmol, 55 %), with an $R_f$ value of 0.42 in 10% MeOH/DCM. **$^1$H NMR (400 MHz, $CDCl_3$) δ:** 7.43 (d, $J$ = 3.7 Hz, 1H), 7.15 (d, $J$ = 8.6 Hz, 2H), 6.87 – 6.75 (m, 2H), 5.36 (s, 2H), 4.58 (d, $J$ = 3.7 Hz, 2H), 3.87 (t, $J$ = 6.5 Hz, 2H), 3.67 – 3.44 (m, 14H), 3.42 – 3.20 (m, 2H), 1.82 – 1.61 (m, 2H), 1.39 (dd, $J$ = 10.7, 5.1 Hz, 2H), 1.21 (s, 28H), 0.90 – 0.73 (m, 3H). **$^{13}$C NMR (100 MHz, $CDCl_3$) δ:** 159.31, 159.29, 145.22, 144.60, 132.87, 129.68, 129.50, 127.78, 126.50, 126.29, 126.25, 122.24, 122.19, 114.78, 71.16, 70.53, 70.49, 70.47, 70.44, 70.41, 70.37, 70.32, 69.85, 69.55, 69.15, 68.47, 67.90, 64.51, 64.48, 53.47, 50.48, 31.77, 29.55, 29.51, 29.45, 29.43, 29.24, 29.21, 29.05, 25.88, 22.53, 21.44, 13.98.**MALDI-TOF MS (M+K):** 681.34 g/mol.

**2.3.1.5.15. Compound (8c)**

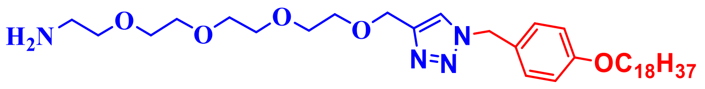


**(8c)**

Mol. formula: $C_{62}H_{116}N_4O_{11}$

Mol. weight: 632.93 g/mol.

Physical appearance: Waxy white solid

Yield: 52 %

The compound **(8c)** is synthesized using general procedure F, using, compound **(8b)** (1.50 g, 2.28 mmol) and $PPh_3$ (2.42 g, 9.1 mmol). The reaction mixture was extracted with DCM and purified using NPC (0.750 g, 1.18 mmol, 52 %), $R_f$ value = 0.29 in 10% MeOH/DCM. **$^1$H NMR (400 MHz, $CDCl_3$) δ:** 7.55 (s, 1H), 7.24 – 7.14 (m, 2H), 6.87 – 6.78 (m, 2H), 5.40 (s, 2H), 4.59 (s, 2H), 3.88 (t, $J$ = 6.6 Hz, 2H), 3.68 – 3.50 (m, 14H), 2.85 (t, $J$ = 5.1 Hz, 2H), 1.78 – 1.68 (m, 2H), 1.43 – 1.36 (m, 2H), 1.21 (s, 29H), 0.86 – 0.81 (m, 3H). **$^{13}$C NMR (100 MHz, $CDCl_3$) δ:** 159.49, 145.30, 144.92, 129.77, 129.70, 128.72, 126.39, 125.96, 124.37, 123.48, 122.65, 122.44, 114.97, 70.80, 70.48, 70.38, 70.30, 70.04, 69.74, 69.68, 68.12, 64.63,

64.49, 53.72, 40.84, 31.93, 31.51, 30.14, 29.70, 29.67, 29.62, 29.59, 29.41, 29.37, 29.21, 26.34, 26.04, 22.69, 14.14.**MALDI-TOF MS (M+Na):** 655.6 g/mol.

###### 2.3.1.5.16. Compound (8d)

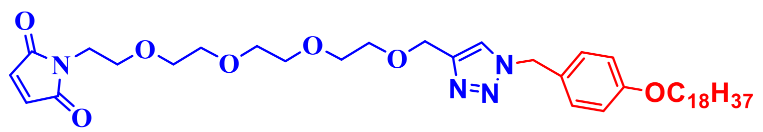


**(8d)**

Mol. formula: $C_{40}H_{64}N_4O_7$

Mol. weight: 712.97 g/mol.

Physical appearance: waxy white solid

Yield: 45 %

The compound **(8d)** is synthesized using general procedure G, using, compound **(8c)** (0.750 g, 1.18 mmol) and N-(methoxycarbonyl) maleimide (0.218 g, 1.41 mmol). Purification by NPC using MeOH/DCM (0.378 g, 0.523 mmol, 45 %), $R_f$ value = 0.33 in 10 % MeOH/DCM. **$^{1}$H NMR (400 MHz, $CDCl_3$) δ:** 7.55 (s, 1H), 7.24 – 7.14 (m, 2H), 6.87 – 6.78 (m, 2H), 5.40 (s, 2H), 4.59 (s, 2H), 3.88 (t, $J$ = 6.6 Hz, 2H), 3.68 – 3.50 (m, 14H), 2.85 (t, $J$ = 5.1 Hz, 2H), 1.78 – 1.68 (m, 2H), 1.43 – 1.36 (m, 2H), 1.21 (s, 29H), 0.86 – 0.81 (m, 3H). **$^{13}$C NMR (100 MHz, $CDCl_3$) δ:** 170.57, 159.36, 159.33, 157.65, 145.23, 145.19, 134.04, 129.57, 129.55, 126.32, 122.35, 114.84, 114.82, 114.00, 70.44, 70.36, 70.32, 69.91, 69.58, 67.98, 67.95, 67.68, 64.50, 64.47, 53.54, 53.51, 53.44, 51.96, 51.91, 37.00, 31.81, 31.57, 31.40, 31.32, 29.59, 29.55, 29.50, 29.47, 29.29, 29.25, 29.09, 28.83, 25.92, 22.58, 14.03. **MALDI-TOF MS (M+K):**751.6 g/mol.

##### 2.3.2. The general procedure of synthesis of the fluoro phosphonate probe

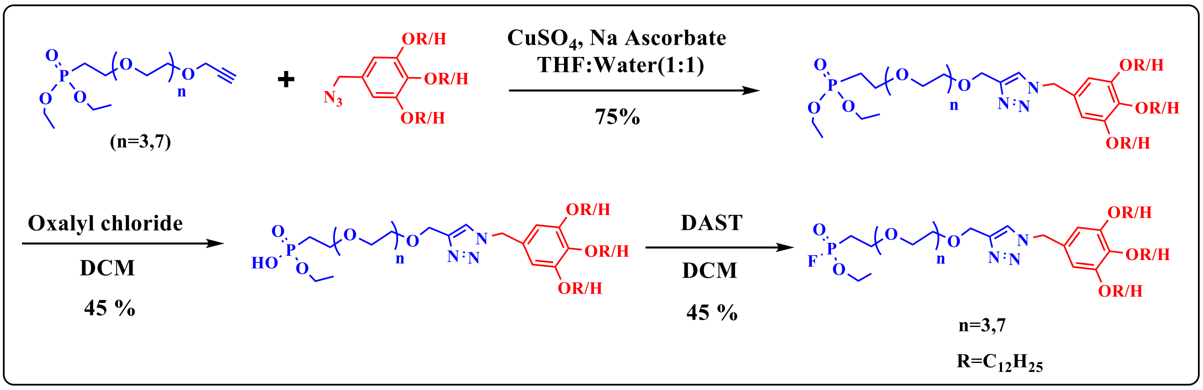


**Supplementary Scheme 8:** General scheme for the synthesis of fluorophosphonate probe.

All the fluorinated phosphonate probes were synthesized using a [2 + 3] dipolar cycloaddition (click reaction) of hydrophilic alkynes (diphosphonate esters) with hydrophobic azides. This reaction was carried out in the presence of sodium ascorbate and copper sulphate ($CuSO_4$) in a 50% mixture of THF and water. The resulting product was then treated with oxalyl chloride in DCM to obtain the monophosphonate ester, which was subsequently fluorinated using diethylaminosulfur trifluoride (DAST) in DCM. The detailed procedure is outlined below.

##### 2.3.2.1. Synthesis of click product-procedure H

Hydrophobic azide (4 eq) and hydrophilic alkyne (1 eq) were dissolved in degassed THF and stirred until a clear solution was obtained. Degassed $H_2O$ was added and stirred vigorously for an additional 10 minutes. Freshly prepared 1M Na ascorbate (0.05 eq) and 1M $CuSO_4$ (0.1 eq) were added to the reaction mixture thrice in intervals of 45 mins and allowed to react for 16 hrs at RT. Upon completion, the reaction mixture was extracted with DCM, and the combined organic layer was dried over $Na_2SO_4$ and concentrated under reduced pressure to obtain the crude product, which was then purified using NPC with a MeOH/DCM solvent system.

##### 2.3.2.2. Synthesis of monophosphonate ester - Procedure I

Diphosphonate ester (1 eq) was dissolved in DCM with stirring. Then, OxCl (4 eq) was added dropwise at RT and allowed to react for 18 hrs under stirring. Upon completion, excess of OxCl and DCM were removed under vacuum. Then, water was added to the residue and stirred for 5 minutes. The resulting mixture was extracted thrice with DCM, and the combined organic layer was dried over $Na_2SO_4$ and concentrated under a vacuum to get the crude product, which was used for the next step without further purification.

##### 2.3.2.3. Synthesis of fluorophosphonate - Procedure J

To the stirring solution of monophosphonate ester (1 eq) in DCM, DAST (4 eq) was added dropwise at RT and allowed to react for 4 hrs. Excess of DAST and DCM were evaporated under reduced pressure. To the obtained residue, water was added and stirred for an additional 2 minutes to quench any residual DAST. The reaction mixture was then extracted thrice with DCM. The combined organic layer was dried over $Na_2SO_4$ and concentrated under vacuum to get the crude product. This final probe was used for protein modification without further purification.

**2.3.2.4. Synthesis of fluoro phosphonate probe and its intermediates**

**2.3.2.4.1. Compound (9c)**

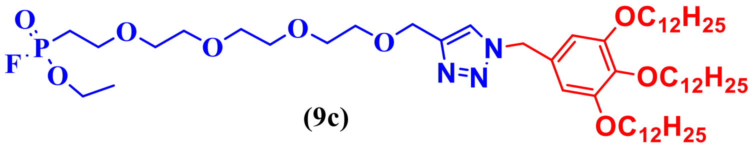


**(9c)**

The compound 9c and its intermediates are synthesized using a previously reported procedure[1].

**2.3.2.4.2. Compound (10c)**

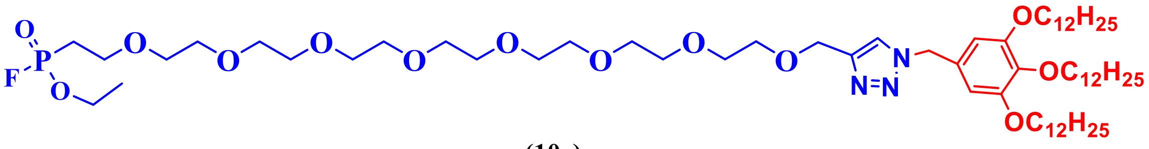


**(10c)**

The compound 10c and its intermediates are synthesized using a previously reported procedure[1].

**2.3.2.4.3. Compound (11a)**

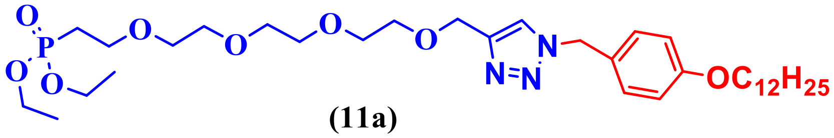


**(11a)**

Mol. formula: $C_{34}H_{60}N_3O_8P$

Mol. weight: 669.64 g/mol

Physical appearance: brownish liquid

Yield: 62 %

The compound **(11a)** was synthesized using general procedure H, from compound **(1g)** (0.240 g, 0.681 mmol) and compound **(3c)** (0.864 g, 2.72 mmol), $CuSO_4$ (17 mg, 0.068 mmol), Na ascorbate (7 mg, 0.034 mmol) in 50 % THF / $H_2O$. The crude product obtained was purified using NPC with MeOH/DCM to yield a brownish liquid (0.282 g, 0.422 mmol, 62%), with an $R_f$ value of 0.40 in 5% MeOH/DCM.**$^1$H NMR (400 MHz, $CDCl_3$) δ:** 7.44 (s, 1H), 7.17 (d, *J* = 8.3 Hz, 2H), 6.83 (d, *J* = 8.5 Hz, 2H), 5.39 (s, 2H), 4.60 (s, 2H), 4.06 (dt, *J* = 11.0, 7.2, 4.0, 3.4 Hz, 4H), 3.90 (t, *J* = 6.6 Hz, 2H), 3.74 – 3.52 (m, 15H), 2.08 (dt, *J* = 18.7, 7.5 Hz, 2H), 1.74 (d, *J* = 7.2 Hz, 2H), 1.40 (dq, *J* = 14.2, 6.5 Hz, 2H), 1.27 (p, *J* = 6.6 Hz, 19H), 0.84 (t, *J* = 6.7 Hz, 3H). **$^{13}$C NMR (100 MHz, $CDCl_3$) δ:** 159.52, 129.71, 126.32, 115.01, 70.57, 70.53, 70.48, 70.44, 70.18, 69.70, 68.14, 65.11, 64.68, 61.72, 61.66, 53.75, 31.92, 29.71, 29.67, 29.64, 29.60, 29.58, 29.39, 29.35, 29.21, 27.64, 26.25, 26.03, 22.70, 16.47, 16.41, 14.14. **MALDI-TOF MS (M+Na):** 692.52 g/mol.

##### 2.3.2.4.4. Compound (11b)

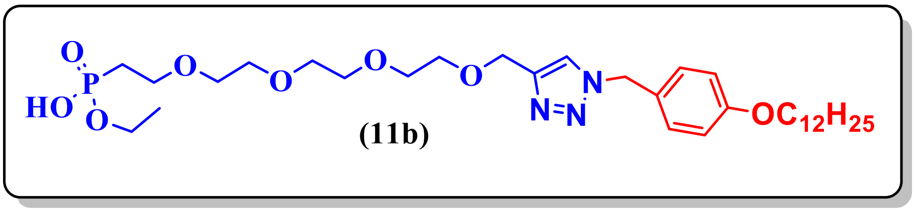


Mol. formula: $C_{32}H_{56}N_3O_8P$

Mol. weight: 641.49 g/mol

Physical appearance: pale yellow liquid

Yield: not determined

The compound **(11b)** was synthesized using the general procedure I. From compound **(11a)** (0.250 g, 0.373 mmol), OxCl (0.189 g, 1.49 mmol) in DCM. The product was carried to the next step without purification. **MALDI-TOF MS (M+Na):**664.51 g/mol.

##### 2.3.2.4.5. Compound (11c)

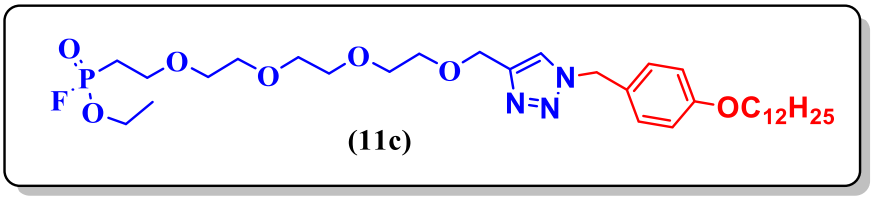


(11c)

Mol. formula: $C_{32}H_{55}FN_3O_7P$

Mol. weight: 643.78 g/mol

Physical appearance: brownish liquid

Yield: not determined

The compound **(11c)** was synthesized using general procedure J, from compound **(11b)** (0.170 g, 0.265 mmol), and DAST (0.170 g, 1.06 mmol) in DCM. The product was used for modification without further purification. **$^{19}$F NMR (377 MHz, $CDCl_3$) δ:** -59.98, -62.81. **MALDI-TOF MS (M+Na):**666.97 g/mol.

##### 2.3.3. The general procedure of synthesis of the N-terminal probe

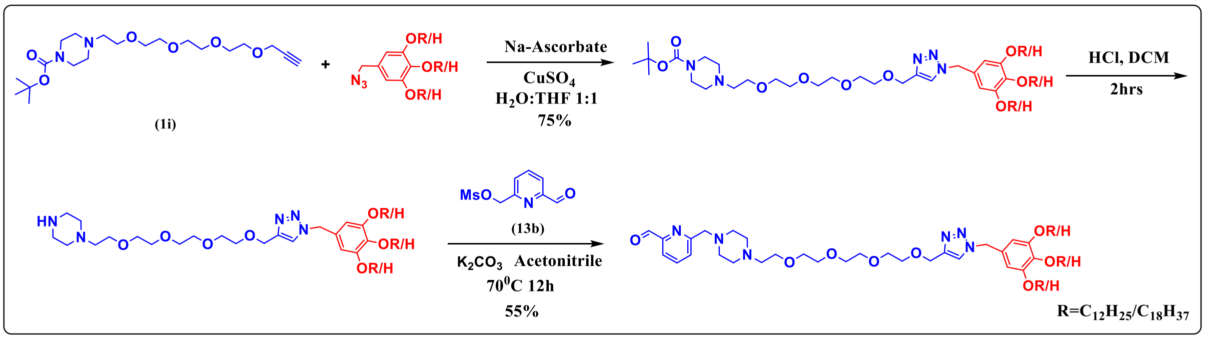


**Supplementary Scheme 9:** General scheme for the synthesis of N-terminal probe.

All the N-terminal probes were synthesized using a [2 + 3] dipolar cycloaddition (click reaction) of hydrophilic alkynes with hydrophobic azides. This reaction was carried out in the presence of sodium ascorbate and copper sulphate ($CuSO_4$) in a 50% mixture of THF and water. The clicked product was treated with concentrated HCl to deprotect the amine, followed by a reaction with a pyridine derivative to obtain the final probe. The detailed procedure is given below:

###### 2.3.3.1. Synthesis of click product- procedure K

Hydrophilic alkyne (1i) (1 eq) and hydrophobic azide (4 eq) were dissolved in degassed THF and stirred until a clear solution was obtained. Then, degassed $H_2O$ was added and stirred vigorously for 10 more minutes. Freshly prepared 1M Na ascorbate (0.05 eq) and 1M $CuSO_4$ (0.1 eq) were added to the reaction mixture thrice in intervals of 45 mins and allowed to react for 16 hrs at RT. Upon completion, the reaction mixture was extracted with DCM, and the combined organic layer was dried over $Na_2SO_4$ and concentrated under reduced pressure to obtain the crude product, which was then purified by NPC using a MeOH/DCM solvent system to yield the clicked product.

###### 2.3.3.2. Synthesis of click product- procedure L

The clicked compound dissolved in DCM, and 2 mL Conc HCl was added to the reaction mixture and stirred for two hrs. After two hrs reaction mixture was quenched by the addition of aq. NaOH solution and extracted with DCM. The organic layer is concentrated and crude used as it is in the next step.

###### 2.3.3.3. Synthesis of click product- procedure M

The crude amine (1eq), mesylate (13b) (4 eq), and $K_2CO_3$ (4 eq) were weighed in RBF. Then the mixture was dissolved in ACN and refluxed at 65 □C. After 16 hrs, the reaction mixture was concentrated and purified by NPC using MeOH/DCM to get the final compound.

###### 2.3.3.4. Synthesis of N-terminal probe and its intermediates

###### 2.3.3.4.1. Compound (14a)

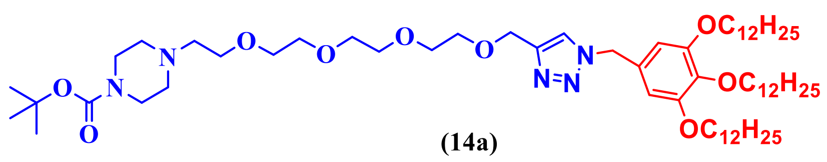


**(14a)**

Mol. formula: $C_{63}H_{115}N_5O_9$

Mol. weight: 1086.64 g/mol

Physical appearance: brownish liquid

Yield: 65 %

The compound **(14a)** was synthesized using general procedure K, from compound **(1i)** (1.5 g, 3.75 mmol) and compound **(2e)** (10.2 g, 15 mmol), $CuSO_4$ (93 mg, 0.375 mmol), Na ascorbate (37 mg, 0.187mmol) in 50 % THF / $H_2O$. The crude product obtained was purified using NPC using MeOH / DCM to get a brownish liquid (2.64 g, 2.43 mmol, 65 %), $R_f$ value

= 0.40 in 5 % MeOH / DCM. **$^{1}$H NMR (400 MHz, $CDCl_3$) δ:** 7.47 (s, 1H), 6.44 (s, 2H), 5.36 (s, 2H), 4.64 (s, 2H), 3.90 (q, *J* = 6.2 Hz, 6H), 3.69 – 3.55 (m, 14H), 3.42 (t, *J* = 5.1 Hz, 4H), 2.58 (t, *J* = 5.7 Hz, 2H), 2.44 (t, *J* = 5.1 Hz, 4H), 1.79 – 1.66 (m, 6H), 1.49 – 1.36 (m, 15H), 1.25 (d, *J* = 8.0 Hz, 49H), 0.86 (t, *J* = 6.7 Hz, 9H). **$^{13}$C NMR (100 MHz, $CDCl_3$) δ:** 154.79, 153.65, 145.53, 138.56, 129.46, 122.53, 114.15, 106.85, 79.69, 73.54, 70.63, 70.58, 70.43, 69.84, 69.32, 68.74, 64.80, 57.87, 54.55, 53.41, 33.90, 32.02, 32.00, 30.40, 29.83, 29.81, 29.79, 29.77, 29.74, 29.72, 29.68, 29.50, 29.47, 29.45, 29.03, 28.50, 26.19, 26.17, 22.77, 14.20, 0.08.**MALDI-TOF MS (M+Na):**1109.69 g/mol.

##### 2.3.3.4.2. Compound (14b)

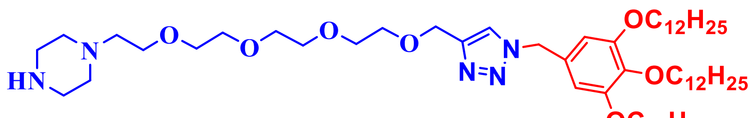


**(14b)**

Mol. formula: $C_{58}H_{107}N_5O_7$

Mol. weight: 1037.78 g/mol

Physical appearance: brownish liquid

Yield: Not determined

The compound **(14b)** was synthesized using general procedure L, from the clicked product **(14a)** and Conc HCl. The crude is used as it is in the next step. **MALDI-TOF MS (M+Na):**1060.87 g/mol.

##### 2.3.3.4.3. Compound (14c)

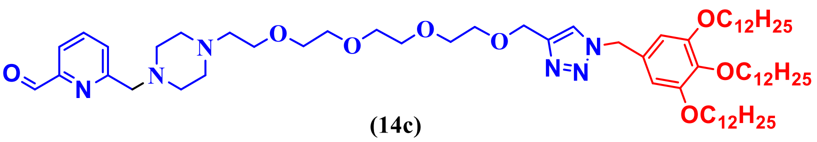


**(14c)**

Mol. formula: $C_{65}H_{112}N_6O_8$

Mol. weight: 1105.65 g/mol

Physical appearance: brownish liquid

Yield: 35 %

The compound **(14c)** was synthesized using general procedure M, using crude compound **(14b)** (1.6 g, 1.54 mmol), $K_2CO_3$ (0.837 g, 6.16 mmol), and mesylate compound **(13b)** (1.32 g, 6.16 mmol). The reaction mixture was purified by NPC using a MeOH/DCM eluent system (0.592 g, 0.539 mmol, 35%), with an $R_f$ value of 0.33 in 10% MeOH/DCM. **$^1$H NMR (400 MHz, $CDCl_3$) δ:** 10.03 (s, 1H), 7.80 (d, *J* = 4.5 Hz, 2H), 7.65 (q, *J* = 4.2 Hz, 1H), 7.47 (s, 1H), 6.43 (s, 2H), 5.35 (s, 2H), 4.62 (s, 2H), 3.88 (q, *J* = 6.3 Hz, 6H), 3.73 (s, 2H), 3.68 – 3.52 (m, 14H), 2.58 (dd, *J* = 11.0, 5.0 Hz, 9H), 1.80 – 1.63 (m, 6H), 1.41 (ddd, *J* = 16.1, 10.4, 6.5 Hz, 6H), 1.29 – 1.21 (m, 48H), 0.84 (t, *J* = 6.7 Hz, 10H). **$^{13}$C NMR (100 MHz, $CDCl_3$) δ:** 193.66, 159.70, 153.60, 152.38, 145.46, 139.26, 138.50, 137.38, 129.43, 127.49, 122.51, 120.21, 114.11, 106.79, 73.48, 70.57, 70.52, 70.36, 69.78, 69.26, 68.74, 64.74, 64.00, 57.73, 54.50, 53.51, 53.08, 33.85, 31.97, 31.96, 31.71, 31.54, 31.46, 30.36, 30.16, 29.78, 29.76, 29.74, 29.69, 29.67, 29.63, 29.54, 29.45, 29.42, 29.40, 29.18, 28.98, 26.15, 26.12, 22.72, 14.16.**MALDI-TOF MS (M+Na):**1128.79 g/mol.

##### 2.3.3.4.4. Compound (15a)

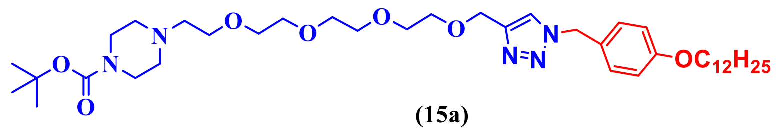

**(15a)**

Mol. formula: $C_{39}H_{67}N_5O_7$

Mol. weight: 717.99 g/mol

Physical appearance: brownish liquid

Yield: 65 %

The compound **(15a)** was synthesized using general procedure K, from compound **(1i)** (0.540 g, 1.34 mmol) and compound **(3c)** (1.7 g, 5.39 mmol), $CuSO_4$ (33 mg, 0.134 mmol), Na ascorbate (13 mg, 0.067 mmol) in 50 % THF / $H_2O$. The crude product obtained was purified using NPC using MeOH / DCM to get a brownish liquid (0.625 g, 0.871 mmol, 65 %), $R_f$ value = 0.32 in 10 % MeOH / DCM. **$^1$H NMR (400 MHz, $CDCl_3$) δ:** 7.41 (s, 1H), 7.19 – 7.09 (m, 2H), 6.81 (dd, *J* = 9.0, 2.3 Hz, 2H), 5.37 (s, 2H), 4.58 (d, *J* = 2.0 Hz, 2H), 3.87 (t, *J* = 6.6 Hz, 2H), 3.62 – 3.48 (m, 14H), 3.37 (t, *J* = 5.1 Hz, 4H), 2.53 (t, *J* = 5.7 Hz, 2H), 2.38 (t, *J* = 5.1 Hz, 4H), 1.70 (dt, *J* = 12.9, 6.6 Hz, 3H), 1.39 (d, *J* = 2.3 Hz, 12H), 1.26 – 1.17 (m, 16H), 0.86 – 0.76 (m, 3H). **$^{13}$C NMR (100 MHz, $CDCl_3$) δ:** 159.44, 154.65, 145.35, 129.62, 126.28, 122.26, 114.92, 79.48, 70.51, 70.49, 70.44, 70.30, 69.64, 68.70, 68.67, 68.04, 64.63, 57.75, 53.63, 53.28, 33.76, 31.85, 31.65, 31.46, 30.09, 29.63, 29.59, 29.57, 29.53, 29.51, 29.32, 29.28, 29.14, 28.88, 28.37, 25.96, 22.62, 14.08.**MALDI-TOF MS (M+Na):**740.64 g/mol.

##### 2.3.3.4.5. Compound (15b)

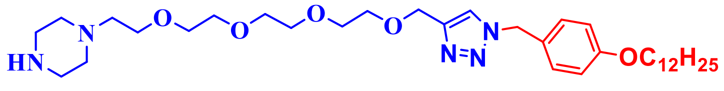


**(15b)**

Mol. formula: $C_{34}H_{59}N_5O_5$

Mol. weight: 617.88 g/mol

Physical appearance: brownish liquid

Yield: Not determined

The compound **(15b)** was synthesized using general procedure L, using the clicked product **(15a)**, and Conc HCl. The crude is used as it is in the next step. **MALDI-TOF MS (M+Na):**640.65 g/mol.

##### 2.3.3.4.6. Compound (15c)

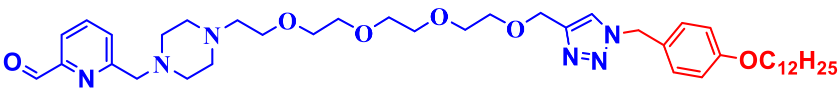


**(15c)**

Mol. formula: $C_{41}H_{64}N_6O_6$

Mol. weight: 717.99 g/mol

Physical appearance: brownish liquid

Yield: 40 %

The compound **(15c)** was synthesized using general procedure M, using Crude Compound **(15b)** (0.263 g, 0.427 mmol), $K_2CO_3$ (0.231 g, 1.70 mmol), and mesylate compound **(13b)** (0.365 g, 1.70 mmol). The reaction mixture was purified by NPC using MeOH/DCM as an eluting system (0.122 g,0.170 mmol, 40 %), $R_f$ value = 0.32 in 10 % MeOH / DCM. **$^1$H NMR (400 MHz, CDCl$_3$) δ:** 10.00 (s, 1H), 7.85 – 7.76 (m, 1H), 7.46 (d, *J* = 2.0 Hz, 1H), 7.21 – 7.10 (m, 2H), 6.86 – 6.75 (m, 2H), 5.38 (s, 2H), 4.57 (s, 2H), 3.88 (t, *J* = 6.5 Hz, 2H), 3.72 (s, 2H), 3.65 – 3.51 (m, 14H), 2.98 (s, 7H), 2.62 (d, *J* = 5.6 Hz, 4H), 1.71 (p, *J* = 6.7 Hz, 2H), 1.42 – 1.35 (m, 2H), 1.21 (d, *J* = 6.6 Hz, 16H), 0.82 (t, *J* = 6.7 Hz, 3H). **$^{13}$C NMR (100 MHz, CDCl$_3$) δ:** 193.59, 159.49, 159.24, 152.29, 145.21, 137.71, 137.55, 129.69, 127.70, 126.23, 123.50, 122.55, 122.52, 120.40, 119.98, 114.99, 114.05, 96.38, 70.44, 70.41, 70.38, 70.22, 70.20, 69.65, 68.17, 68.13, 64.49, 63.72, 63.69, 57.53, 57.46, 54.67, 53.74, 53.26, 53.05, 52.60, 52.43, 50.29, 49.14, 33.79, 31.88, 31.64, 29.66, 29.62, 29.60, 29.56, 29.54, 29.35, 29.31, 29.16, 28.92, 25.99, 22.65, 14.09.**MALDI-TOF MS (M+Na):** 740.44 g/mol.

##### 2.3.3.4.7. Compound (16a)

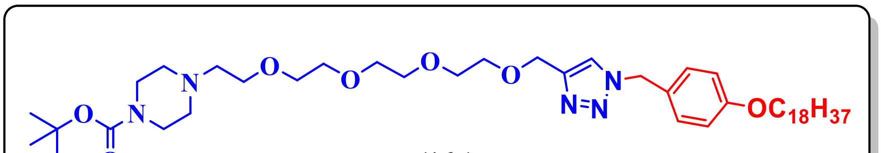


**(16a)**

Mol. formula: $C_{45}H_{79}N_5O_7$

Mol. weight: 802.16 g/mol

Physical appearance: White solid

Yield: 65 %

The compound **(16a)** was synthesized using general procedure K, from compound **(1i)** (0.600, 1.49 mmol) and compound **(4c)** (2.4 g, 5.99 mmol), $CuSO_4$ (37 mg, 0.149 mmol), Na ascorbate (1 mg, 0.074 mmol) in 50 % THF / $H_2O$. The crude product obtained was purified using NPC with MeOH/DCM to yield a brownish liquid (0.776 g, 0.968 mmol, 65%), with an $R_f$ value of 0.40 in 10% MeOH/DCM. **$^1$H NMR (400 MHz, $CDCl_3$) δ:** 7.43 (s, 1H), 7.23 – 7.14 (m, 2H), 6.90 – 6.80 (m, 2H), 5.41 (s, 2H), 4.62 (s, 2H), 3.91 (t, $J$ = 6.6 Hz, 2H), 3.69 – 3.54 (m, 14H), 3.41 (t, $J$ = 5.0 Hz, 4H), 2.56 (t, $J$ = 5.8 Hz, 2H), 2.41 (t, $J$ = 5.1 Hz, 4H), 1.79 – 1.71 (m, 2H), 1.43 (s, 12H), 1.23 (s, 31H), 0.85 (t, $J$ = 6.7 Hz, 3H). **$^{13}$C NMR (100 MHz, $CDCl_3$) δ:** 159.61, 154.83, 129.78, 126.36, 122.40, 115.08, 79.67, 70.67, 70.65, 70.63, 70.59, 70.45, 69.79, 68.84, 68.22, 64.79, 57.89, 53.83, 53.42, 32.01, 31.59, 29.78, 29.75, 29.69, 29.66, 29.48, 29.45, 29.29, 28.52, 26.11, 22.78, 14.22, 0.09. **MALDI-TOF MS (M+Na):** 825.8 g/mol.

**2.3.3.4.8. Compound (16b)**

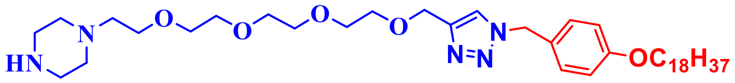


**(16b)**

Mol. formula: $C_{40}H_{71}N_5O_5$

Mol. weight: 702.04 g/mol

Physical appearance: White solid

Yield: 62 %

The compound **(16b)** was synthesized using general procedure L, using the clicked product **(16a)** and Conc HCl. The crude is used as it is in the next step. **MALDI-TOF MS (M+Na):** 725.98 g/mol.

**2.3.3.4.9. Compound (16c)**

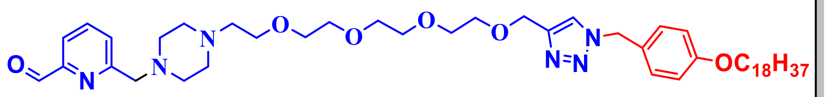

**(16c)**

Mol. formula: $C_{47}H_{76}N_6O_6$

Mol. weight: 821.16 g/mol

Physical appearance: White solid

Yield: 35 %

The compound **(16c)** was synthesized using general procedure M, using crude compound **(16b)** (1.3 g, 1.85 mmol), $K_2CO_3$ (1.00 g, 7.4 mmol), and mesylate compound **(13b)** (1.59 g, 7.4 mmol). The reaction mixture was purified by NPC using MeOH/DCM as the eluting system (0.531 g, 0.647 mmol, 35%), with an $R_f$ value of 0.39 in 10% MeOH/DCM. **$^1$H NMR (400 MHz, $CDCl_3$) δ:** 10.02 (s, 1H), 7.79 (d, $J$ = 4.5 Hz, 2H), 7.64 (q, $J$ = 4.2 Hz, 1H), 7.43 (s, 1H), 7.21 – 7.12 (m, 2H), 6.88 – 6.77 (m, 2H), 5.39 (s, 2H), 4.59 (s, 2H), 3.89 (t, $J$ = 6.6 Hz, 2H), 3.73 (s, 2H), 3.61 (dd, $J$ = 5.6, 2.7 Hz, 2H), 3.57 (td, $J$ = 5.4, 2.9 Hz, 11H), 2.59 (t, $J$ = 5.8 Hz, 8H), 1.83 – 1.64 (m, 2H), 1.44 – 1.33 (m, 3H), 1.21 (s, 29H), 0.83 (t, $J$ = 6.7 Hz, 3H). **$^{13}$C NMR (100 MHz, $CDCl_3$) δ:** 193.64, 159.56, 159.51, 152.35, 145.36, 137.42, 129.70, 127.54, 126.29, 123.33, 122.37, 121.53, 120.25, 119.93, 115.00, 70.52, 70.50, 70.47, 70.31, 69.69, 68.54, 68.13, 64.65, 63.91, 57.67, 54.68, 53.73, 53.43, 52.92, 52.76, 52.63, 51.68, 50.46, 49.50, 33.82, 31.93, 31.69, 31.51, 30.14, 29.70, 29.66, 29.61, 29.58, 29.40, 29.36, 29.20, 29.16, 28.95, 26.03, 22.69, 14.14.**MALDI-TOF MS (M+Na):**844.79 g/mol.

#### References


(1) Sandanaraj, B. S.; Reddy, M. M.; Bhandari, P. J.; Kumar, S.; Aswal, V. K. Rational Design of Supramolecular Dynamic Protein Assemblies by Using a Micelle-Assisted Activity-Based Protein-Labeling Technology. *Chemistry – A European Journal* **2018**, *24* (60), 16085–16096. https://doi.org/10.1002/chem.201802824.

**Compound (5a)**

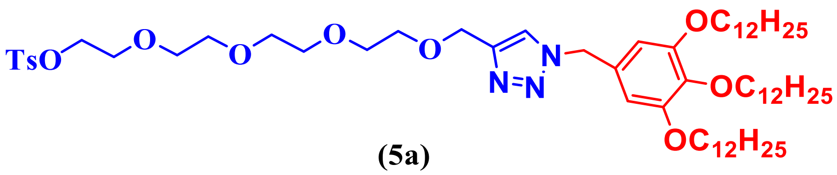


**(5a)**

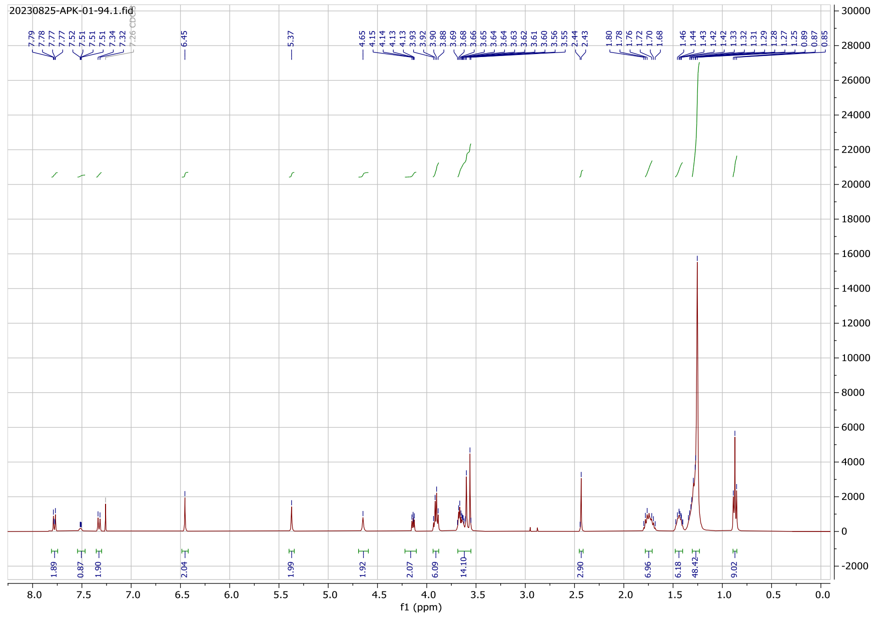

#### $^1$H NMR Spectrum

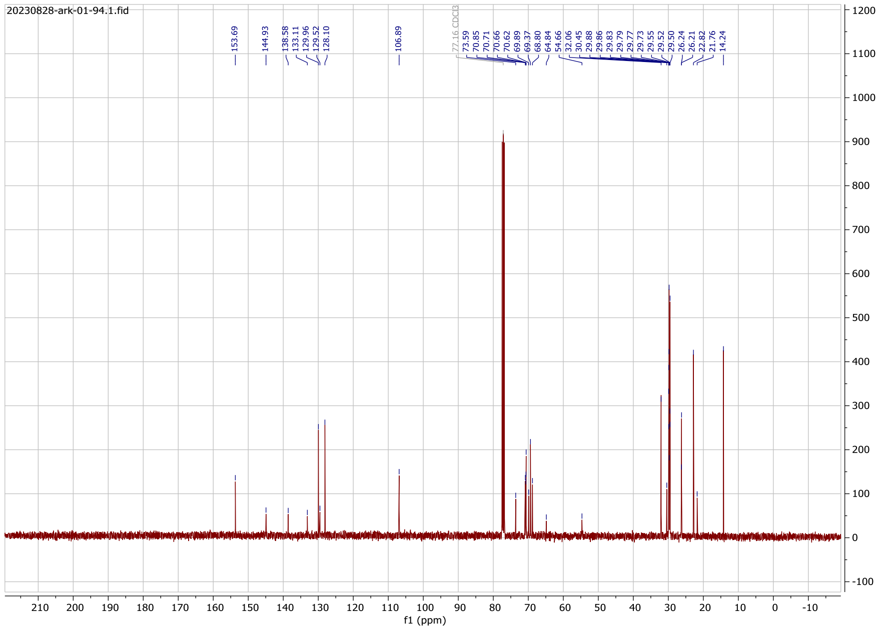

#### $^{13}$C NMR Spectrum

#### Compound (5b)

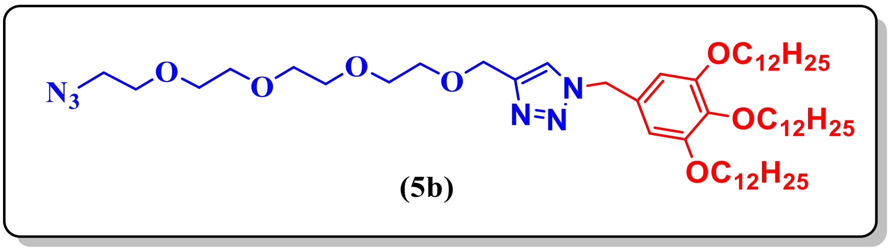

(5b)

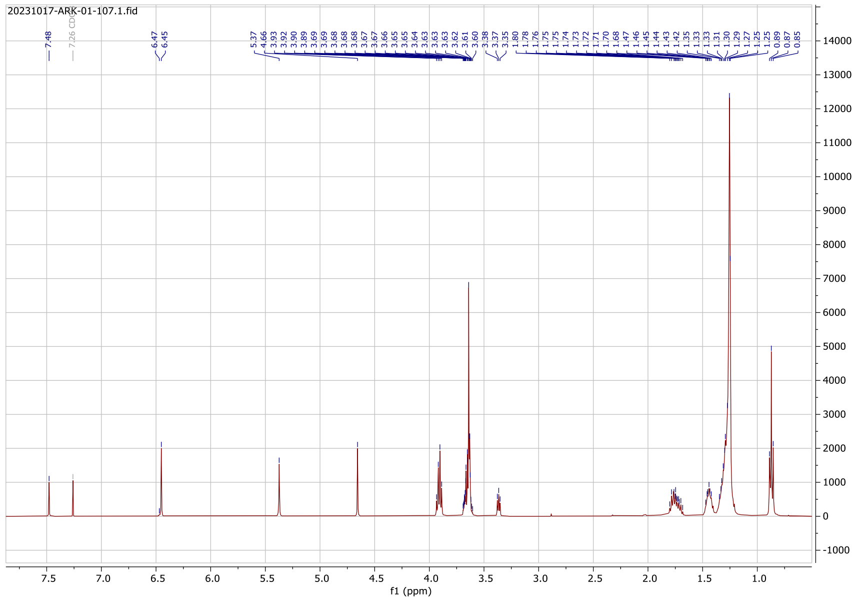

#### $^1$H NMR Spectrum

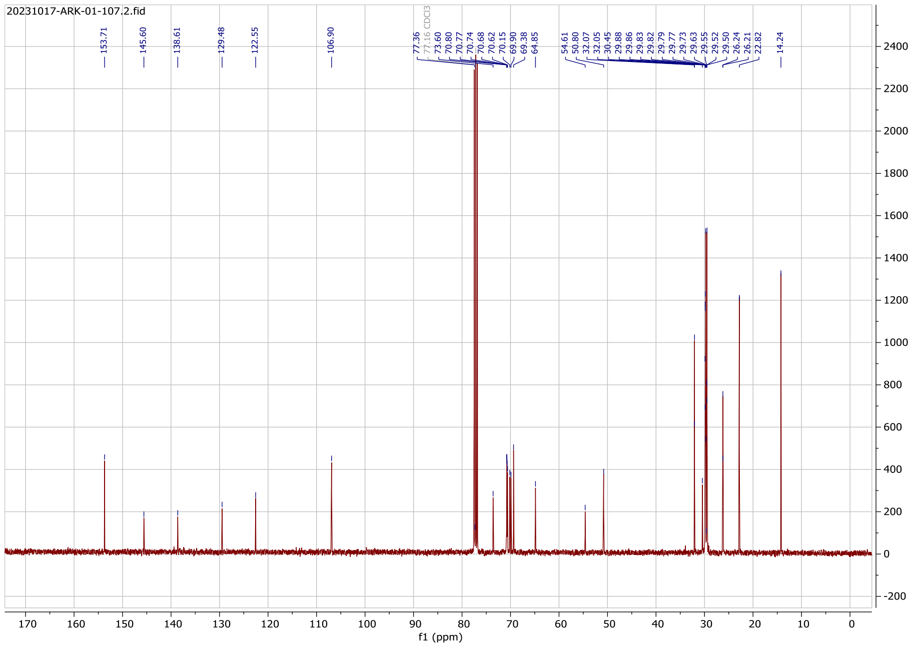

#### $^{13}$C NMR Spectrum

#### Compound (5c)

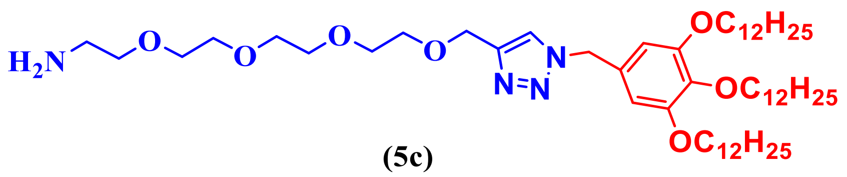

(5c)

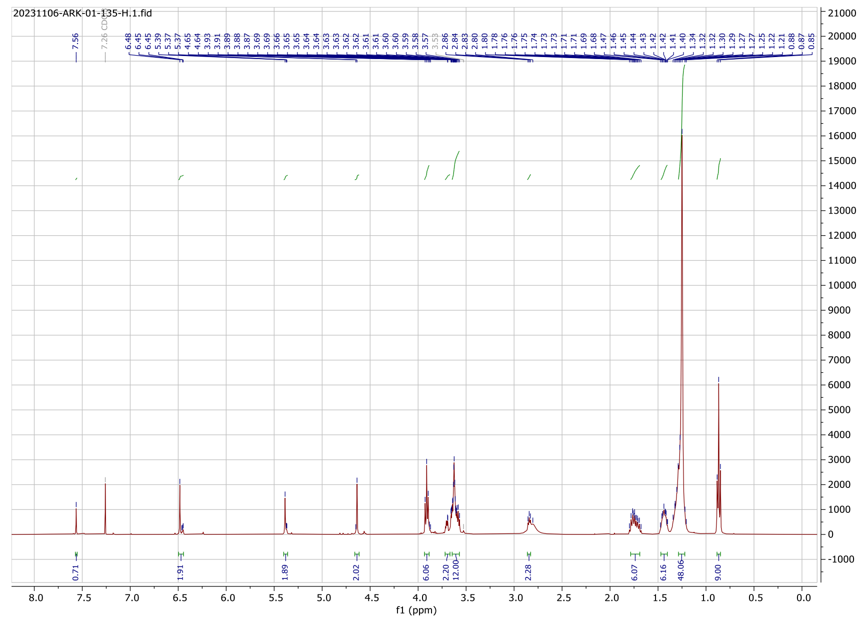

### [1]H NMR Spectrum

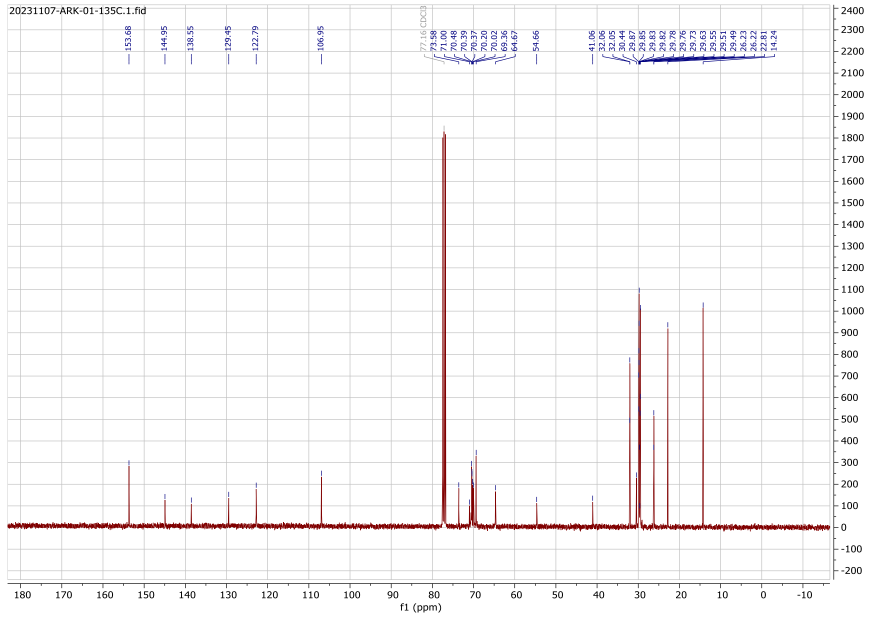

### [13]C NMR Spectrum

### Compound (5d)

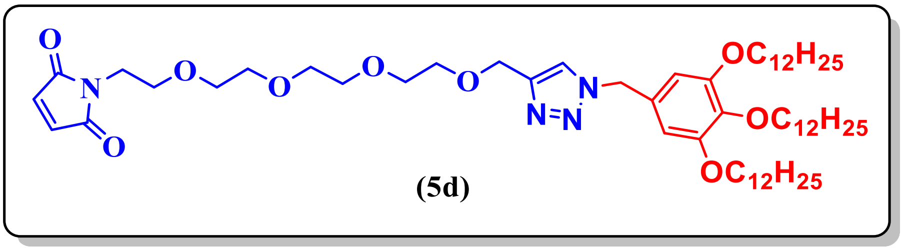

(5d)

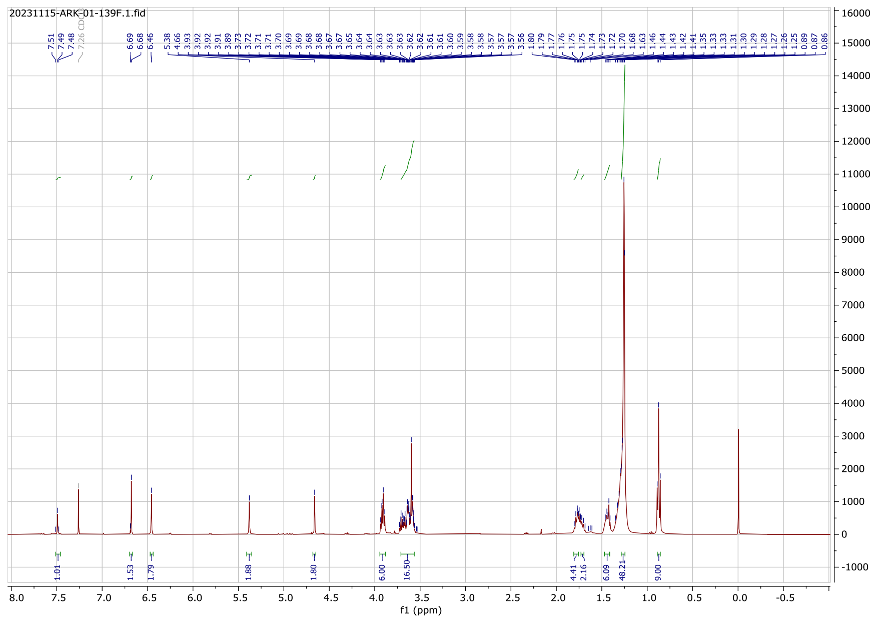

### [1]H NMR Spectrum

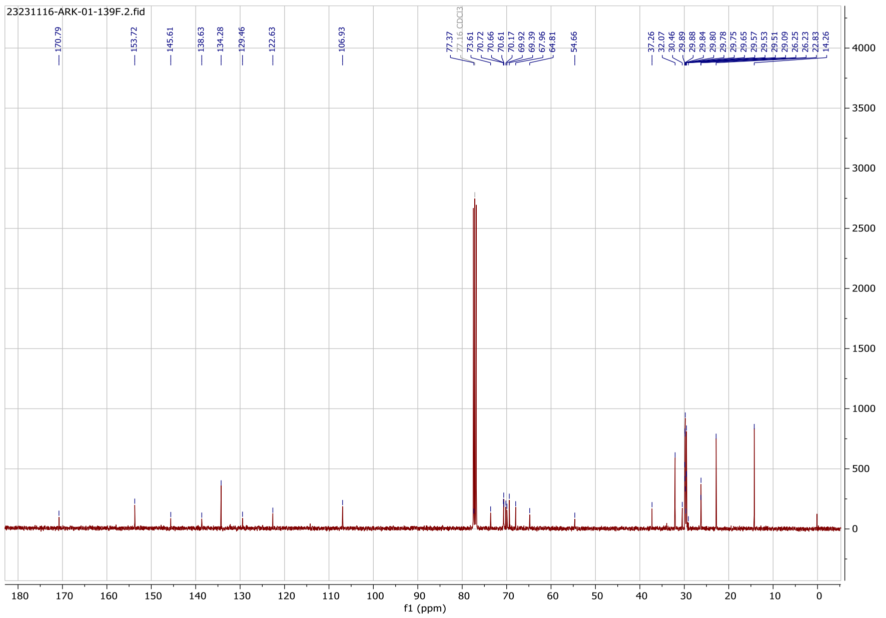

### [13]C NMR Spectrum

### Compound (6a)

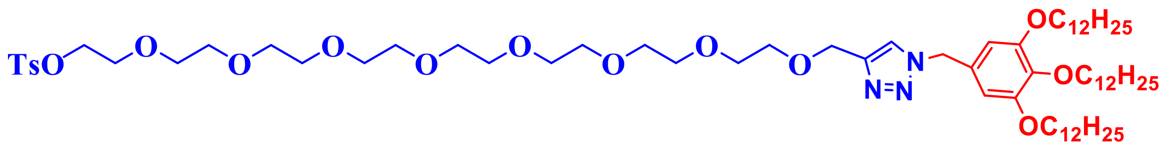


**(6a)**

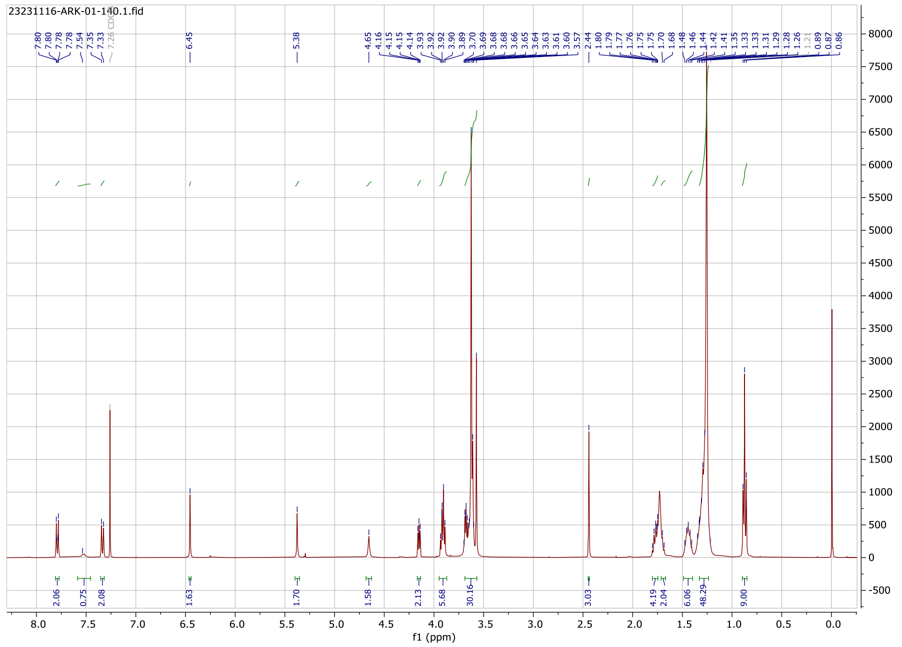


#### $^{1}$H NMR Spectrum

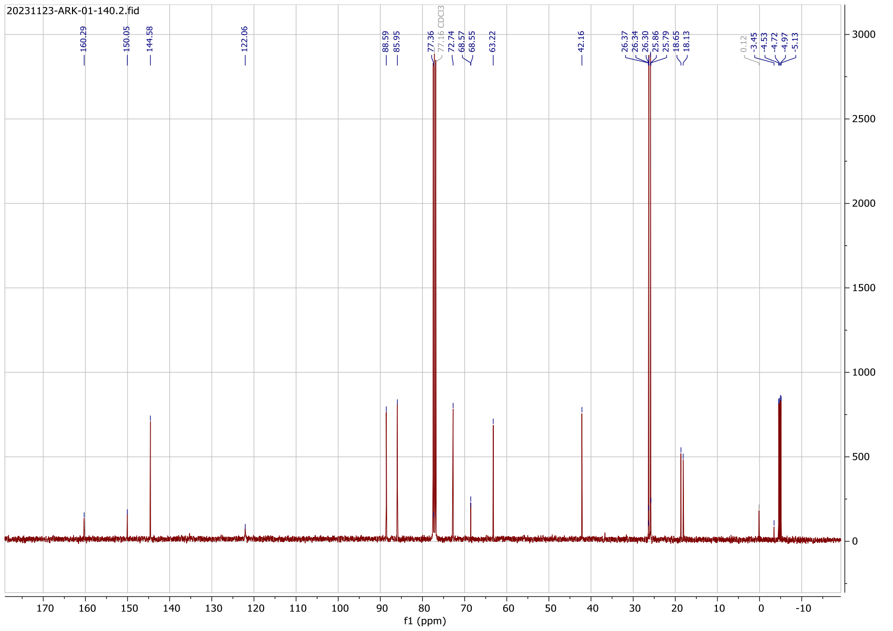


#### $^{13}$C NMR Spectrum

### Compound (6b)

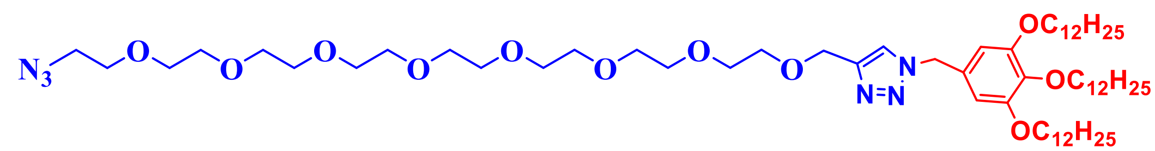


(6b)

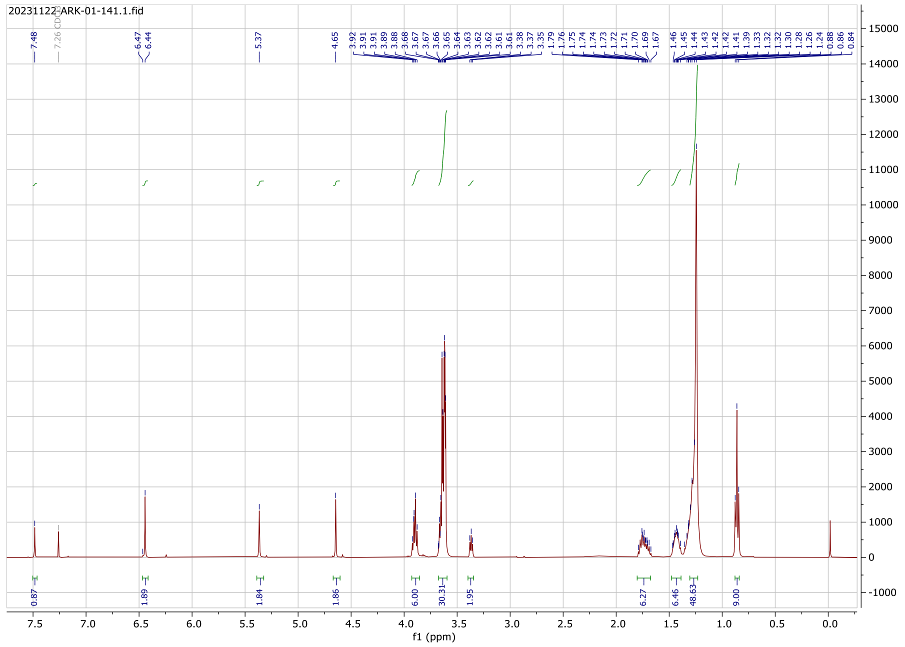

#### [1]H NMR Spectrum

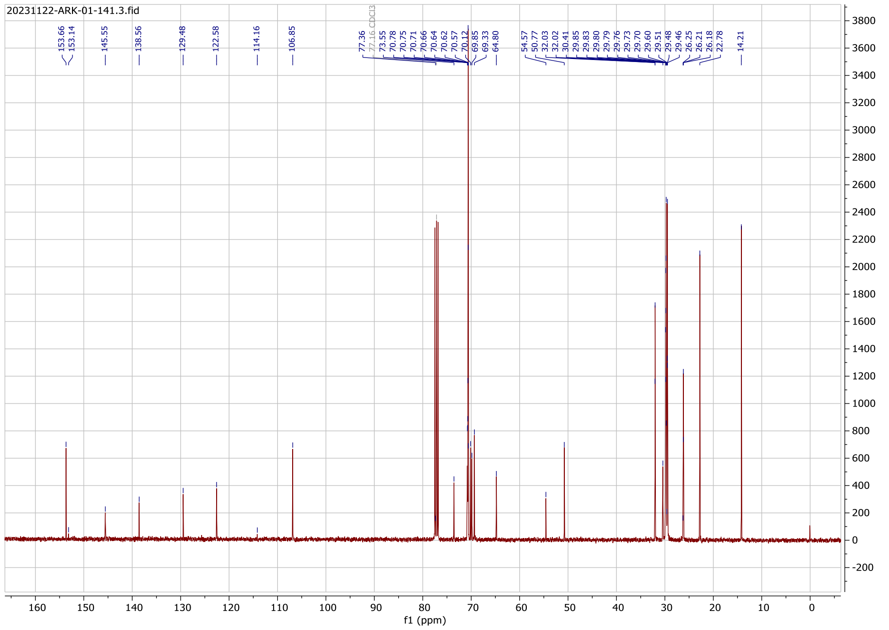

#### $^{13}$C NMR Spectrum

#### Compound (6c)

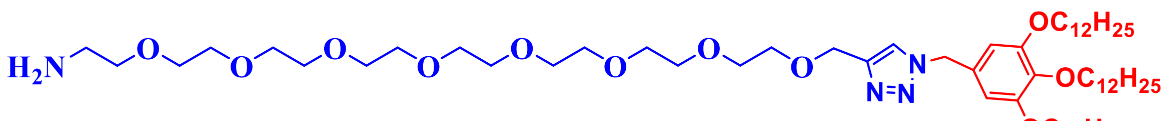

**(6c)**

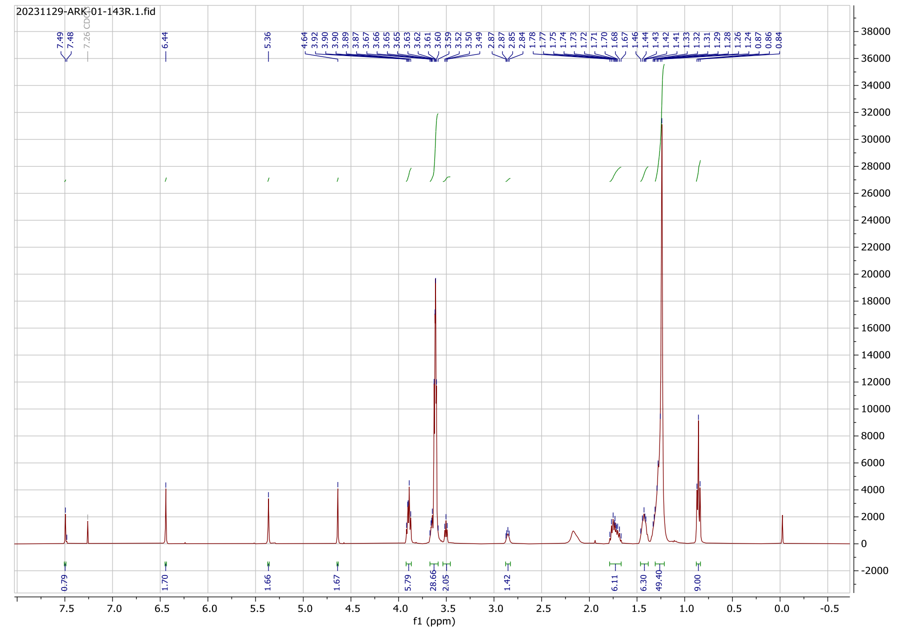

#### $^{1}$H NMR Spectrum

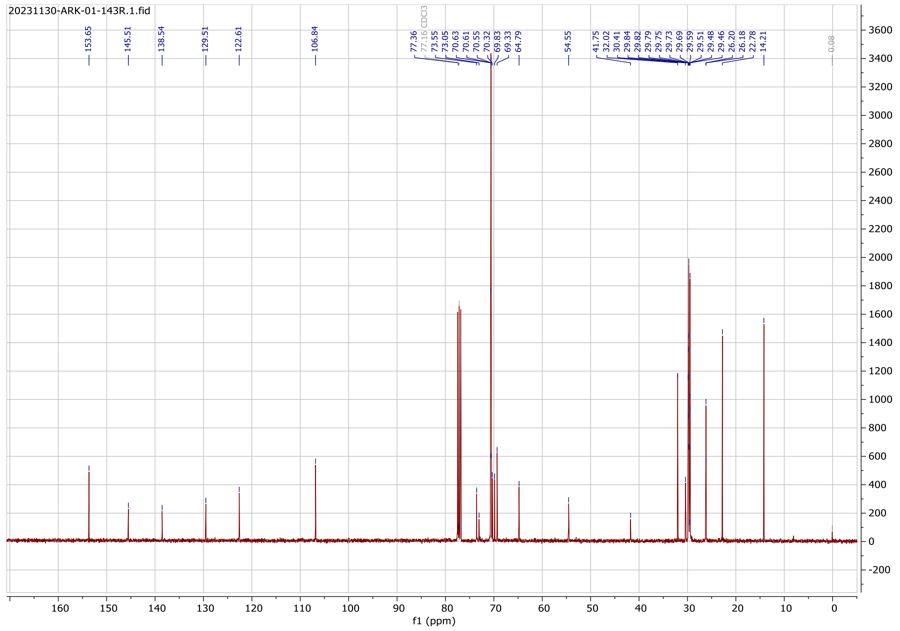

#### $^{13}$C NMR Spectrum

#### Compound (6d)

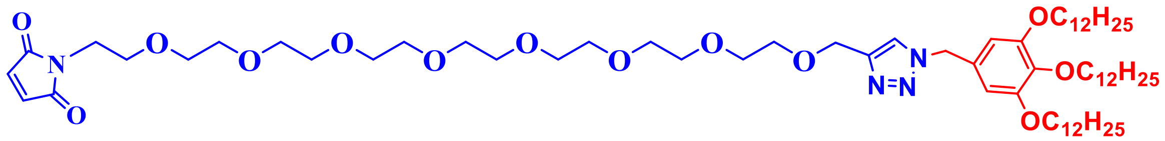


(6d)

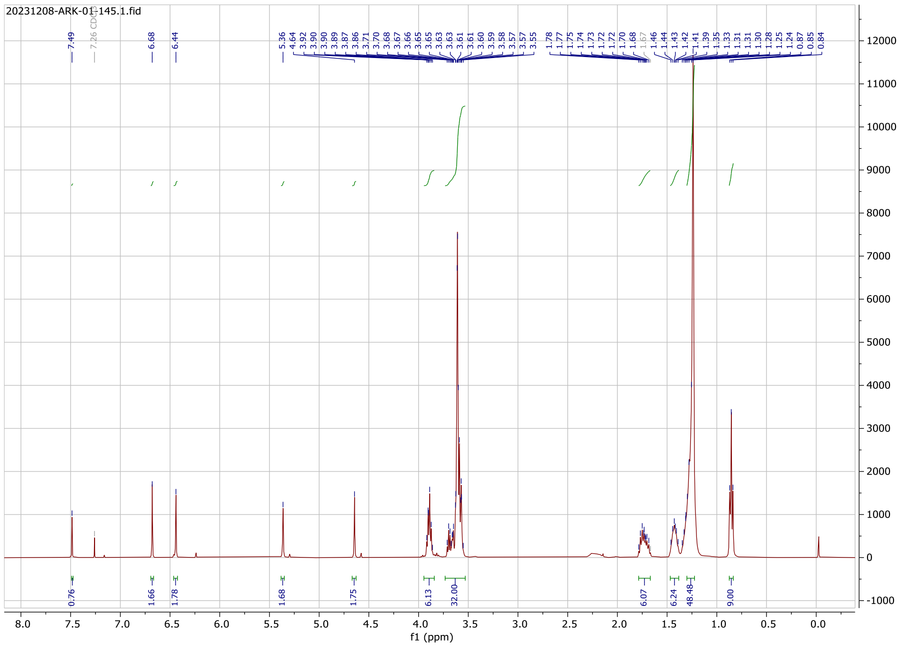


#### $^{1}$H NMR Spectrum

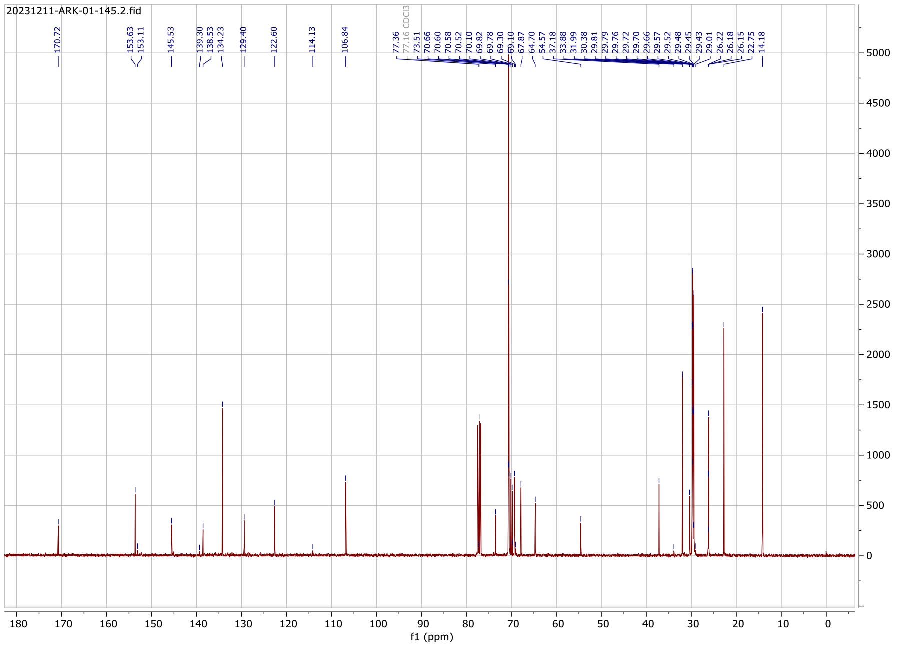

#### $^{13}$C NMR Spectrum

#### Compound (7a)

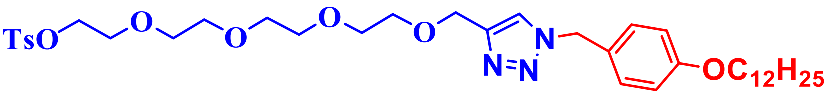


(7a)

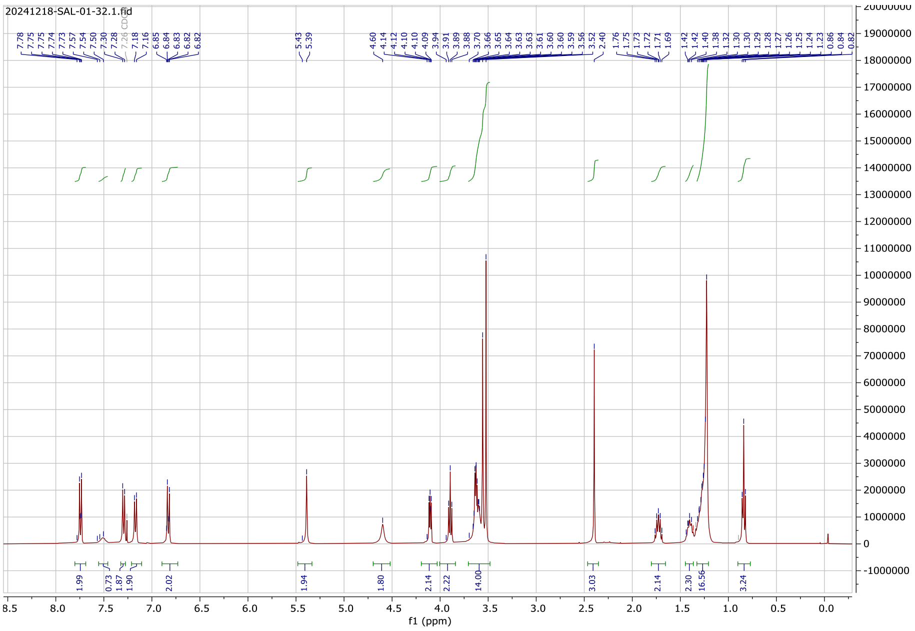


#### $^{1}$H NMR Spectrum

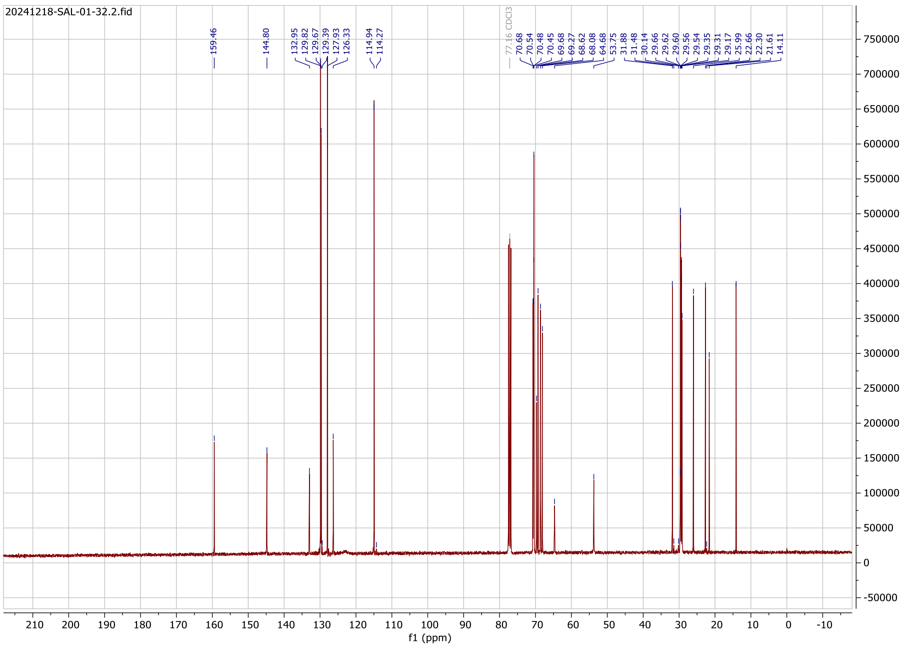

### $^{13}$C NMR Spectrum

### Compound (7b)

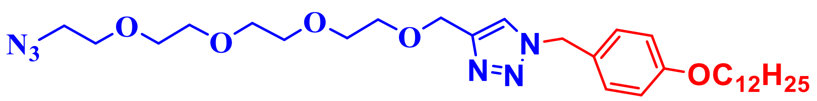


(7b)

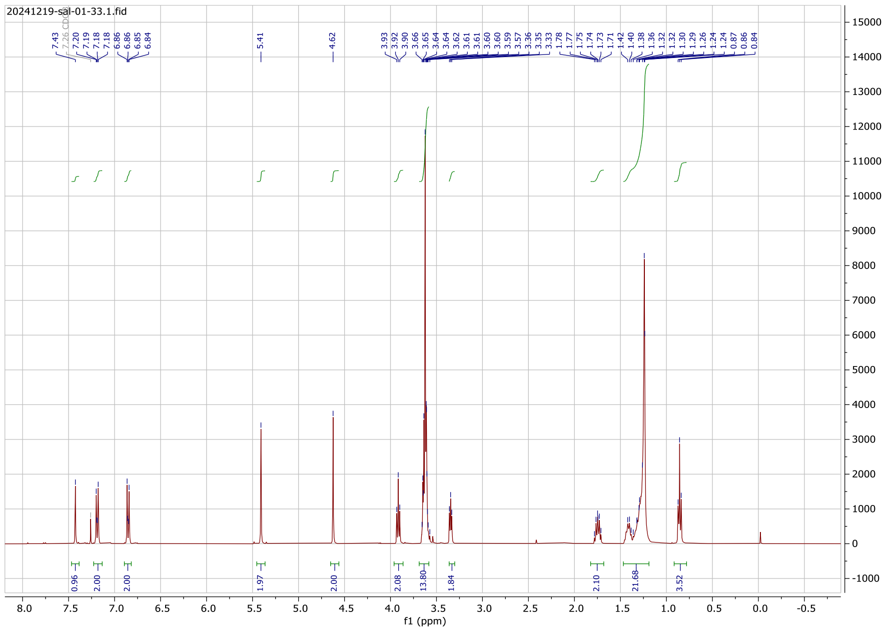


### $^{1}$H NMR Spectrum

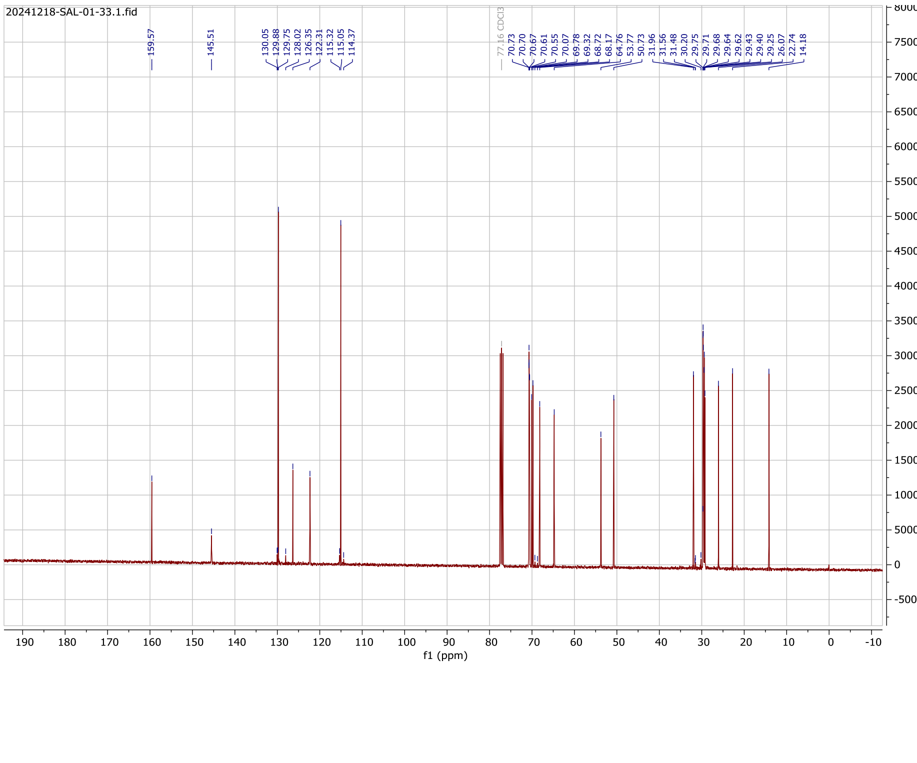

### $^{13}$C NMR Spectrum

### Compound (7c)

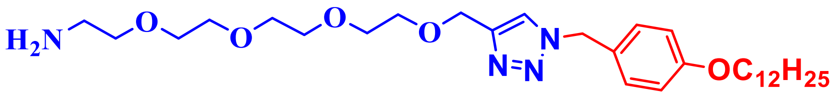


**(7c)**

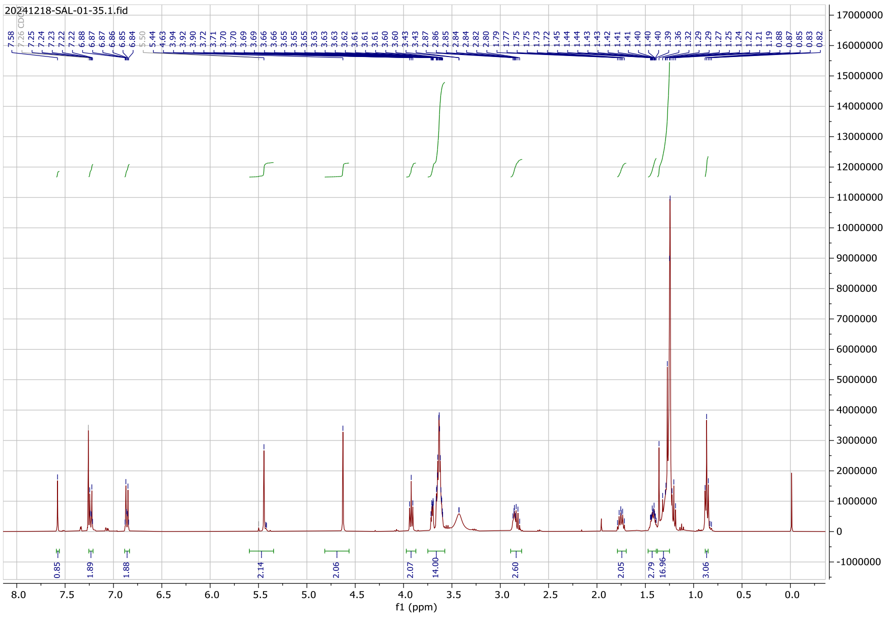


### $^{1}$H NMR Spectrum

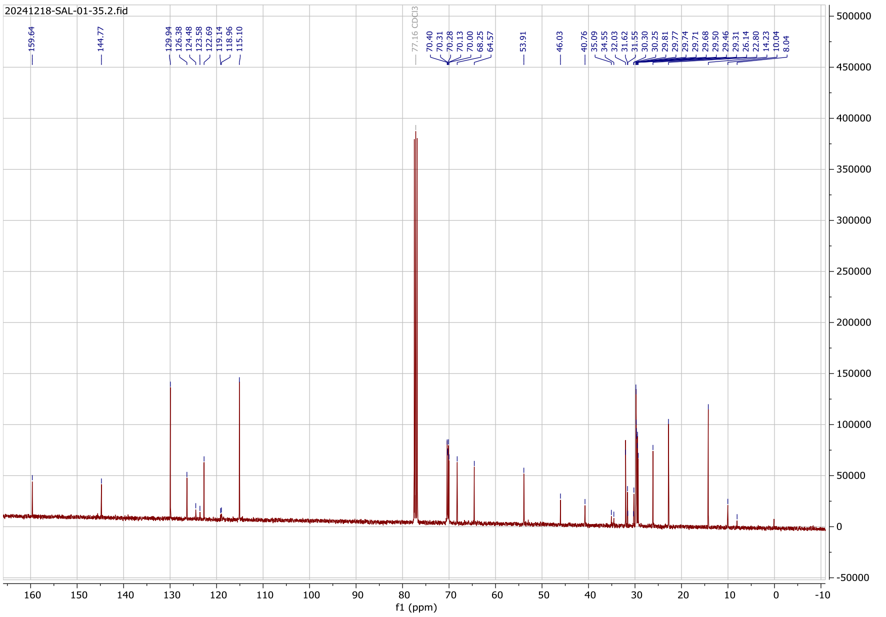


### $^{13}$C NMR Spectrum

#### Compound (7d)

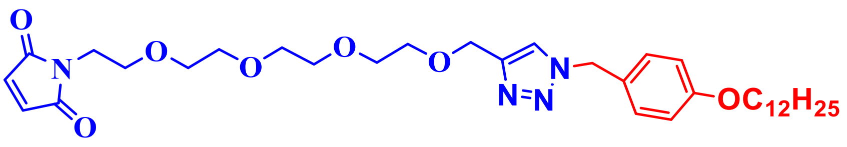


(7d)

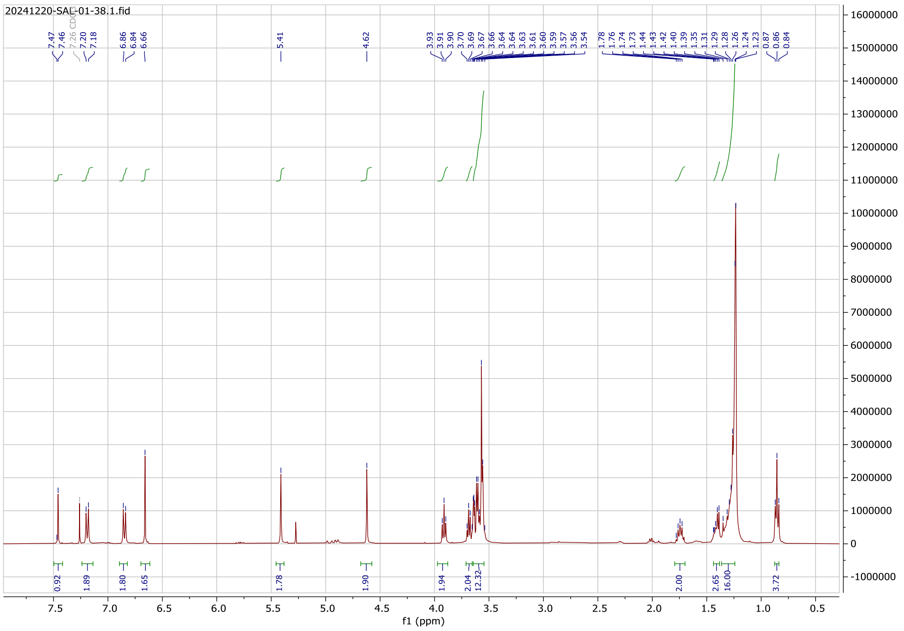

#### ¹H NMR Spectrum

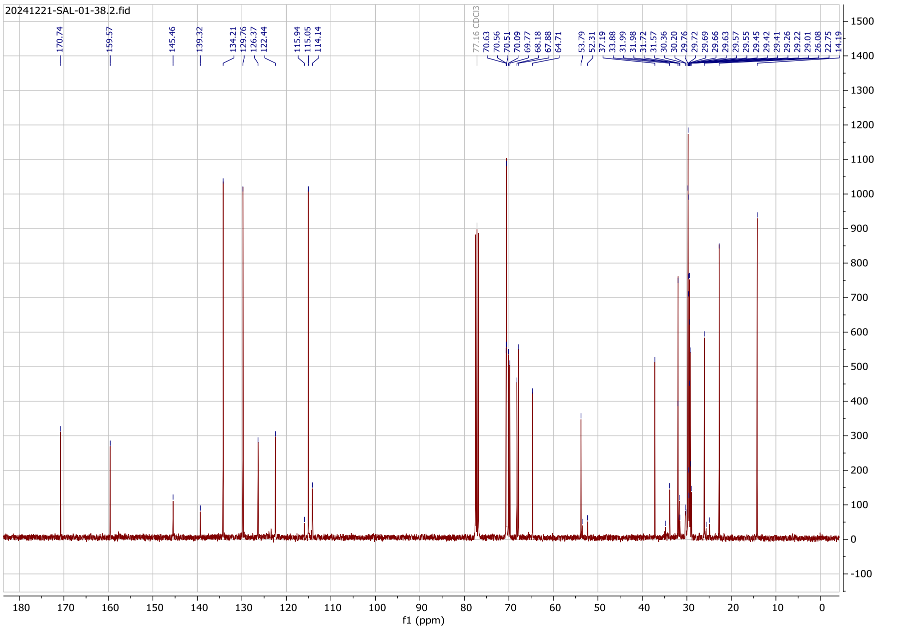

#### $^{13}$C NMR Spectrum

#### Compound (8a)

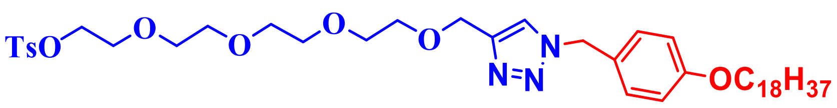


(8a)

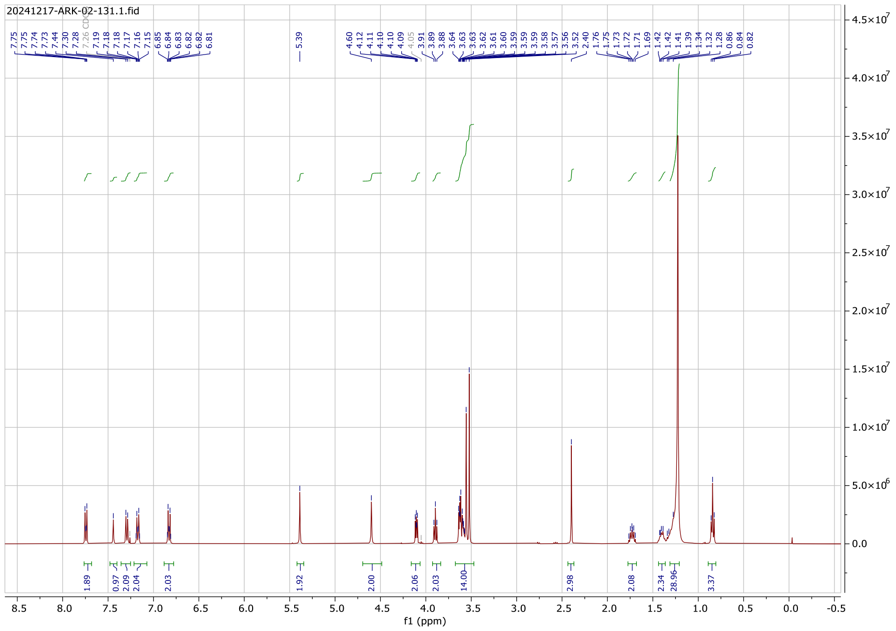


#### $^{1}$H NMR Spectrum

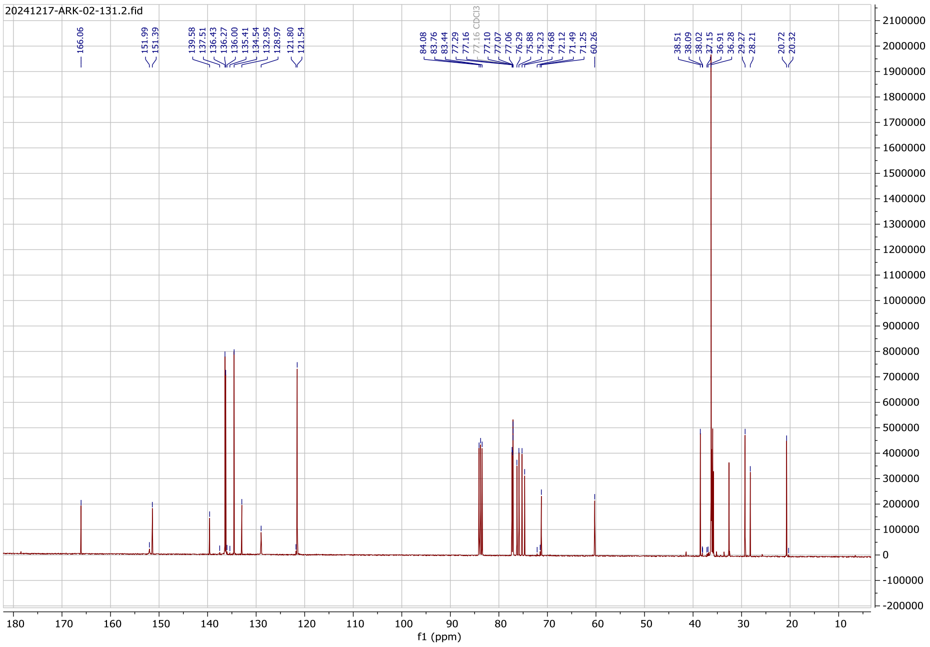

### $^{13}$C NMR Spectrum

### Compound (8b)

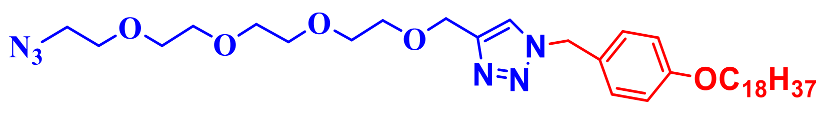


(8b)

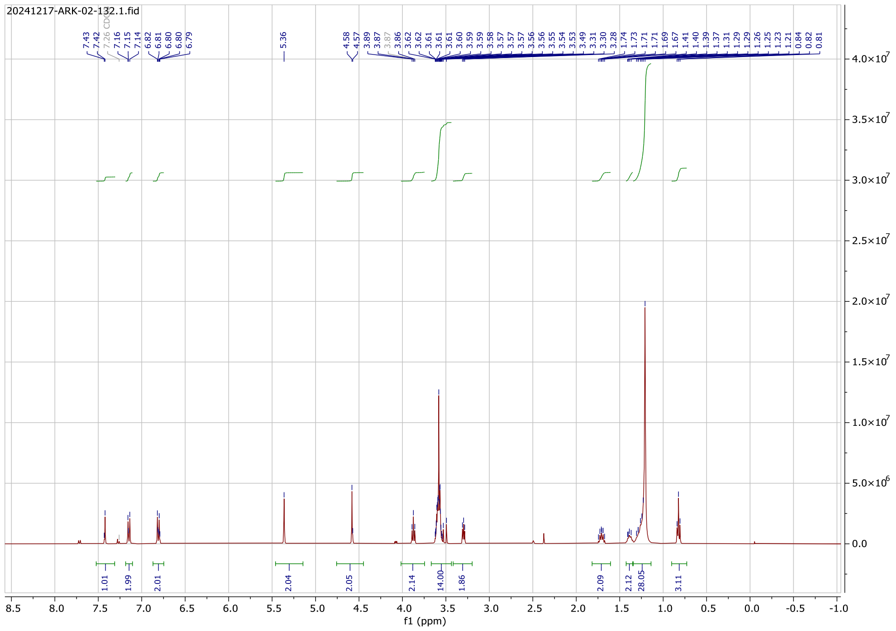


### $^{1}$H NMR Spectrum

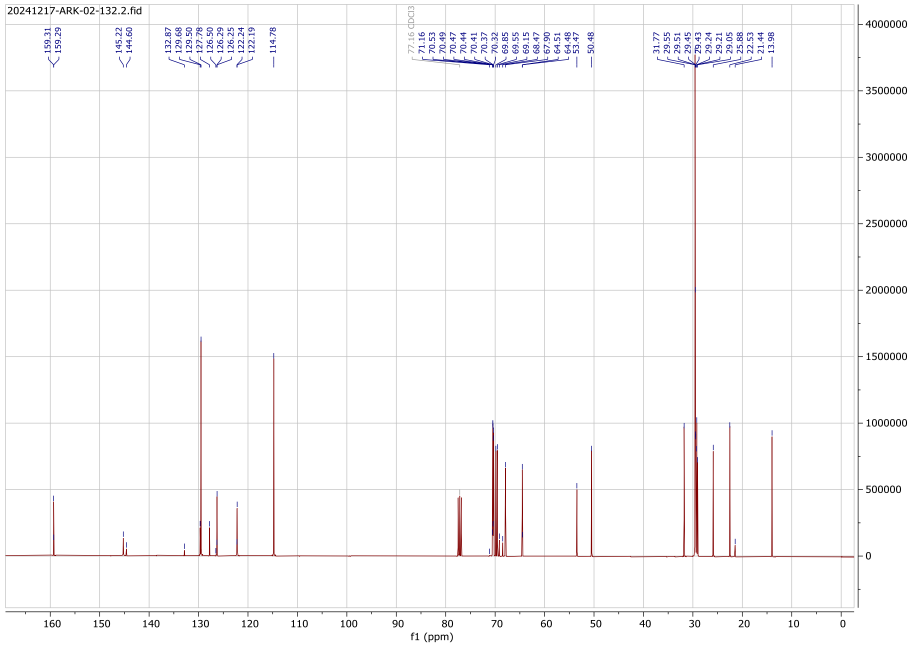

#### $^{13}C$ NMR Spectrum

#### Compound (8c)

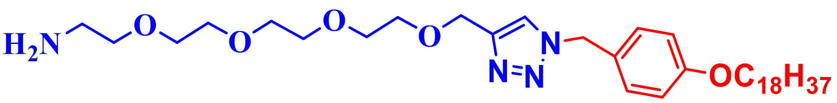

**(8c)**

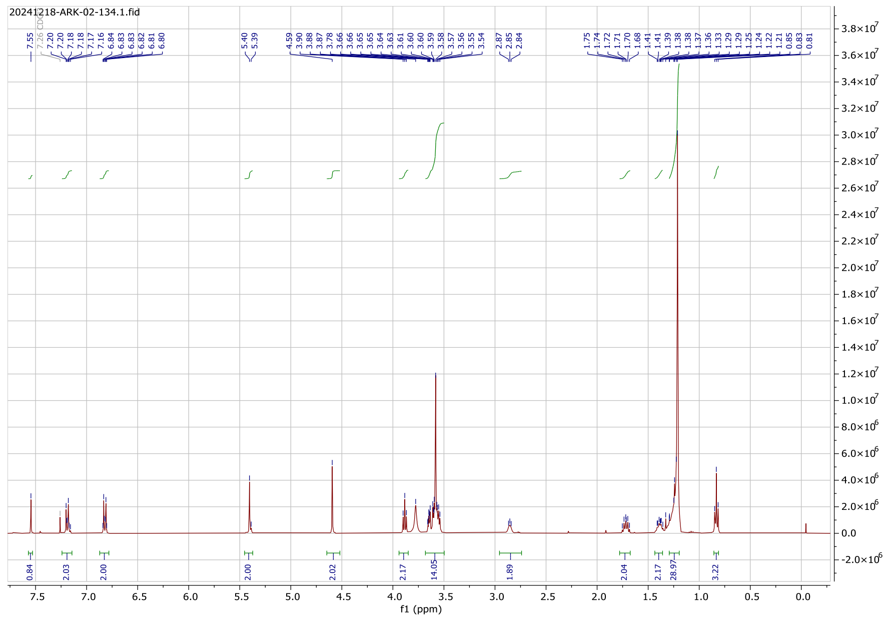

#### $^{1}H$ NMR Spectrum

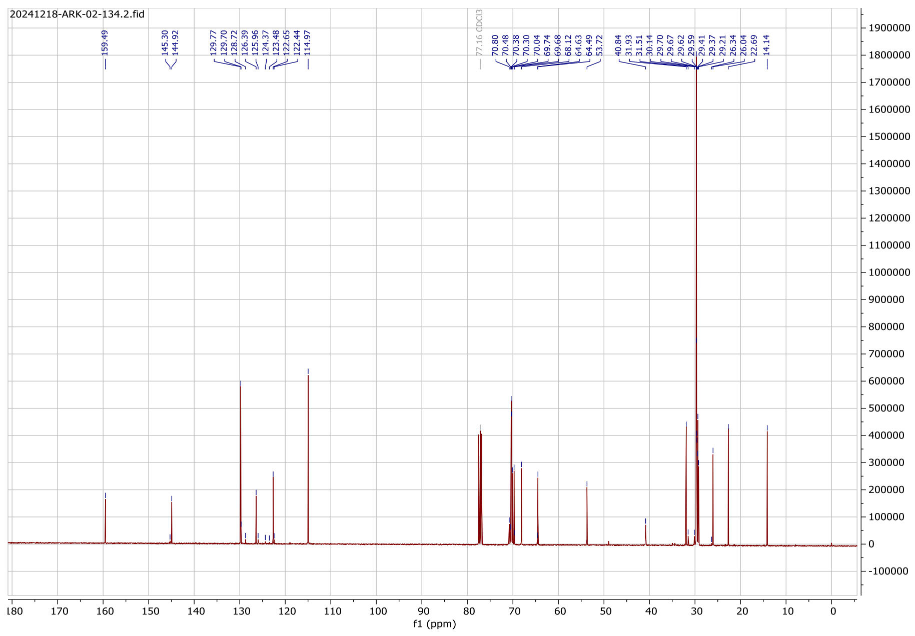

#### $^{13}$C NMR Spectrum

#### Compound (8d)

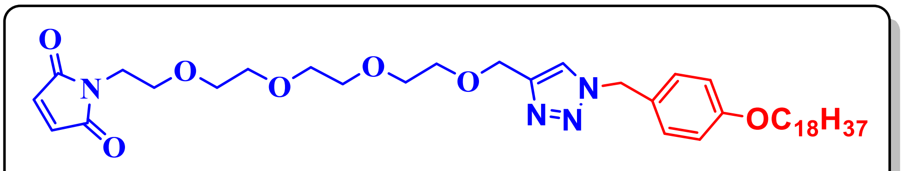

(8d)

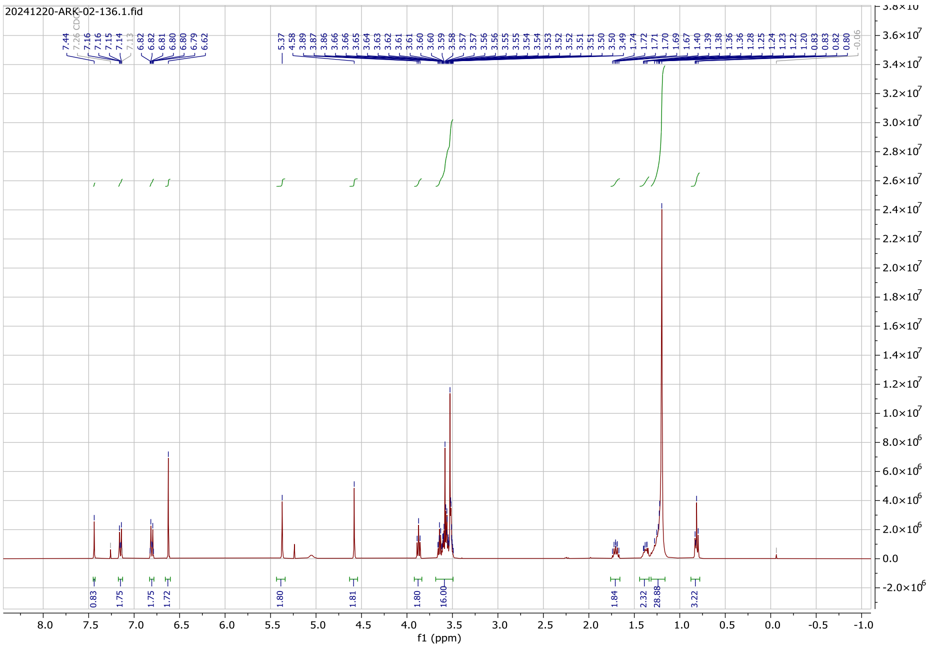

#### $^{1}$H NMR Spectrum

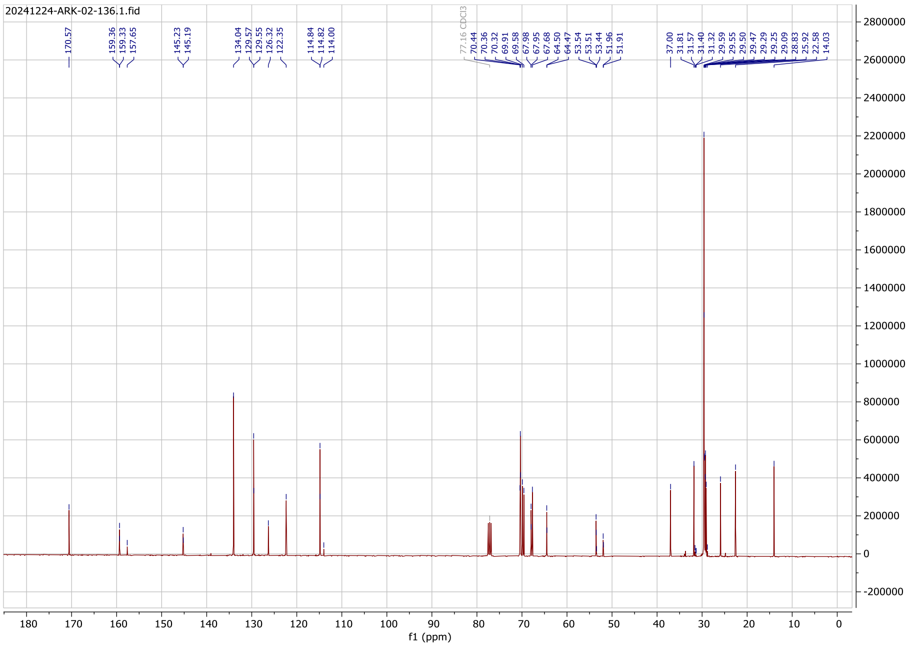

### $^{13}$C NMR Spectrum

#### Compound (11a)

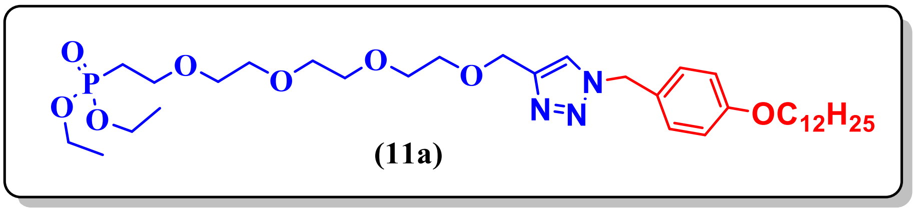


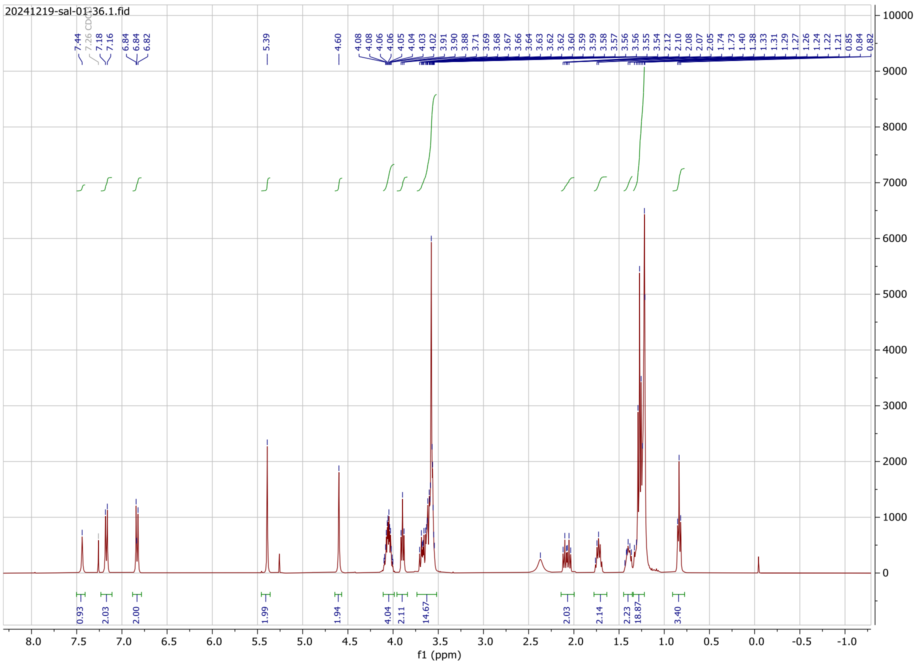

### $^{1}$H NMR Spectrum

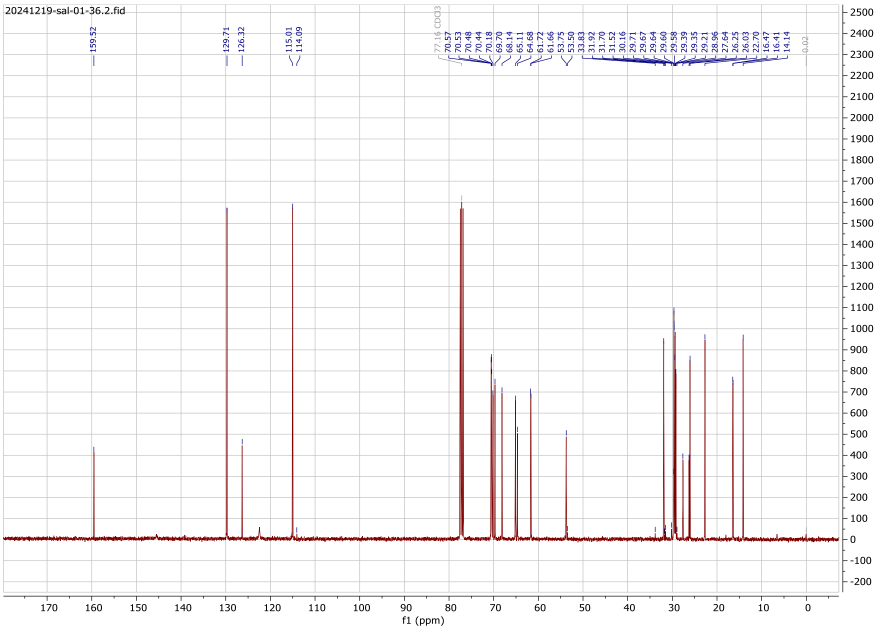

#### $^{13}$C NMR Spectrum

#### Compound (11c)

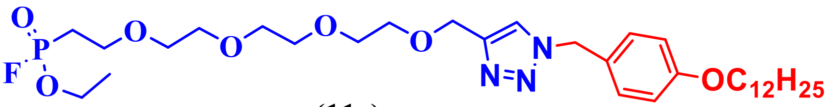


(11c)

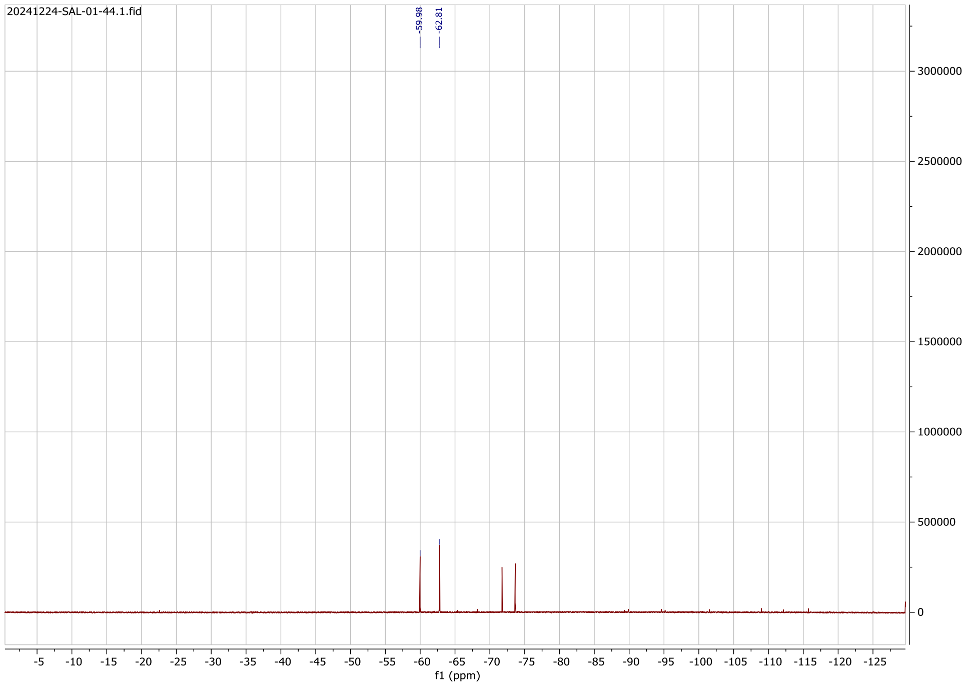


#### $^{19}$F NMR Spectrum

### Compound (1i)

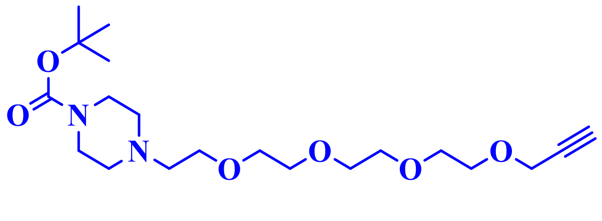

(1i)

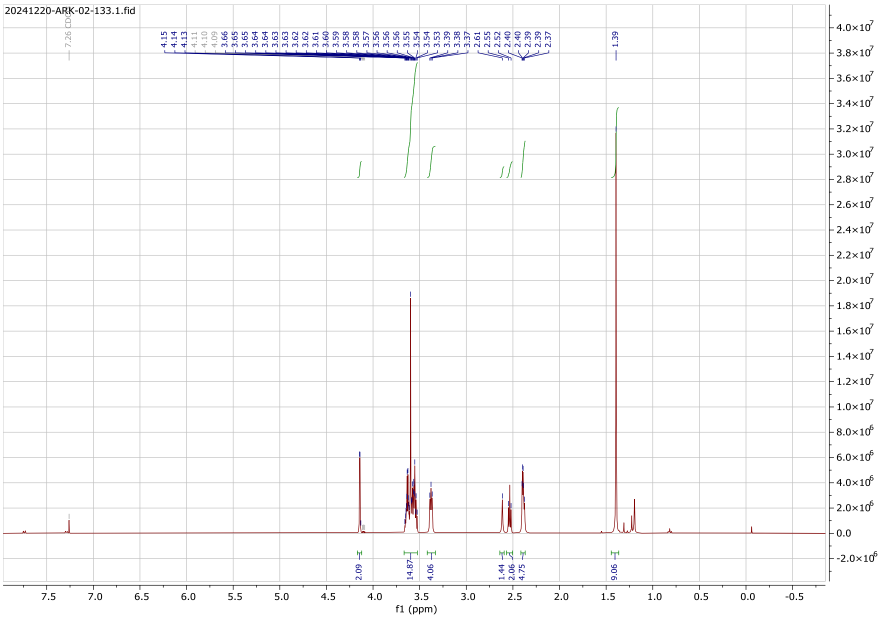


### $^1$H NMR Spectrum

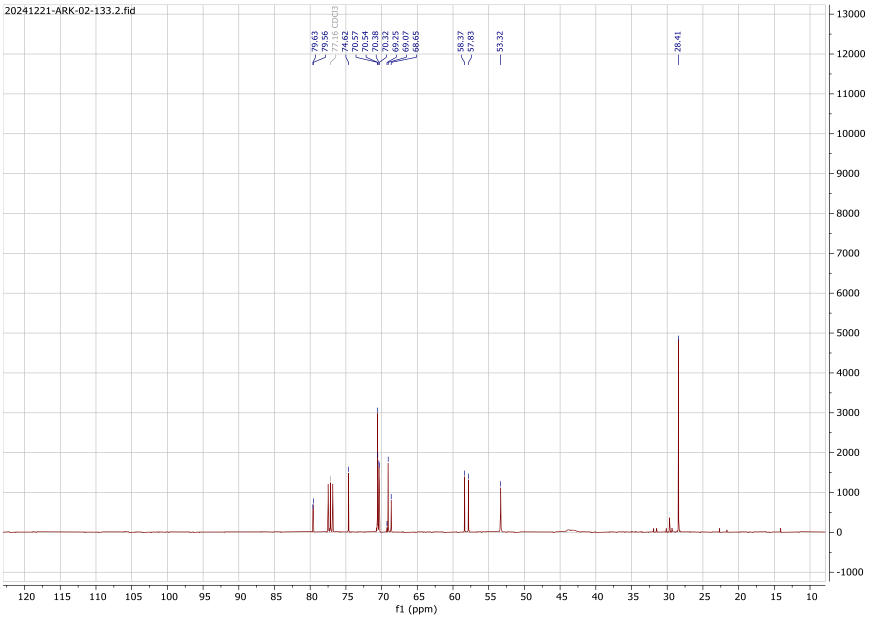

### $^{13}C$ NMR Spectrum

#### Compound (13a)

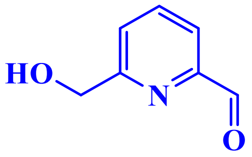

(13a)

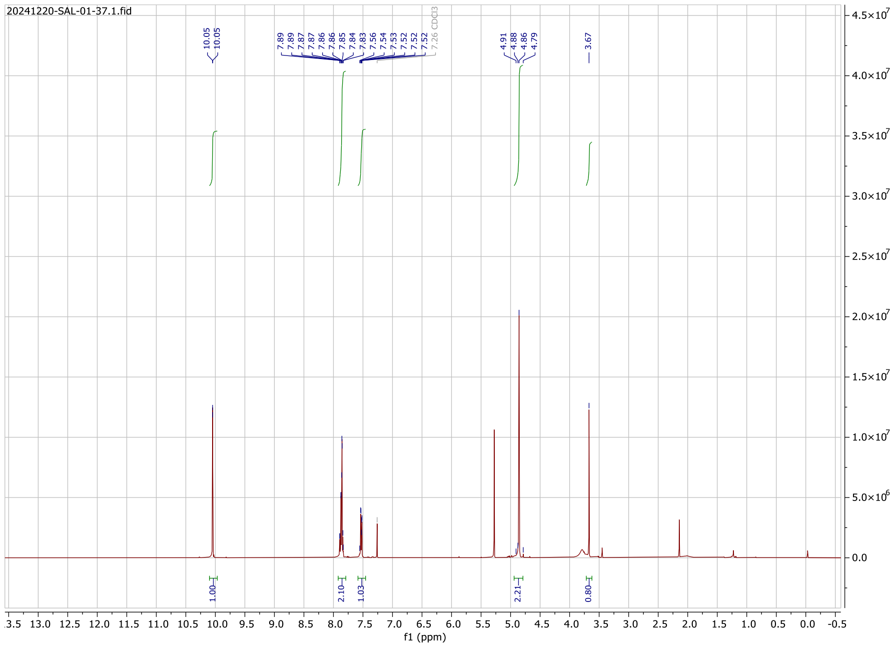

### $^{1}H$ NMR Spectrum

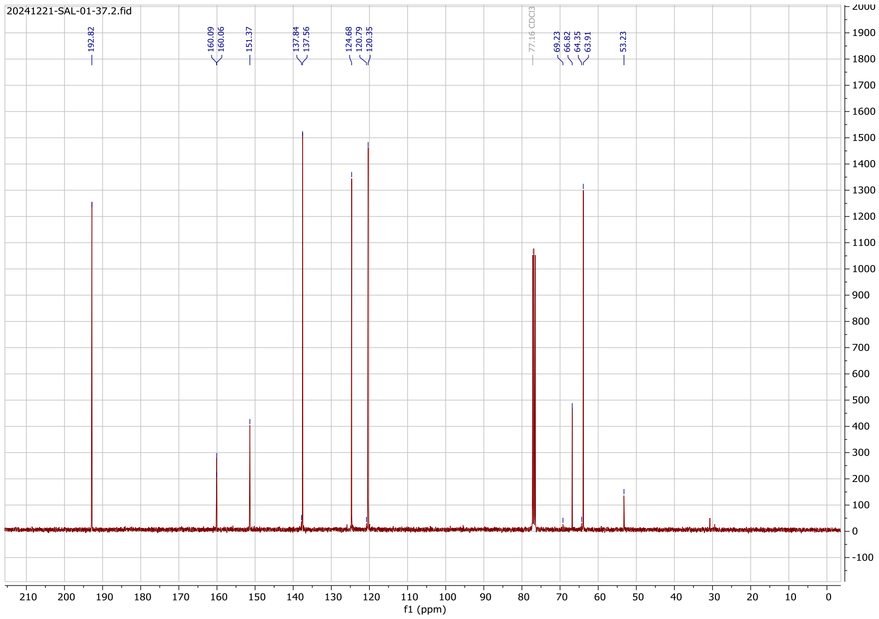

### $^{13}$C NMR Spectrum

#### Compound (13b)

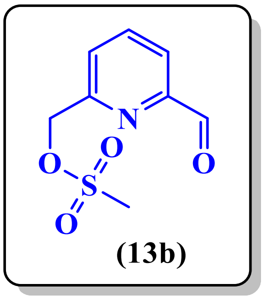


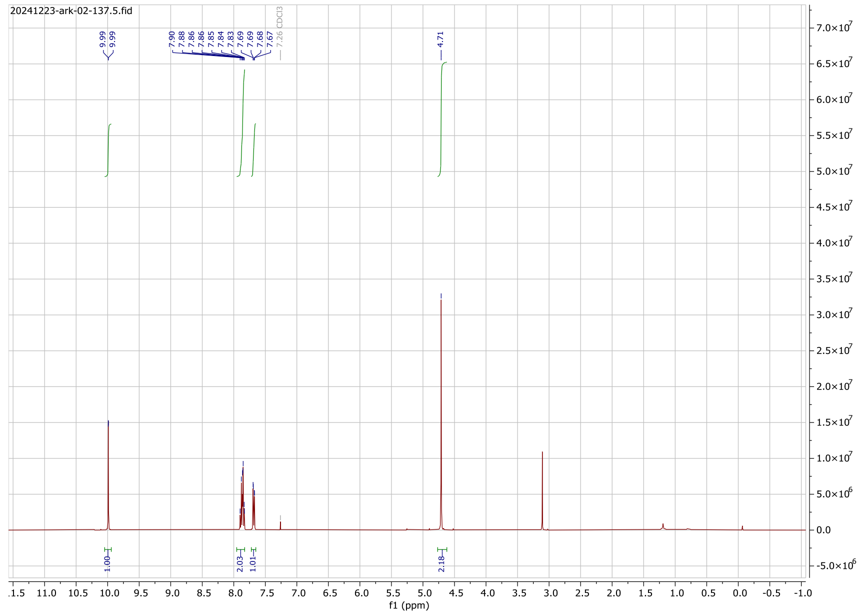


### $^{1}$H NMR Spectrum

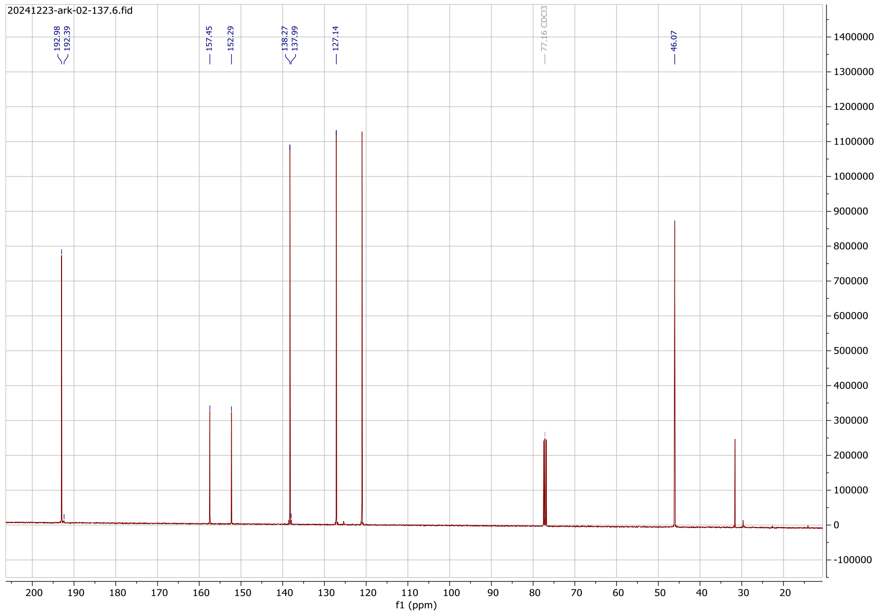

#### $^{13}$C NMR Spectrum

#### Compound (14a)

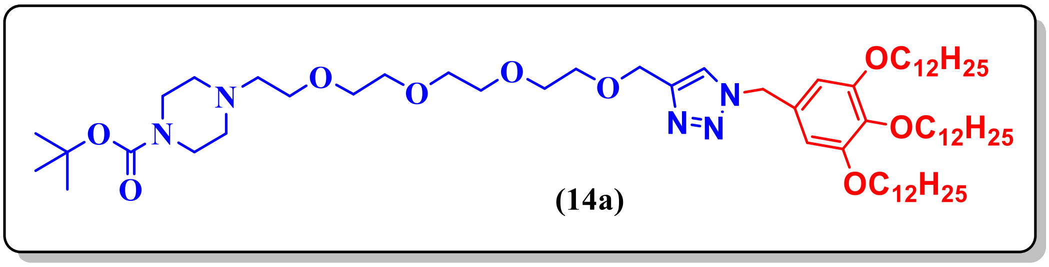


(14a)

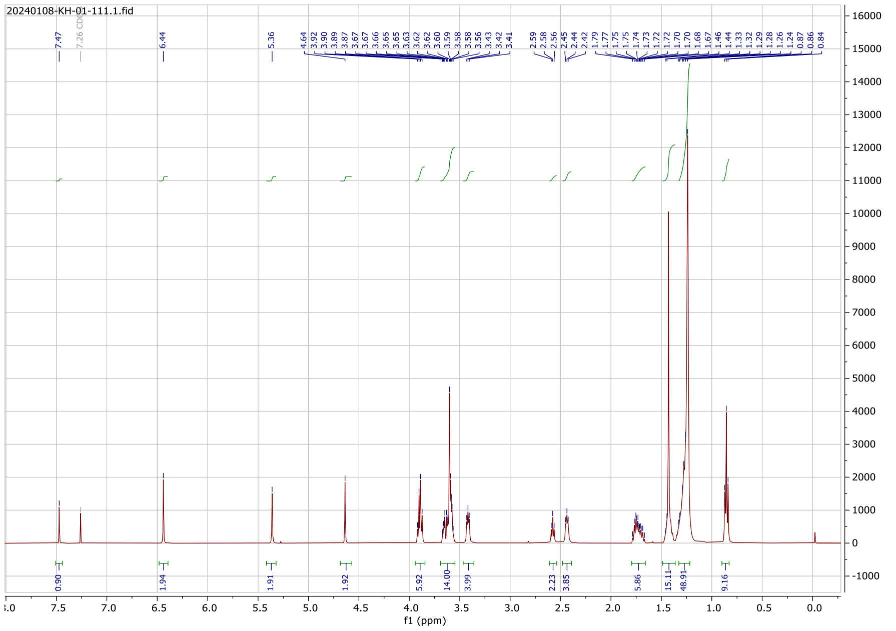


#### $^{1}$H NMR Spectrum

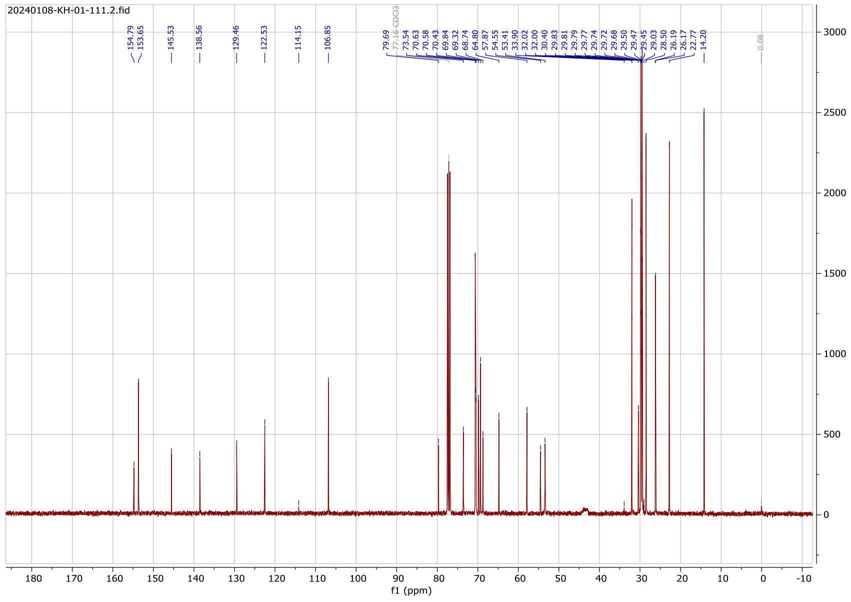

#### $^{13}$C NMR Spectrum

#### Compound (14c)

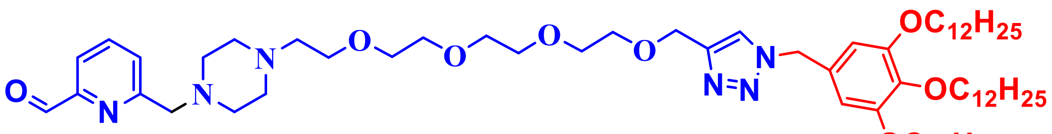


**(14c)**

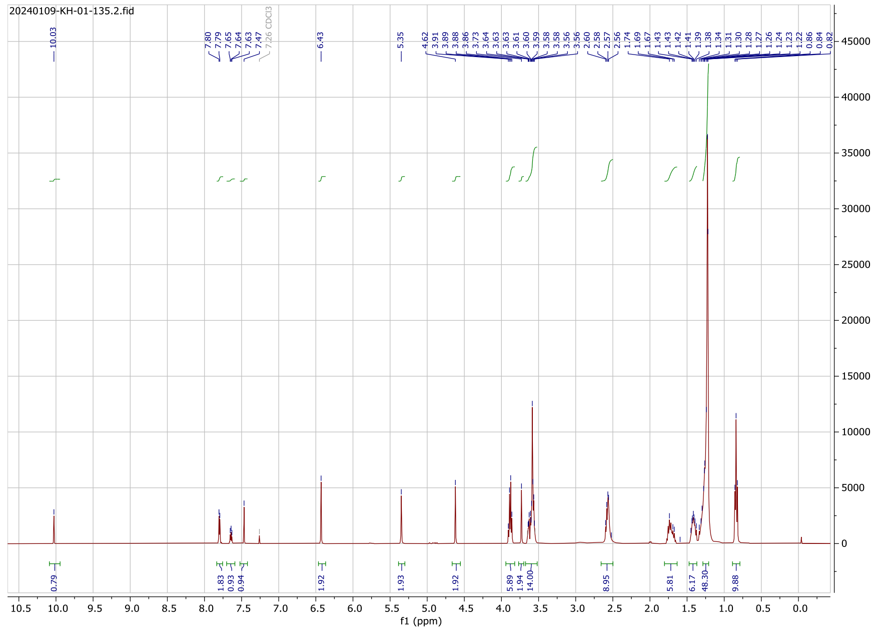


#### $^{1}$H NMR Spectrum

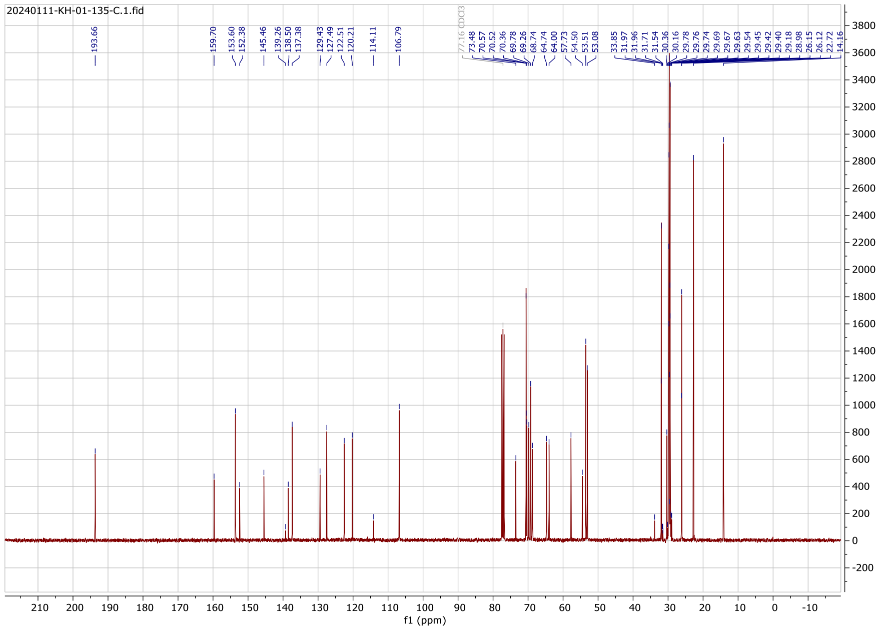

### $^{13}$C NMR Spectrum

### Compound (15a)

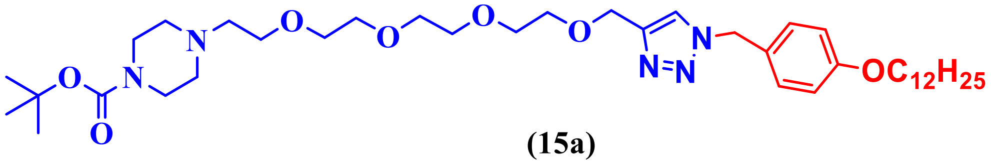


(15a)

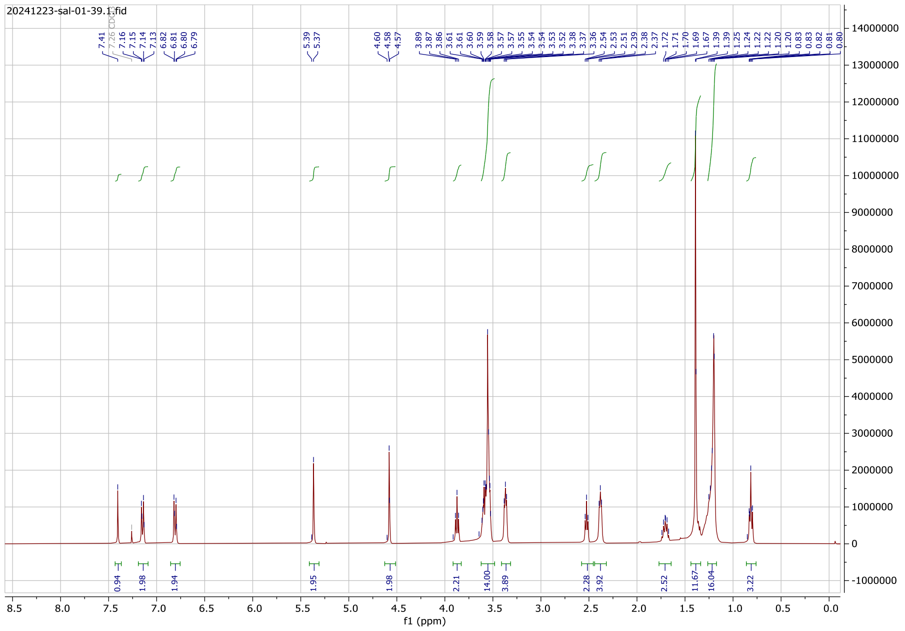


### $^{1}$H NMR Spectrum

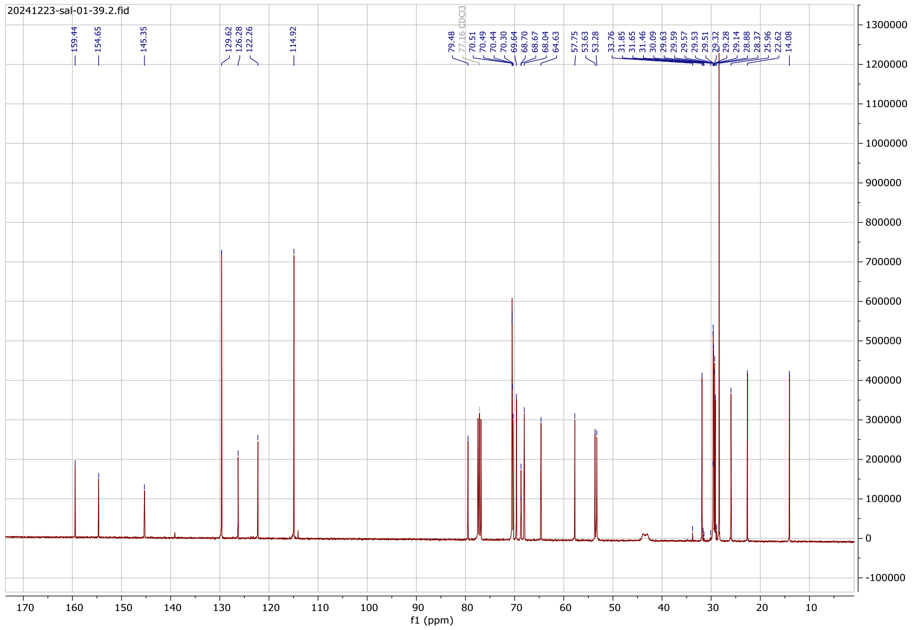

#### $^{13}$C NMR Spectrum

#### Compound (15c)

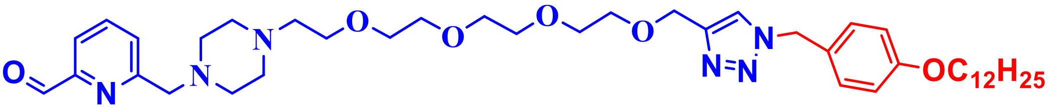


**(15c)**

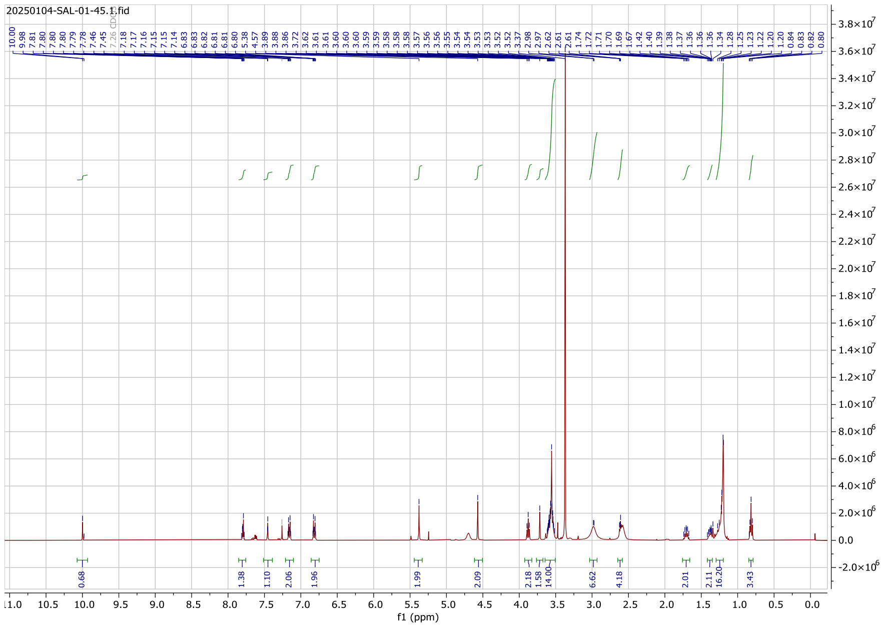


#### $^{1}$H NMR Spectrum

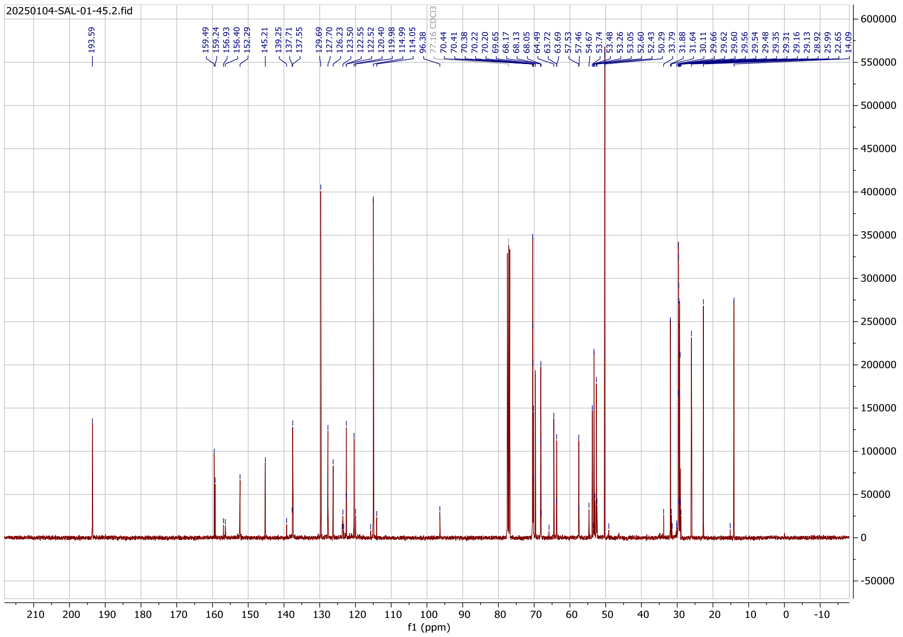

#### $^{13}$C NMR Spectrum

#### Compound (16a)

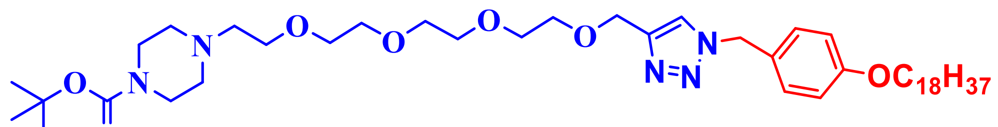


(16a)

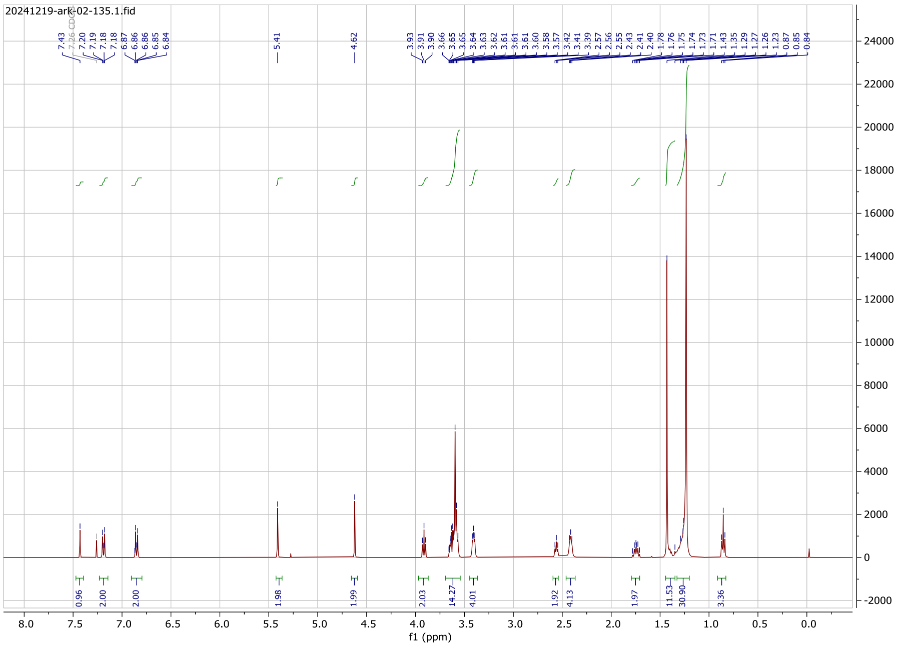


#### $^{1}$H NMR Spectrum

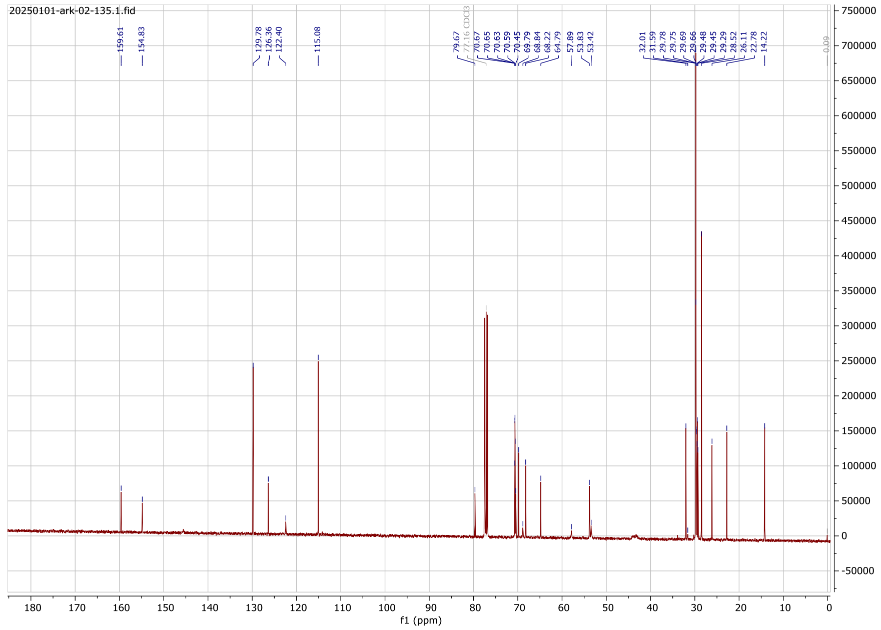

#### $^{13}C$ NMR Spectrum

#### Compound (16c)

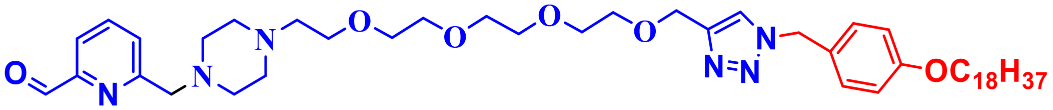


**(16c)**

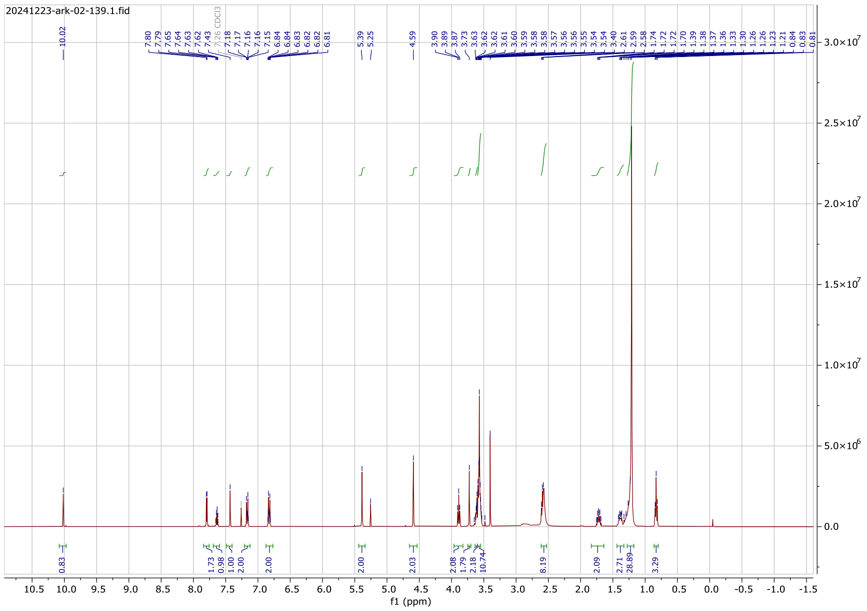


#### $^{1}H$ NMR Spectrum

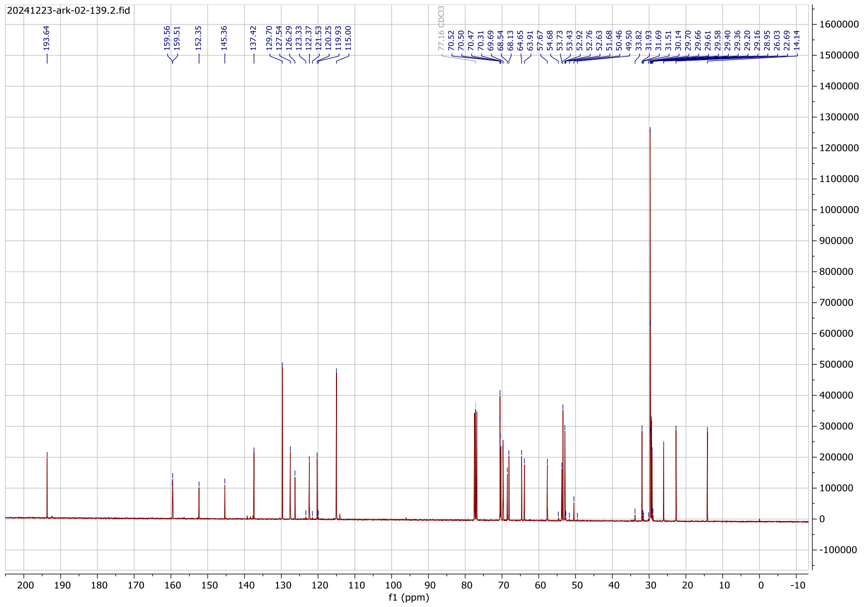

#### $^{13}$C NMR Spectrum
